# A Simulation Study of mRNA Diffusion and Mitochondrial Localization

**DOI:** 10.1101/614883

**Authors:** Paul Horton

## Abstract

**Background:** We have observed a negative correlation between rapid translation initiation of mRNA and their localization at mitochondria. One potential explanation of this anti-correlation is that mRNA which initiate translation away from mitochondria experience a significant drop in mobility and thus remain there. To explore this possibility, we conducted an initial simulation of diffusion and compared those results to gene-specific experimental measurements of mRNA mitochondrial localization.

**Methods:** Here, we conduct a follow-up simulation study to complement the initial one. In particular we attempt a more quantitative analysis, deriving linear scale estimates of mitochondrial localization probability from sequencing based measurements. We compare this data to simulated mitochondrial localization probabilities under a variety of simulation parameter settings.

**Conclusions:** We conclude that if a change in mRNA mobility after translation initiation is a significant factor in explaining the negative correlation between mRNA localization and translation initiation efficiency, then 1) the effective diffusion coefficient of mRNA must be strongly reduced upon translation initiation (e.g. 20x) and 2) mRNA molecules which have not yet initiated translation when approaching a mitochondrial surface must have a non-zero probability of anchoring there.

## 1 Background

Here we describe a follow-up mRNA diffusion simulation study performed to complement an initial study [1]. We have tried to make this document self-contained in terms of reporting what we did and the results. To understand the context and motivation for this simulation the reader is referred to the main study. The initial simulation is also described there. An appendix at the end of this document compares the methods and results of the initial simulation study and this one.

## 2 Dataset

For the follow-up simulation study described here, we used a subset of the datasets used for the main paper. Specifically, we included genes which had a measured value for each of the following features:

- Average Translation Initiation Time [2]
- Average Number of Ribosomes per mRNA molecule [3]
- Om45 ribosome proximity profiling based mRNA mitochondrial localization measurement [5]

which resulted in the 497 genes. After removing two of those which had read counts under 30 in the two relevant experiments of Williams et al. [5], we arrived at the 495 genes listed in file `yeast_genes_mitoLocData.tsv`.

## 3 Estimated mRNA Mitochondrial Localization Probability

For this study we wished to obtain a dataset which could be interpreted directly as the fraction of mRNA molecules for each gene which localize at mitochondria, i.e. the probability that a randomly chosen mRNA molecule of each gene would be localized at a mitochondrial surface at any given time. Will abbreviate this quantity as MLP, the Mitochondrially Localized Proportion of molecules (or equivalently the Mitochondrial Localizing Probability of each molecule), of a given gene.

The next generation sequencing based measurements of Williams et al. [5] seemed the most quantitative; as unlike the light intensities of microarrays which are measured on a log scale, sequencing based technology in principle measures frequency on a linear scale. Williams et al.’s experiments provide a gene-specific measurement of the amount of mRNA molecules engaged with ribosomes (total) and in a separate experiment a gene-specific measurement of the amount of mRNA molecules which are both engaged with ribosomes *and* in the immediate vicinity of a mitochondrial outer membrane (biotin pulldown). Since the former condition includes the later, the ratio of the two measurements should be roughly proportional to the probability that a ribosome-engaged mRNA is localized at a mitochondrial surface. They conducted experiments both with and without addition of the translation inhibitor cycloheximide. Figure 1 shows a scatter plot of the data without addition of cycloheximide which seems a more natural choice for the purposes of modeling *in vivo* conditions.

**Figure 1:**
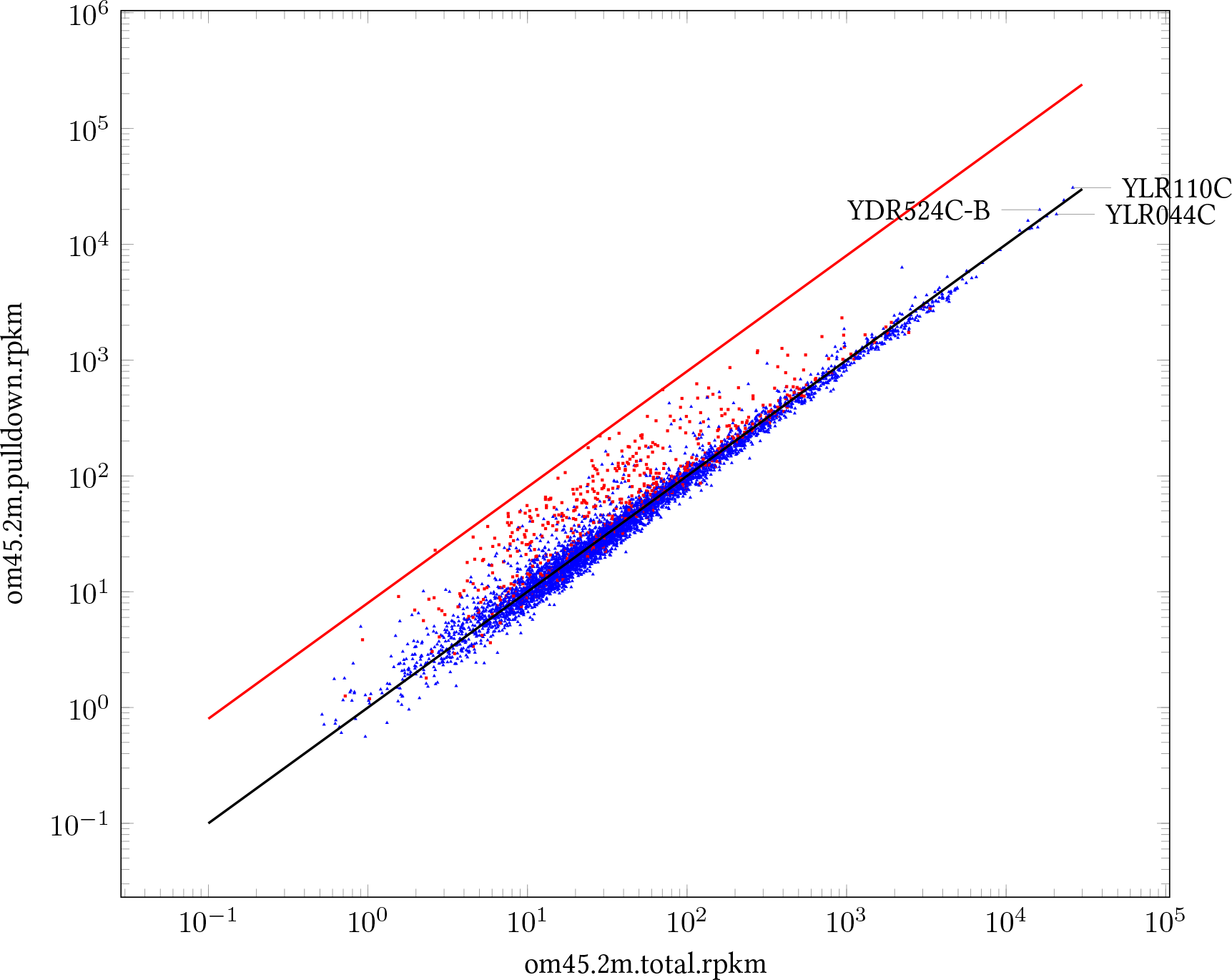
Scatter plot of RPKM measurements [5] of ribosome coverage, total versus biotin pulldown (mitochondrial proximity) for 5201 yeast genes with at least 30 mapped reads under both experiments. The red squares mark the 495 genes with mitochondrially imported protein products used for our simulation. The horizontal axis shows the measurement for total genes and the vertical axis shows the same measurement for ribosomes proximal to mitochondria. The lines *y* = *x* (black) and *y* = 8*x* (red) are shown for reference. Three probably not mitochondrial genes (loci: YLR110C, YLR044C, YDR524C-B) with high pulldown RPKM are labeled.

It is perhaps overly ambitious to derive probability from data which is usually only interpreted as fold-change. However we feel it is a worthy goal, as the biological consequence of a change in mitochondrial localization from say 1% to 2%, could easy differ from the consequence of a change from say 50% to 100%, even though those are both 2-fold changes. And it makes sense in our context, because the results of our computer simulation can be naturally be compared to measured mitochondrial localization probabilities.

The remaining question is what the proportionality constant should be. This requires an additional assumption. The one we chose to make is that some genes are translated nearly exclusively at the surface of mitochondria. If one accepts this assumption, then the proportionality can be adjusted so that the highest probability is close to 1. To see what might be a reasonable value for that constant, we considered the distribution of the ratio of RPKM (Read Per Kilobase Per Million Reads Mapped) listed in their supplementary material as `om45.2m.pulldown.rpkm` and `om45.2m.total.rpkm` for the 5201 genes with at least 30 reads mapped both conditions (columns `om45.2m.pulldown.cts and om45.2m.total.cts`) in their supplemental material. Figure 2 shows the distribution of the `pulldown` to `total` RPKM ratio. The four genes with the highest ratios were YMR207C (8.65), YBR084W (7.89), YOR211C (7.77) and YMR302C (7.30). As expected, all of these genes encode mitochondrially imported proteins and also exhibit relatively high mitochondrial localization in microarray based measurements as well. For example, in the microarray measurements of Sylvestre et al. [4] reporting scores with mean and standard deviation of 0.083 ± 0.083 (median= 0.82), the scores of YMR207C, YBR084W, YOR211C, YMR302C are 0.104, 0.288, 0.199, and 0.168 respectively. In accordance with our assumption that the localization probability of the genes which most strongly localize to mitochondria should be close to 100%, we initially thought it would be reasonable to estimate the MLP of genes with RPKM ratios of 8 or more to be 100% as in this initial formula:

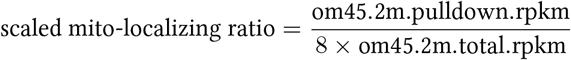

**Figure 2:**
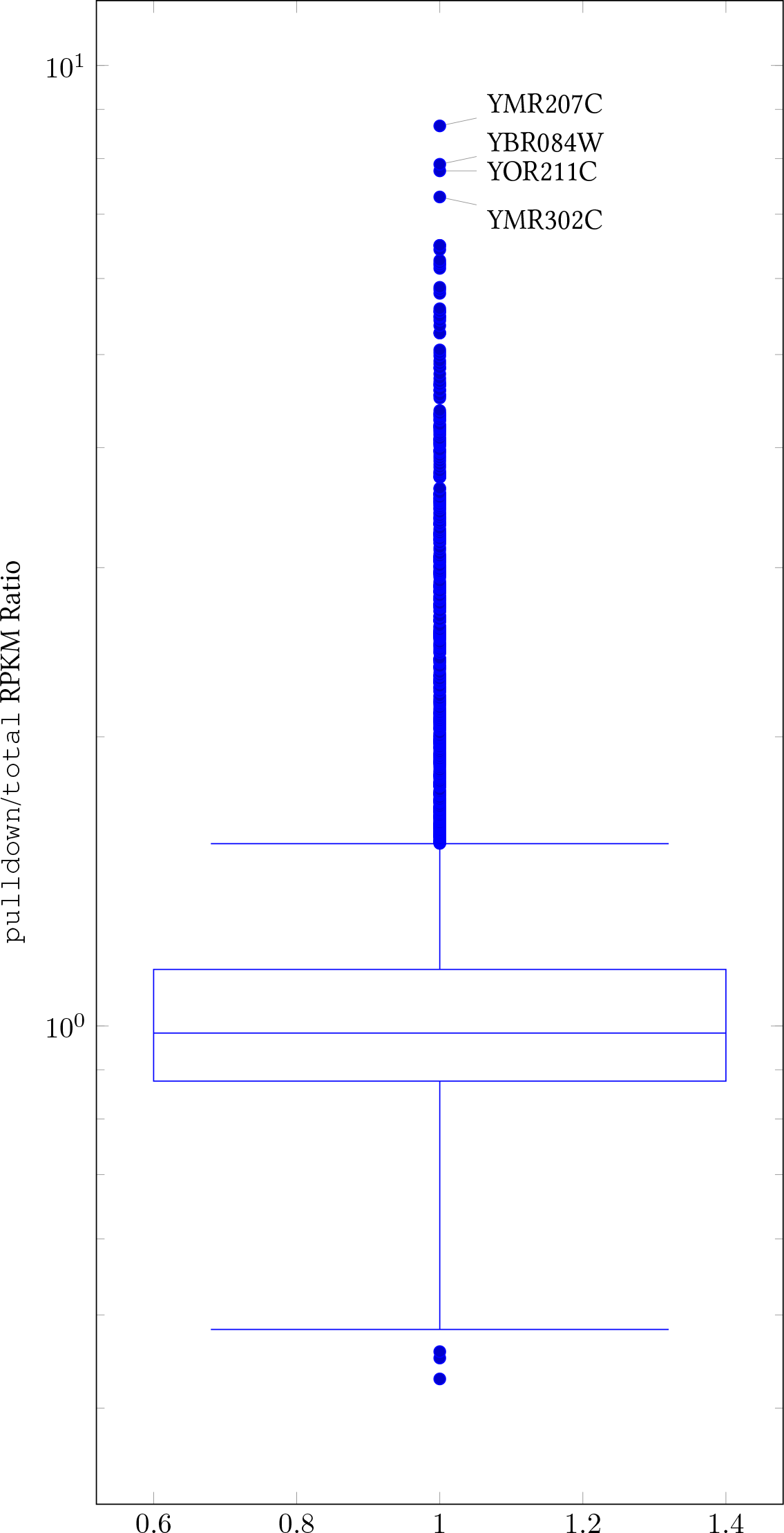
The distribution of the ratio of total versus biotin pulldown RPKM [5] is shown. The genes with the highest ratios are labeled with their locus names.

(Except we used a probability of 1.0 for YMR207C, the only gene with a ratio higher than 8.)

### Background Adjusted Mitochondrial Localization Probability

Unfortunately, simply scaling the RPKM ratio leads to a distribution in which most genes encoding non-mitochondrially imported proteins have a 5–15% MLP (figure 3; top half, upward bars), which is higher than the mode of the probability assigned to many mRNAs encoding mitochondrially imported proteins (figure 3; top half, downward bars)!

**Figure 3:**
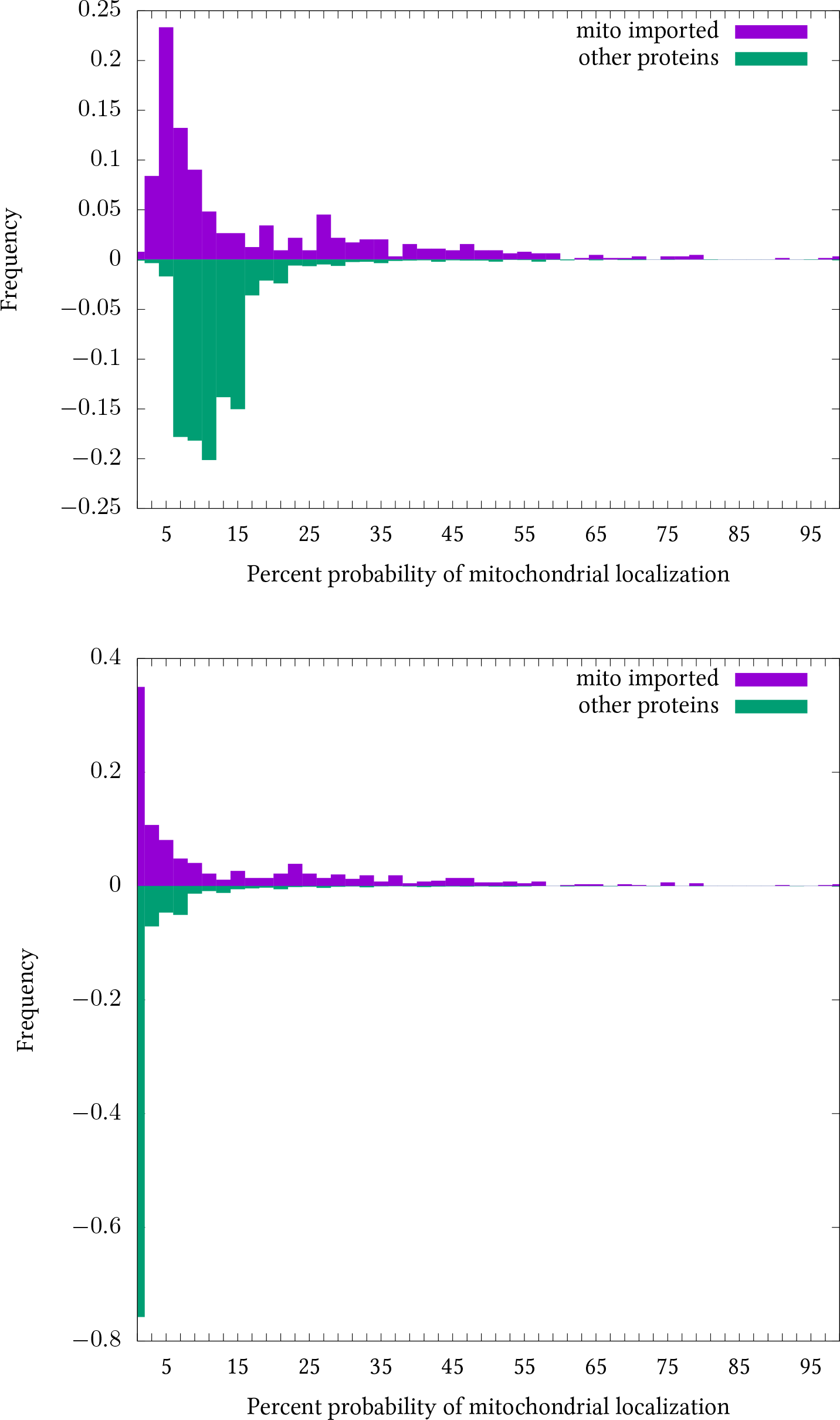
Bidirectional histograms showing the frequency of genes with given mitochondrial localization probabilities for each bin of width 2%. Frequencies for proteins not imported into mitochondria are shown as negative numbers. Top) probabilities estimated by scaled ratio of RPKMs. Bottom) background adjusted probabilities.

However, looking at the black diagonal line in the scatter plot of figure 1, we noticed that (when viewed on a log scale) the pulldown RPKM for most genes is roughly proportional to their total RPKM, even though most of those genes are unrelated to mitochondria. Indeed the top five pulldown RPKM genes are well annotated and three of them (YLR110C, YLR044C and YDR524C-B) do not appear to have a special connection to mitochondria. We speculate that most of the genes with an RPKM ratio near one do not specifically localize to mitochondria, but rather there is some non-specific background component included in the pulldown RPKM. For example, it could be the case that the 2 minutes in which the cells are flushed with biotin is sufficient time for some mobile mRNA-ribosome complexes to have chance encounters with the BirA biotin ligase anchored to the mitochondrial outer membrane, or perhaps some technical factor in the experiment is involved. Therefore we used the following adjusted formula to compute MLP:

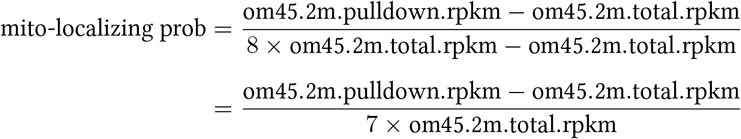

(Except values which would be less than zero by this formula were set to 0.0, and those which would exceed one were sent to 1.0) After this adjustment for background, the estimated MLP of the genes not encoding for mitochondrially imported proteins is drastically reduced while the MLP of genes encoding for mitochondrially imported proteins is reduced only slightly (figure 3; bottom). As a result the separation is much better, with 32% of genes encoding mitochondrially imported proteins being assigned an MLP > 0.14, but only 2% of other genes exceeding that threshold. This is especially impressive given that some of those 2% may turn out to in fact be mitochondrially imported, but simply not yet annotated as such. The successful separation between mitochondrially important proteins and other proteins notwithstanding, overall our estimated MLPs still seems too low for the genes encoding mitochondrially imported proteins. Nevertheless, we adopted this MLP.

Figure 4 shows the adjusted MLP values versus average translation initiation time for genes encoding mitochondrially imported proteins. There is much scatter in the plot and many of the genes show nearly zero MLP. Nevertheless one can discern a tendency for some genes with slow translation initiation to have a relatively high MLP. One can also see that, as expected, genes with fast translation initiation tend to have more ribosomes. There also seems to be some connection with MLP, in that the quickly initiating genes with high MLPs tend to have many ribosomes. However in the simulations reported in this manuscript we did not further explore the role of the number of ribosomes.

**Figure 4:**
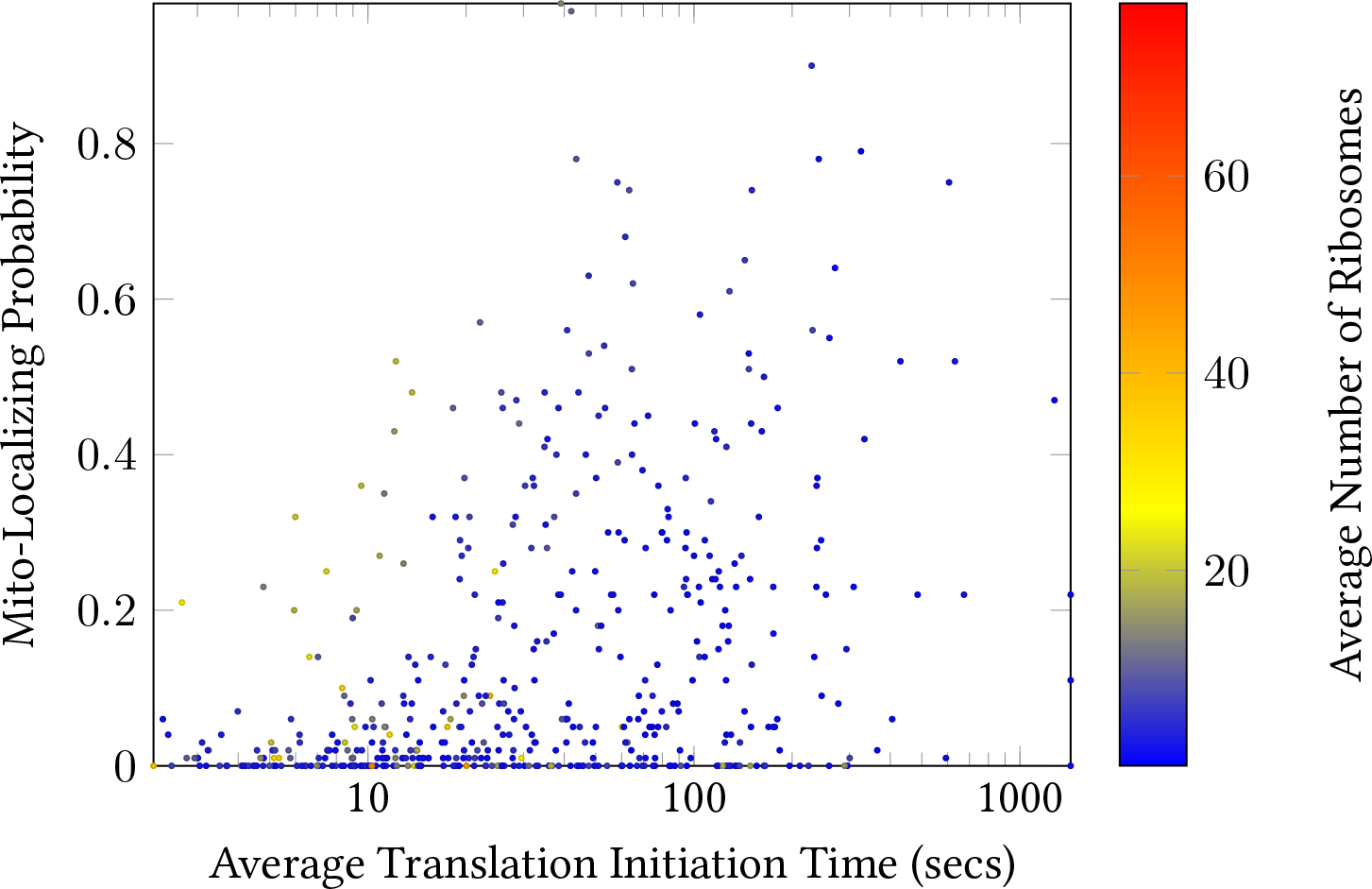
Scatter plot of the RPKM ratio based mitochondrial localizing probability and average translation initiation time for 495 yeast whose protein products are imported into mitochondria. Points are colored according to their average number of ribosomes.

## 4 Simulation

We implemented a discrete time random walk simulation of an mRNA molecule diffusing through the cytosol; and possibly anchoring to a mitochondria. As detailed in section 4, in each time step the molecule moves a small distance in a randomized way. The simplified architecture of the cell used in the simulator is shown in figure 5. Mitochondria are placed randomly in the spherical shell with an inner radius of 1*µ*m and outer radius of 2.5*µ*m, in such a manner that they do not overlap the nucleus or each other. The mRNA diffusion starts at the surface of the cell nucleus and continues until the mRNA has anchored to a mitochondrion or 11 minutes (a typical half-life of yeast mRNA) has passed. The speed of diffusion changes upon initiating translation. At the start the mRNA has not yet initiated translation, but at each step it may do so with probability equal to the length of a time step divided by the average translation initiation time (as measured by Shah et al. [2]) of the gene it represents. For a given set of parameters, the simulation is repeated many times and the mitochondrial localization probability is estimated as the fraction of runs for which the mRNA anchored to a mitochondria. At a conceptual level, the simulation has four parameters which we tried varying: the effective diffusion coefficients of mRNA before and after initiating translation, and the mitochondrial anchoring probability before and after initiating translation. Additional technical parameters describe the details of how the particle moves (section 4).

**Figure 5:**
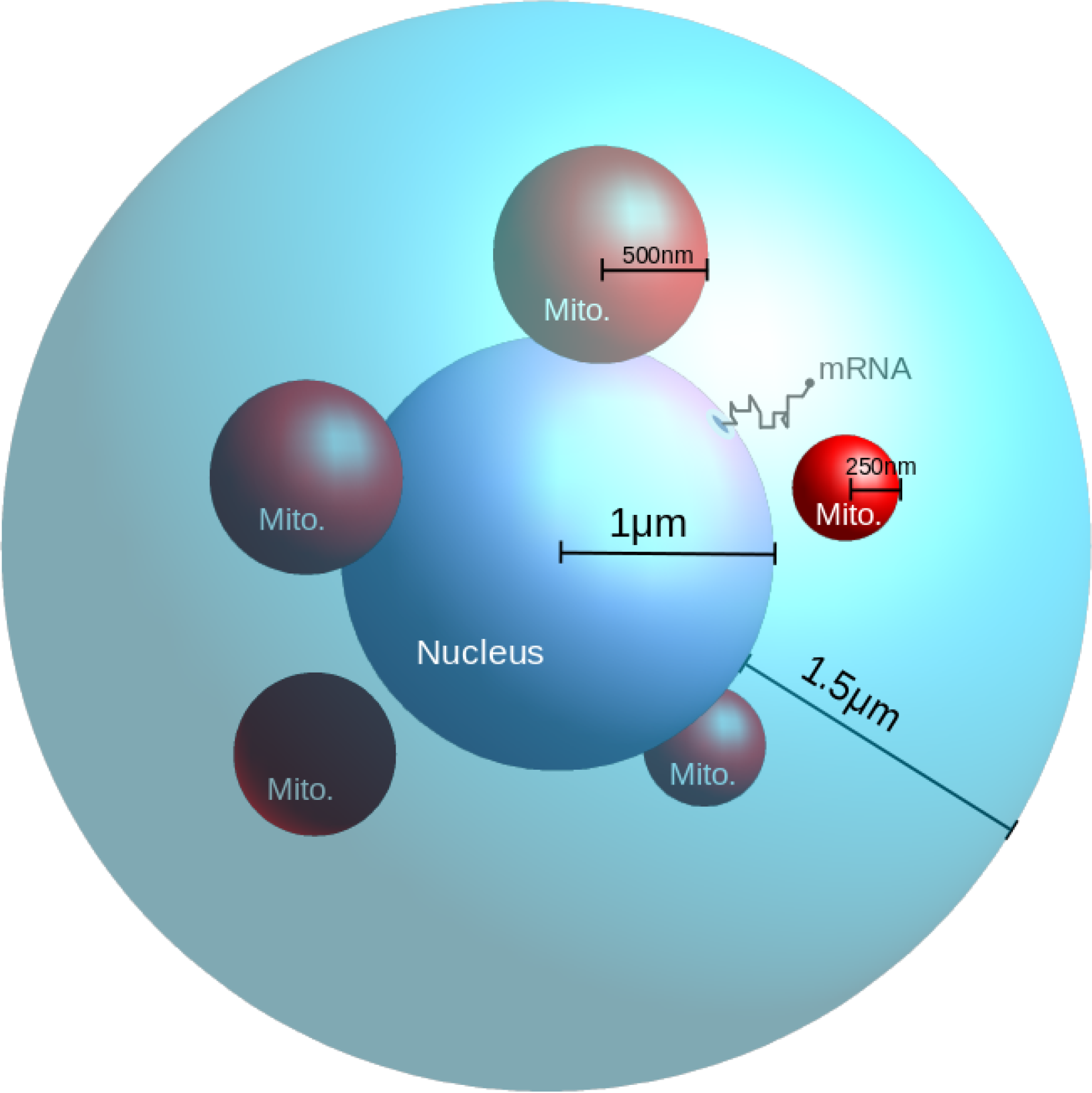
Schematic depiction of the mRNA diffusion simulation. All objects are spherical. The cell has a radius of 2.5*µ*m, the nucleus 1*µ*m, and the 5 mitochondria have radii of 250 to 500nm. The mRNA is a point which undergoes a 3D random walk in the cytosol, starting from the surface of the nucleus.

### Operations done at each simulation step

The following operations are done at each time step of the simulation. If the mRNA has not yet initiated translation, it is given a chance to do so with probability inversely proportional to its gene-specific average translation initiation time, and if it does indeed initiate translation its diffusion coefficient and anchoring probabilities are updated appropriately. Then the mRNA attempts to move one step. If the attempted step does not cause the mRNA to collide with anything, the move is accepted and the simulation goes to the next time step. If the attempted move would have placed the mRNA inside of the nucleus or outside of the cell the tentative move is rejected and the mRNA stays put for that time step. If the attempted move would have placed the mRNA inside of a mitochondria; the mRNA is given a chance to anchor to it with probability equal to its current anchoring probability. If the mRNA anchors that event is recorded and the run is terminated. On the other hand, if the mRNA does not anchor the tentative move is rejected and the mRNA stays put for that time step.

### Interpretation of Diffusion Coefficient

In this follow-up study the mobility of mRNA molecules is summarized with a single “effective diffusion coefficient”. By “effective” we mean that it represents the average mobility of the mRNA, even if in reality the mRNA is only freely diffusing for some of the time (with a higher diffusion coefficient) and has limited mobility the rest of the time, perhaps exhibiting stationary periods or corralled diffusion.

### Relevant Gene Features

The simulation is essentially a monotonic mapping from a gene’s average translation initiation value to its mRNA mitochondrial localization probability. This is so because in our simulation the diffusion coefficient and sometimes the anchor probability differs before and after translation initiation. (The software has been implemented to allow the anchor probability after translation initiation to be a function of the average number of ribosomes for that genes mRNA, but we did not explore that.)

### Detailed Movement of Particle

#### Movement Per Time Step

To model diffusion, it is necessary that the particle exhibits what we call Gaussian displacement over time spans long enough to be of interest. This constraint dictates that if the simulation were run repeatedly for a sufficient time span the net movement (displacement) of the particle in any direction should approximate a normal distribution with mean zero and variance equal to 2*Dt* (twice the product of the length of time and the diffusion coefficient). This constraint allows for some variation in the technical details of how the particle moves in each step. Our software currently provides the option of moving the particle in each step in one of three ways:

- `gauss`: Variable distance movement along all three axes; those distances independently drawn from a normal distribution with mean zero and variance 2*t*/*D*, i.e. *N* (0, 2*t*/*D*).
- `fixed`: Fixed distance movement along all three axes; those distances independently randomly selected from 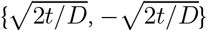.
- `fixed1`: Fixed distance along one randomly chosen axis; the distance randomly selected from 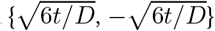.

(Historically, we first implemented `fixed1` and performed the bulk of this study, but then thought it would be prudent to explore the other alternatives). In terms of the Guassian displacement; the `gauss` option realizes it immediately, and after a sufficient number of steps, the second and third options also fulfill it via the law of large numbers.

#### Granularity of the simulation and how the simulated diffusion constant is varied

The simulated diffusion cofficient is proportional to: the movement variance per step divided by the length of time of each step. Thus the granularity of either (or both) of these may be adjusted to achieve a desired diffusion coefficient. Our software implementation allows for either the mean distance traveled per step or the length of each time step to be held constant during the simulation; while the value of the other parameter is adjusted to obtain a desired diffusion coefficient. When the mean distance traveled per step is to be held constant, its default value is 0.001*µ*m; and when the time step is to be held constant its default value is 1 millisecond.

#### Mean squared displacement of the particle under various movement options

To get a feel for how the simulation behaves, we tallied summary statistics of free random walks (collisions not considered) starting from the origin under six parameter setting: each of the three movement per step options {`Gauss, fixed, fixed1`} combined with two modes: `varydist` and `varytime`. Under varydist the simulator expects the time per step to be stipulated by the user and adjusts the movement per step accordingly to simulate a desired diffusion coefficient; conversely under the `varytime` option the user stipulates a movement size and the simulator adjusts the time step length as necessary. Table 1 shows summary statistics obtained by averaging over 100,000 random walks with a desired diffusion coefficient of 0.002*µ*/s, walking 1000 steps per walk. The program `printDisplacementStats` records the displacement along each axis at the end of each walk and computes the first three raw statistical moments. As expected for a random walk, the 1st and 3rd raw moments are close to zero along each of the (x,y,z) axes, and the 2nd raw moment (MSD) is as expected given the stipulated diffusion coefficient and amount of time simulated for each walk.

**Table 1:** Summary of results from three ways to move the particle in each step of the diffusion simulation. Run times are median wall clock time of three runs on a ThinkPad X1 Carbon machine running debian linux with 4 Intel(R) Core(TM) i7-337U cores. Results obtained by running `printDisplacementStats -r 1525947060 -t` DiffusorSpec `0.001 0.002 1000 100000`.

| DiffusorSpec | MSD<br>per step | Time<br>per step | Displacement raw moments (x,y,z) | | | MSD /<br>$6 \times \text{walkTime}$ | Wall clock<br>runtime |
| --- | --- | --- | --- | --- | --- | --- | --- |
|  |  |  | 1st (means) | 2nd (MSDs) | 3rd |  |  |
| varydist:gauss | 0.001 | $1.2 \times 10^5$ | -3.64E-5, 1.89E-7, 9.81E-5 | 0.00397, 0.00399, 0.00398 | -7.93E-7, 2.80E-6, -4.15E-6 | 0.00199 | 22.0s |
| varydist:fixed | 0.001 | $1.2 \times 10^5$ | 2.97E-5, 1.46E-4, -6.68E-6 | 0.00396, 0.00403, 0.00402 | -1.31E-7, 4.12E-6, -6.50E-7 | 0.00200 | 3.9s |
| varydist:fixed1 | 0.001 | $1.2 \times 10^5$ | -1.87E-4, -5.61E-5, -1.17E-4 | 0.00402, 0.00401, 0.00401 | 3.42E-6, -4.59E-6, -7.20E-6 | 0.00201 | 3.6s |
| varytime:gauss | 0.0833 | 0.001 | -3.32E-4, 1.72E-6, 8.96E-4 | 0.330, 0.333, 0.332 | -6.03E-4, 2.13E-4, 3.16E-4 | 0.00199 | 21.8s |
| varytime:fixed | 0.0833 | 0.001 | 2.71E-4, 1.34E-3, -6.10E-5 | 0.330, 0.336, 0.335 | -9.99E-5, 3.14E-3, 4.95E-4 | 0.00200 | 3.9s |
| varytime:fixed1 | 0.0833 | 0.001 | -1.71E-3, -5.12E-4, -1.06E-3 | 0.335, 0.334, 0.334 | 2.60E-3, -3.49E-3, -5.48E-3 | 0.00201 | 3.6s |

Our conclusions from these statistics is that a fixed length movement yields reasonably similar summary statistics to the Gaussian option, and runs several times faster. The option `fixed` which moves the particle a fixed distance along each axis in each step seems the most attractive as it is almost as fast as `fixed1` and may be a slightly better approximation. The drawback of the fixed length options is that they cause the particle to unnaturally jump discretely in space, moving on a kind of 3D grid. Fortunately however the mitochondria are randomly placed without respect to this 3D grid, so hopefully the grid does not cause any artifacts in terms of initial collisions with mitochondria.

## 5 Frequency of primary and secondary encounters with mitochondria

To understand the behavior of the simulation we investigated the number of steps needed for a diffusing mRNA molecule to encounter a mitochondria for the first, second and third times (with no anchoring). As shown in figures 6, 7, over random trials the distribution of the number of steps to initially reach a mitochondrion has a (somewhat left-skewed) bell shape when plotted as the logarithm of the number of steps. For a random walk stride of 0.01μm (diffusion constant of 0.01667μ²/s) plotted in the figure, a typical number of steps needed to initially encounter a mitochondrion is around 2^12^ to 2^19^ steps (roughly 4–524 seconds). However once the mRNA molecule encounters a mitochondrion, it very often bumps back into it right away, (30–35%) of the time on the first step! This strong tendency to quickly encounter the same mitochondrion again is to be expected. For example, consider the limit as the random walk stride becomes negligibly small relative to the radius of a mitochondrion. In that case the surface of the mitochondrion locally looks like a plane (in general at some angle not perfectly aligned with any of the axes of the random walk 3D lattice). Under those conditions any move of the mRNA is equally likely to move towards the surface of the mitochondrion as away from it. Occasionally however, the mRNA happens to move away from the immediate vicinity of the mitochondrion it just collided with and eventually encounters one of the other four mitochondria in the simulation. In this case the distribution (red bars at bottom of the second and third bars of the triples shown in figures 6, 7) of the number of steps taken is roughly similar to number of steps taken to initially reach a mitochondrion when starting from the nuclear exit point.

**Figure 6:**
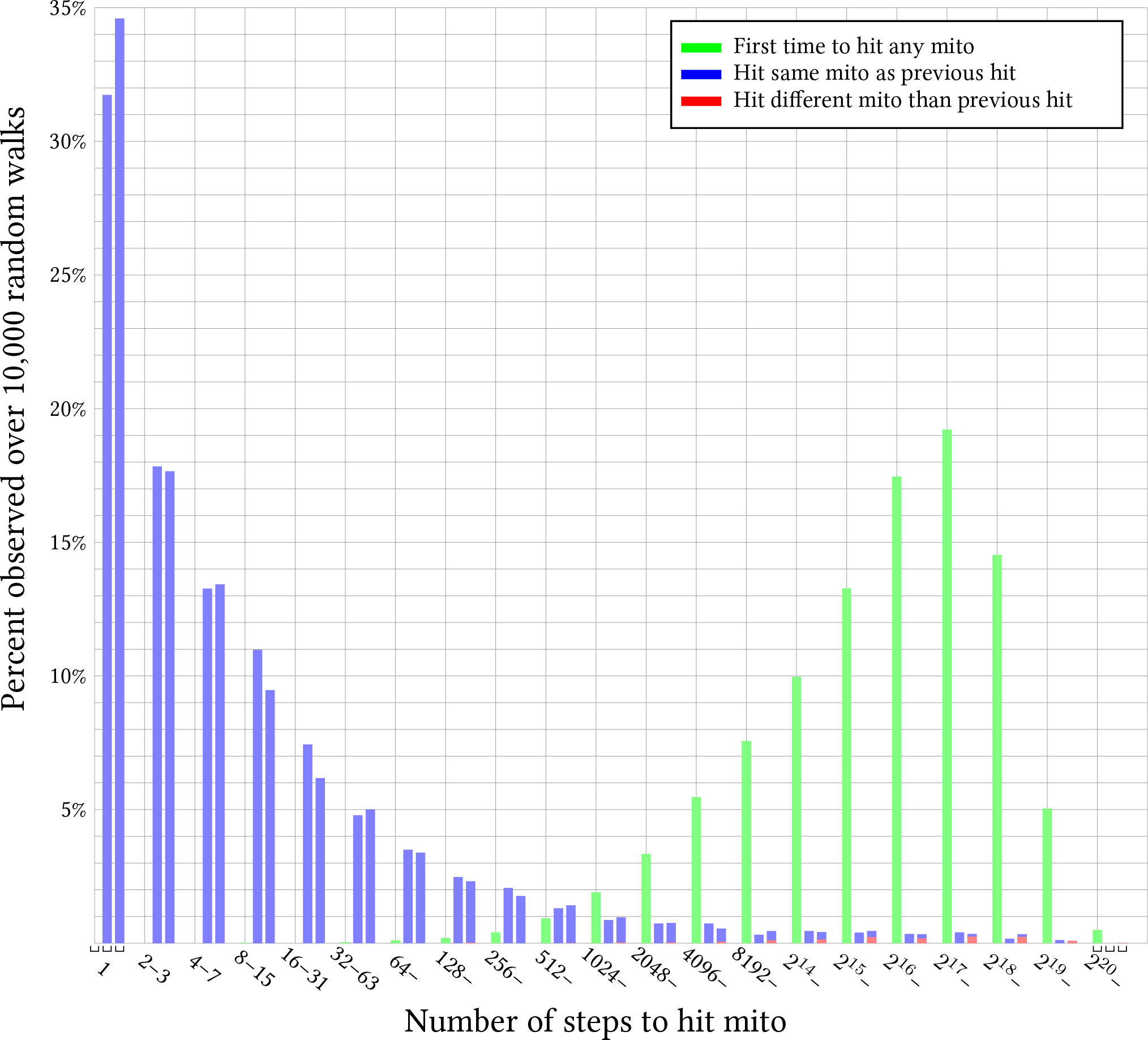
Histogram of the number of random walk steps to reach a mitochondrion for the first, second and third time are shown in triples (Using the “fixed1” diffusion movement option). Each triple sums together steps in a semi-closed range from [2^*x*^, 2^*x*+1^). In each triple the leftmost bar (in green) represents an initial random walk starting at the nuclear exit point. The second bar represents the number of additional steps to have a second mitochondrial encounter either with the same mitochondrion (blue) or a different one (red at bottom of bar). Similarly for the third bar, but for the steps between the second and third mitochondrial encounter. Missing bars (bars of zero length) represent events never observed during 10,000 total random trials. The data plotted here is based on the output of the command: `bin/printNumStepsTo-HitMitoThrice fixed1 -r 1503881565 0.01 10000`

**Figure 7:**
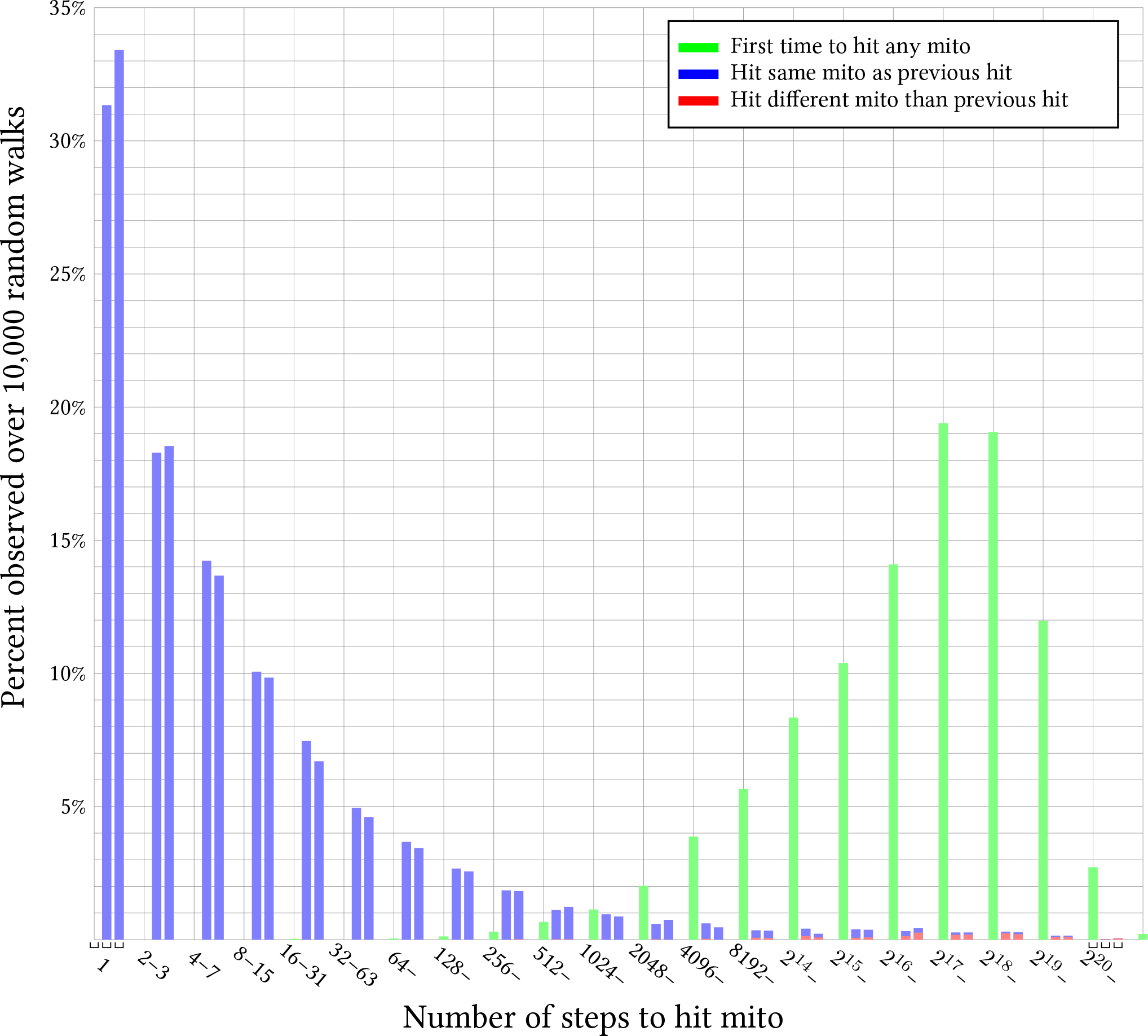
Histogram of the number of random walk steps to reach a mitochondrion for the first, second and third time are shown in triples (Using the “fixed” diffusion movement option). Each triple sums together steps in a semi-closed range from [2^*x*^, 2^*x*+1^). In each triple the leftmost bar (in green) represents an initial random walk starting at the nuclear exit point. The second bar represents the number of additional steps to have a second mitochondrial encounter either with the same mitochondrion (blue) or a different one (red at bottom of bar). Similarly for the third bar, but for the steps between the second and third mitochondrial encounter. Missing bars (bars of zero length) represent events never observed during 10,000 total random trials. The data plotted here is based on the output of the command: `bin/printNumStepsToHit-MitoThrice fixed -r 1552210278 0.01 10000`.

The two figures 6, 7 differ only in the movement options used. Although the distribution sampled under the fixed option seems to have a longer tail on the number of steps for the first encounter; other than that the two look qualitatively quite similar. This is reassuring, as subsequent figures in this manuscript were computed using `fixed1` movement (before we thought to investigate other movement strategies) and we hope the conclusions we draw from them do not depend on technical details of how the particle is moved during the simulation.

## 6 Feasible Range of Effective Diffusion Coefficient

To get an idea of what range of values for the effective diffusion coefficient might be worth further exploration we made a program which runs the diffusion simulation with anchoring turned off and with translation initiation having no effect. The results shown in figure 8 indicate that for a diffusion coefficient somewhere between 0.0001 and 0.0002*µ*m^2^/s, a molecule will encounter mitochondria in about 10% of runs, while at a diffusion coefficient of around 0.01*µ*m^2^/s the molecule will encounter mitochondria in about 90% of runs. As expected from the results of the previous section, at a given percentile value, the curve transitions sharply from zero mitochondrial encounters to many encounters.

**Figure 8:**
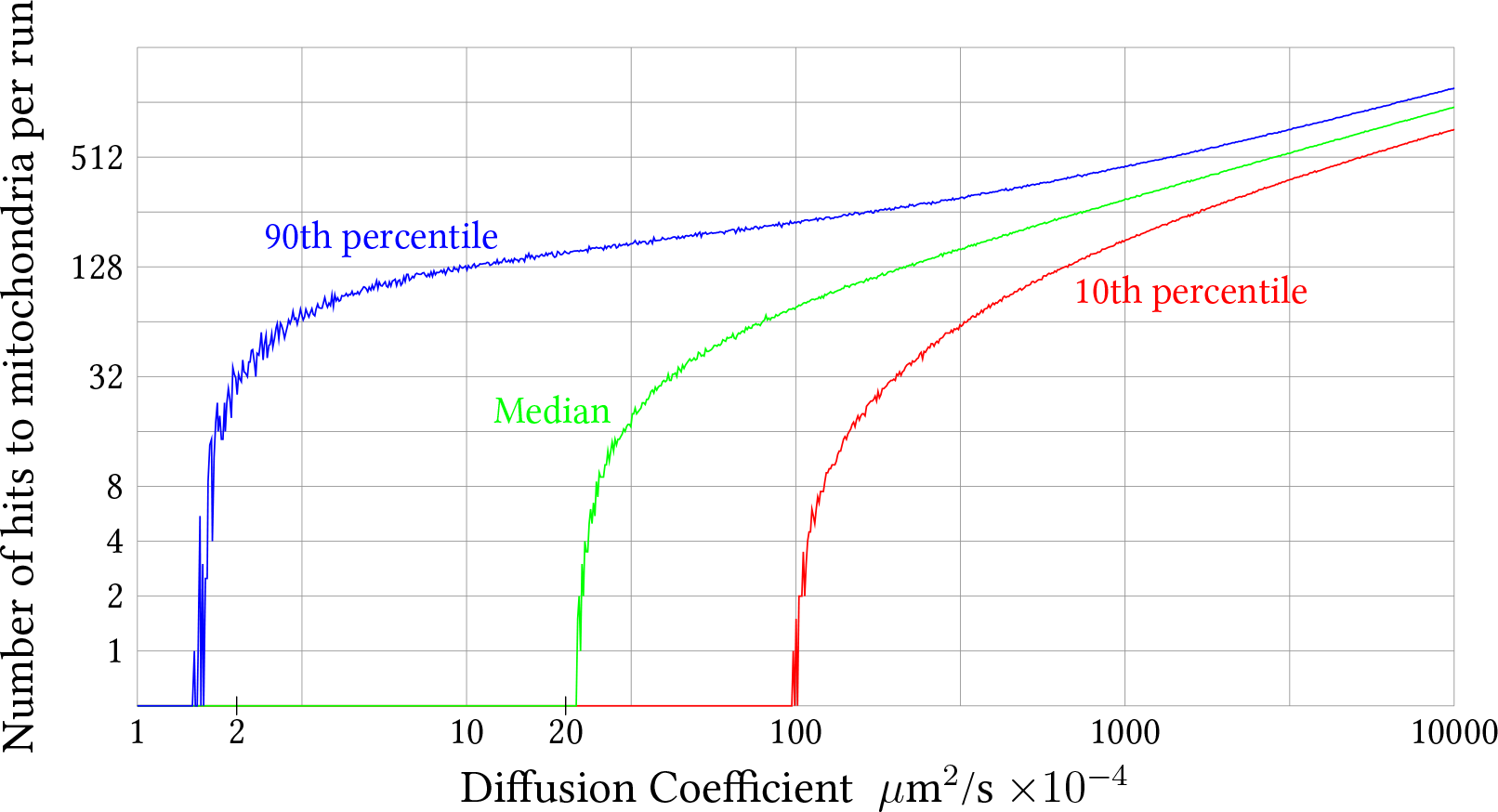
The relationship between diffusion coefficient and number of times a non-anchoring mRNA molecule encounters mitochondria is shown. The diffusion coefficient on the horizontal axis is on a decimal log scale, and the number of mitochondrial encounters on the vertical axis is on a binary log scale. The data for the plot was obtained by running the simulation with no mitochondrial anchoring and a single fixed diffusion coefficient per run (i.e. no effect of translation initiation). For each diffusion coefficient value (in units of *µ*m^2^/s) in the set: {1.0, 0.99, 0.99^2^, …, ≈ 0.0001}; 10,000 random simulation runs were performed and the number of hits to mitochondria in each run was recorded. The middle curve labeled “Median” shows the median value (i.e. 50th percentile), over the 10,000 runs performed for each diffusion coefficient value, of the number of times mitochondria were encountered per run. Similarly, the curves at left and right show the 90th and 10th percentiles respectively. For example, position 10 on the horizontal axis, for which the 90th percentile is approximately 128 and the median and 10th percentiles are zero, indicates that at a diffusion coefficient of 10 × 10^*−*4^*µ*m^2^/s = 0.001*µ*m^2^/s, most runs do not hit a mitochondria at all, but 10% of the runs hit a mitochondria 128 or more times. The data plotted here is based on the output of the command: `bin/recordNumMitHits -r 1507260043 1.0 0.0001 0.99 10000`

## 7 Virtual Genes Averaged by Translation Initiation Rate

As mentioned above, we wished to compare our simulation results with measured values on a linear scale. However, as can be seen in figure 4, at the level of individual genes there is a lot of scatter in the data, which would making it difficult to visually compare the data to simulated results. Therefore we smoothed the data by binning the genes by average translation initiation time. The first bin containing the 15 genes with the shortest average translation initiation time, the next bin containing the 15 genes with the next shortest time, up to the final 33th bin containing the 15 genes with the longest average translation initiation time. The effect is to smooth out the scatter in the plot of individual genes as shown in figure 9.

**Figure 9:**
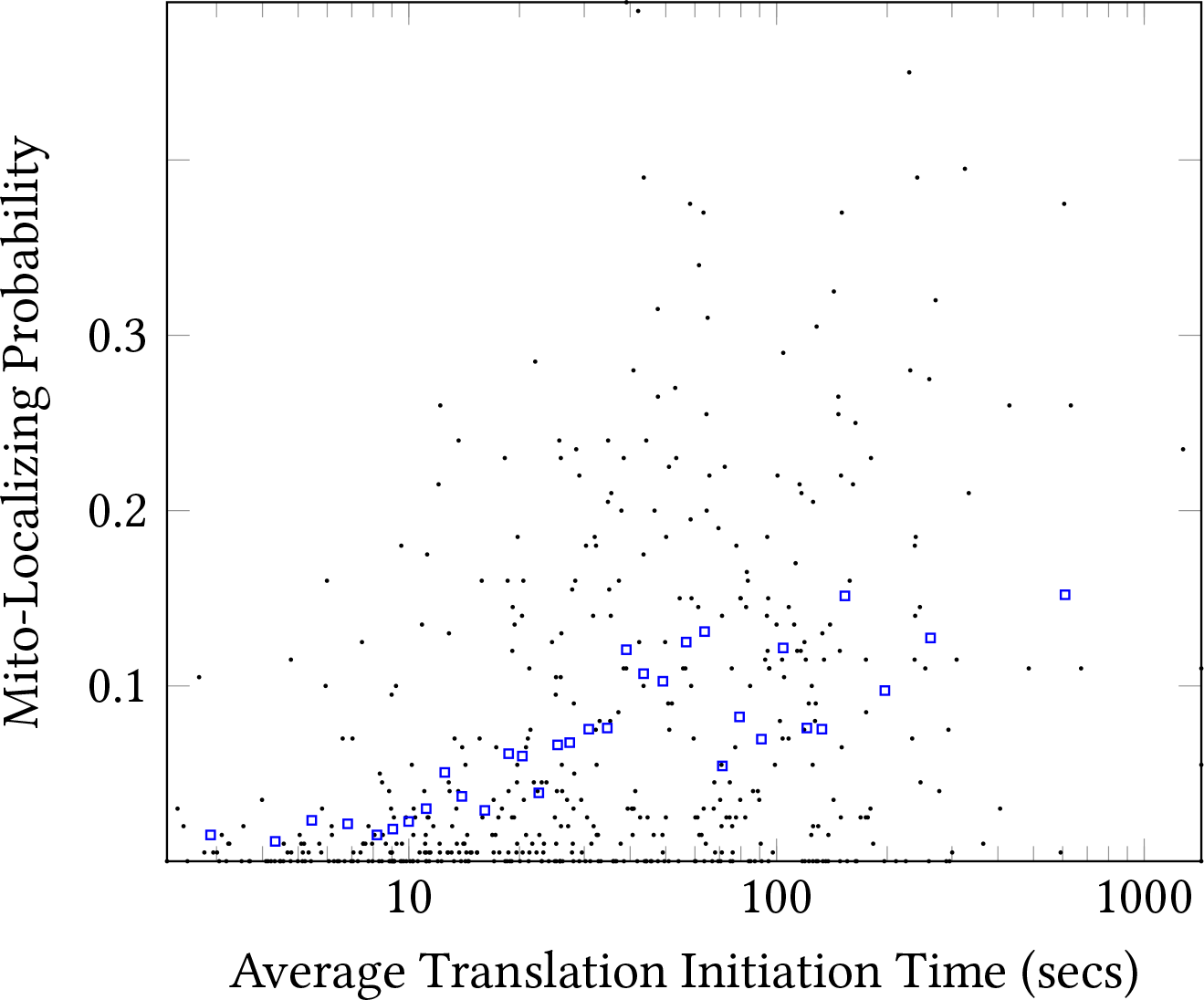
Scatter plot of the RPKM ratio based mitochondrial localizing probability and average translation initiation time. Each black dot represents one of 495 yeast whose protein products are imported into mitochondria. Each blue box represents the averaged values of 15 genes binned by average translation initiation time.

## 8 Exploration of Various Parameter Values

We performed a rough search over parameter value combinations (so called “grid search”) to explore the effect of varying the diffusion coefficient and anchoring probabilities before and after mRNA initiate translation. In the preceding sections we observed that (without anchoring) mRNA molecules seldom hit mitochondria a few times, but rather typically hit mitochondria either many times or not at all. Thus careful optimization of the anchoring probability is not meaningful. Instead we used the executable `doGridSearch` with a setting of 1000 repeated simulation runs per parameter setting to perform coarse-grained search over combinations of parameter values; obtaining the results shown in table 5 at the end of this document.

One clear trend from table 5 is that when the anchoring probability of not-yet-translating mRNA is zero, the correlation between simulated and measured mitochondrial localization is typically negative or at best near zero. This makes sense because if not-yet-translating mRNA never anchor to the mitochondria they encounter, the mRNA mitochondrial localization of genes with slow translation initiation would be expected to be lower than quickly initiating genes. Another clear trend is a requirement for a strong loss of mobility upon translation initiation to fit the data. The parameter values combinations which produced relatively good fit to sequencing based measurement based values (cyan rows in table 5) assume an increase in diffusion coefficient of 20–100x upon translation initiation.

### A closer look at two parameter value sets

Several combinations of parameter values yield simulation results with fairly good agreement to measured values. Indeed from the mean absolute error values one can see that the mitochondrial localization probabilities of the best ones were typically within about 5 percentage points of the bin-averaged virtual genes (cyan rows in table 5). To visualize the simulated versus measured mitochondrial localization probabilities we arbitrarily picked a couple of the good parameter sets (those marked in bold in table 5), and used walkGenes with a setting of 10,000 repeated runs to record the simulated mRNA mitochondrial localization probability for each bin-averaged virtual gene (figure 10).

**Figure 10:**
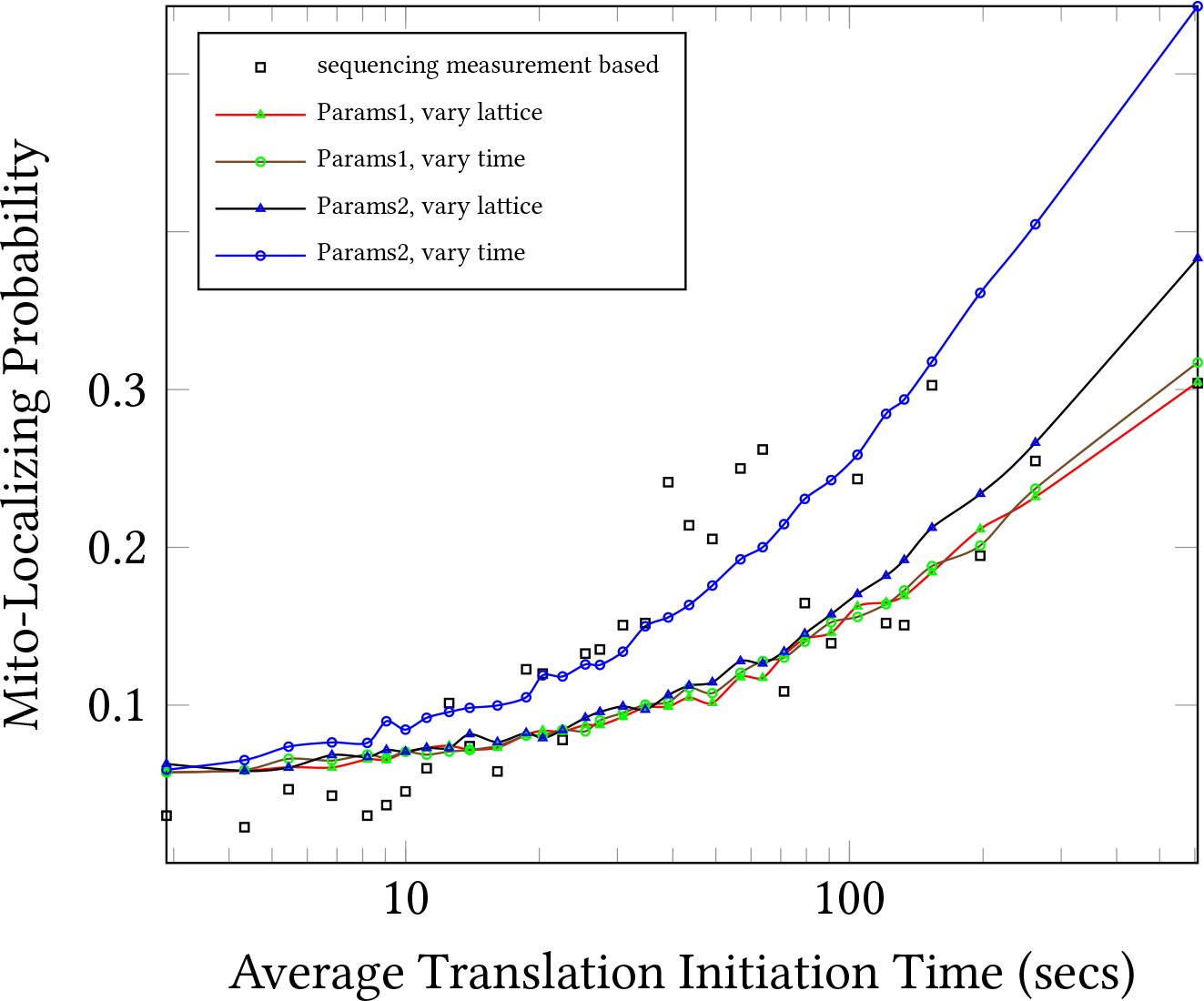
Measured and simulated mitochondrial localization probabilities (MLP) of bin-averaged virtual genes is shown. Simulated MLPs were obtained by running the walkGenes executable with 10,000. Params1 and Params2 represent (DC before, DC after, anchor prob before, anchor prob after) values of (0.0015625, 7.8125e-05, 1, 1) and (0.00625, 6.25e-05, 0.01, 1) respectively; where DC denotes diffusion coefficient in *µ*m^2^/s and before/after denote before and after initiating translation.

The simulated MLPs appear to match the measured ones reasonably well. At first glance it seems curious that the two curves for the “Params2” set of parameter values differs much more than the two curves for the “Params1” set, with the results simulated under the “vary time” exhibiting a higher MLP, especially for slow translation initiation time virtual genes. This seems to be an artifact of the simplistic way we defined the anchoring probability — as simply once chance to anchor per mitochondrial encounter with no adjustment for the time step length. The Params2 diffusion coefficient before translation initiation, 0.00625*µ*m^2^/s, is the same for both curves, but (under the default settings) the vary time option simulates that diffusion coefficient with a lattice size of 0.001μm and a time step 2.66667e-05s, while the vary lattice option uses a lattice size of 0.00612372*µ*m^2^/s and time step of 0.001s. With these time step lengths, the simulation under the vary time option will take 37.5x more steps per random walk than under the vary lattice option. With more fine grained steps the mRNA would be likely to rehit a mitochondrion more times and thus have more chances to anchor. This effect is not seen with Params1 because the anchor probability before initiating translation is 1, so mitochondria are only hit once in any case.

## 9 Simulation Software

For this follow-up study we re-implemented the simulation software with a modular design to facilitate readability and extensibility. This re-implementation essentially performs the same computation as the original one, but the software architecture is redesigned from scratch in an attempt to be easier to verify and extend. The main modules, auxiliary modules and executable are described in tables 2, 3 and 4 respectively; and their connections are schematically depicted in figure 11. The source code is available at: https://gitlab.com/paulhorton/diffusionsimulatormlr.

**Figure 11:**
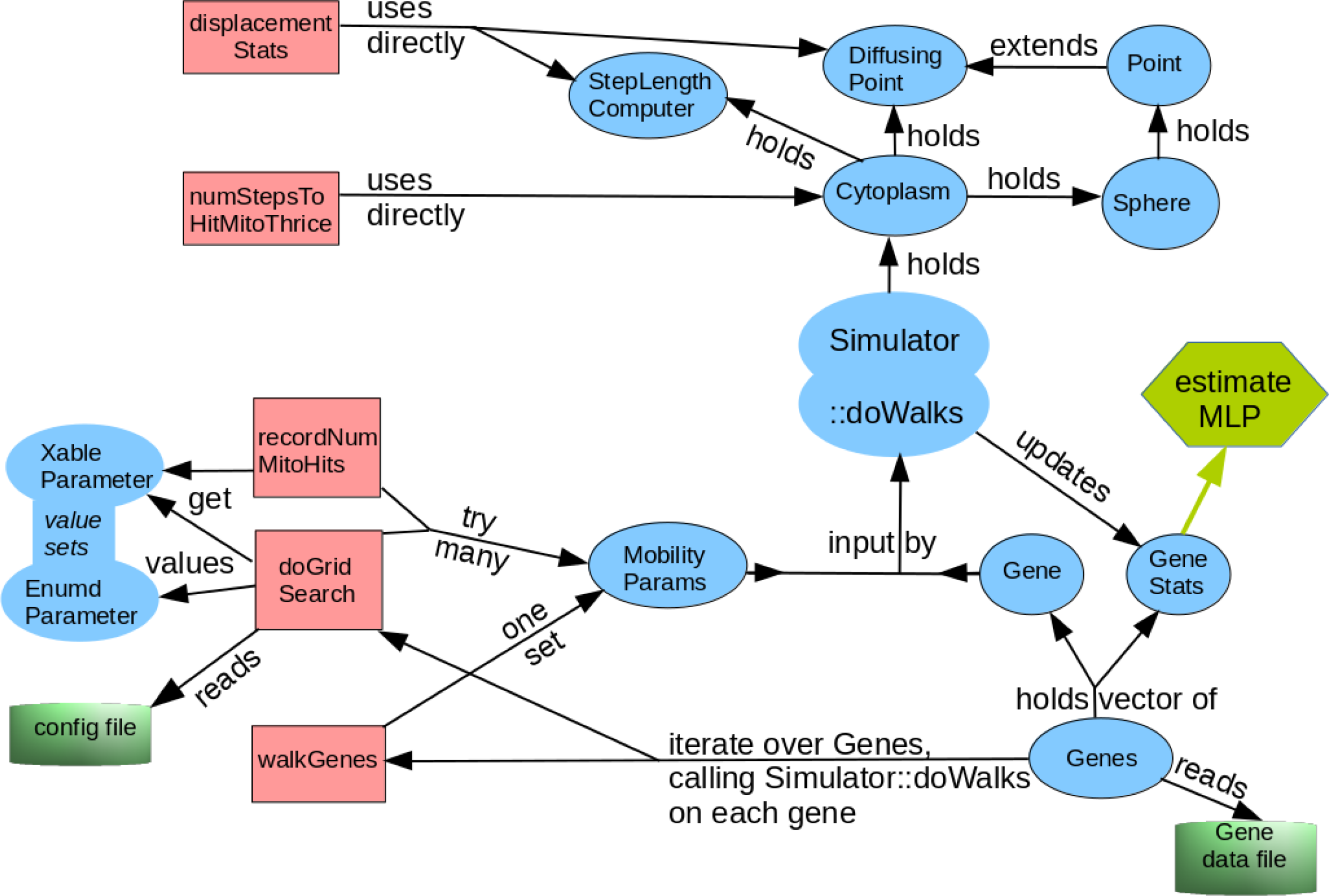
Schematic overview of the structure of the simulation software. Blue ovals, red rectangles and green cylinders represent classes, executables, and disk I/O respectively. The hexagon at right represents the estimation of statistics regarding the simulation results, particularly the mRNA mitochondrial localization probability. The labeled arrows represent interactions between modules. For example, “holds” represents composition; the `Sphere` class contains a `Point` object to represent its center and the `Cytoplasm` class contains `Sphere` objects representing mitochondria. In addition to the relationships shown, all of the executables (light green rectangles) use `ArgvParser_forSimulator` to parse command line arguments, but this has been omitted to reduce visual clutter in the schematic.

**Table 2:** The main classes used to implement the simulation are summarized.

|  |  |
| --- | --- |
| Point | Holds x,y,z coordinates. Knows how to measure its distance from another point. |
| DiffusingPoint | Extension of Point used to represent an mRNA molecule. Knows how to take one random step or undo its most recent step. |
| Sphere | Used to represent mitochondria. Holds Point origin and radius. Knows how to place itself randomly within a spherical shell and how to detect collisions with other Spheres or Points. |
| Cytoplasm | Holds DiffusingPoint mRNA, vector of Sphere mitochondria, and constants representing the size of the cell, nucleus, small and large mitochondria. Knows how to place mitochondria in random positions which do not overlap the nucleus or each other. Knows how to make the mRNA take a random step within the cytoplasm and report the collision if the step would have placed the mRNA inside a mitochondria. |
| Gene | Holds average translation initiation time and average number of ribosomes for a gene. Knows how to compute the translation initiation probability per second from that. |
| GeneStats | Used to record statistics such as the number of anchoring attempts and successes for a gene (over many simulation runs). |
| Genes | Holds vector of Gene, GeneStats pairs. Knows how to read Gene info in from a file. Knows how to estimate the simulated mitochondrial localization probability of the genes it holds based on the statistics accumulated during simulation runs held in its GeneStats vector. |
| StepLengthComputer | Functions to adjust time step or movement step size to match a desired diffusion coefficient. Keeps either the time or the lattice step length constant, depending on if it was constructed with a “varyGrid” (grid means lattice here) or a “varyTime” option. |
| Simulator | Holds Cytoplasm and StepLengthComputer pointer. Provides method doWalks which walks a Gene according to the parameters passed as a MobilityParams reference and updates a GeneStats reference in accordance with the results of each walk. |
| MobilityParams | Holds gene-independent simulation parameters: diffusion coefficient and mitochondrial anchor probability for mRNA before and after initiating translation. |

**Table 3:** Auxiliary classes and files used to implement the simulation are summarized.

|  |  |
| --- | --- |
| ArgvParser | Simple, generic command line argument parser. |
| ArgvParser_<br>forSimulator | Extension of ArgvParser which handles and records the number used to seed the random generator. Convenient for reproducibility. |
| EnumdParam | For parameter search. Supports iteration over parameters which should take one of several enumerated values, e.g. `anchorProb` $\in \{0, 0.01, 0.1, 1\}$. Knows how to construct itself from a string. |
| XableParam | For parameter search. Supports iteration over parameters which should be varied multiplicatively over a range of values. For example setting diffusion coefficient to 0.2, 0.1, 0.05, ..., 0.0015625. Knows how to construct itself from a string. |
| commonMacros<br>UsingTypedefs.hh | Provides some common preprocessor Macros, C++ using and typedef statements. |
| utils.hh utils.cc | Provide some simple utility functions. |

**Table 4:**
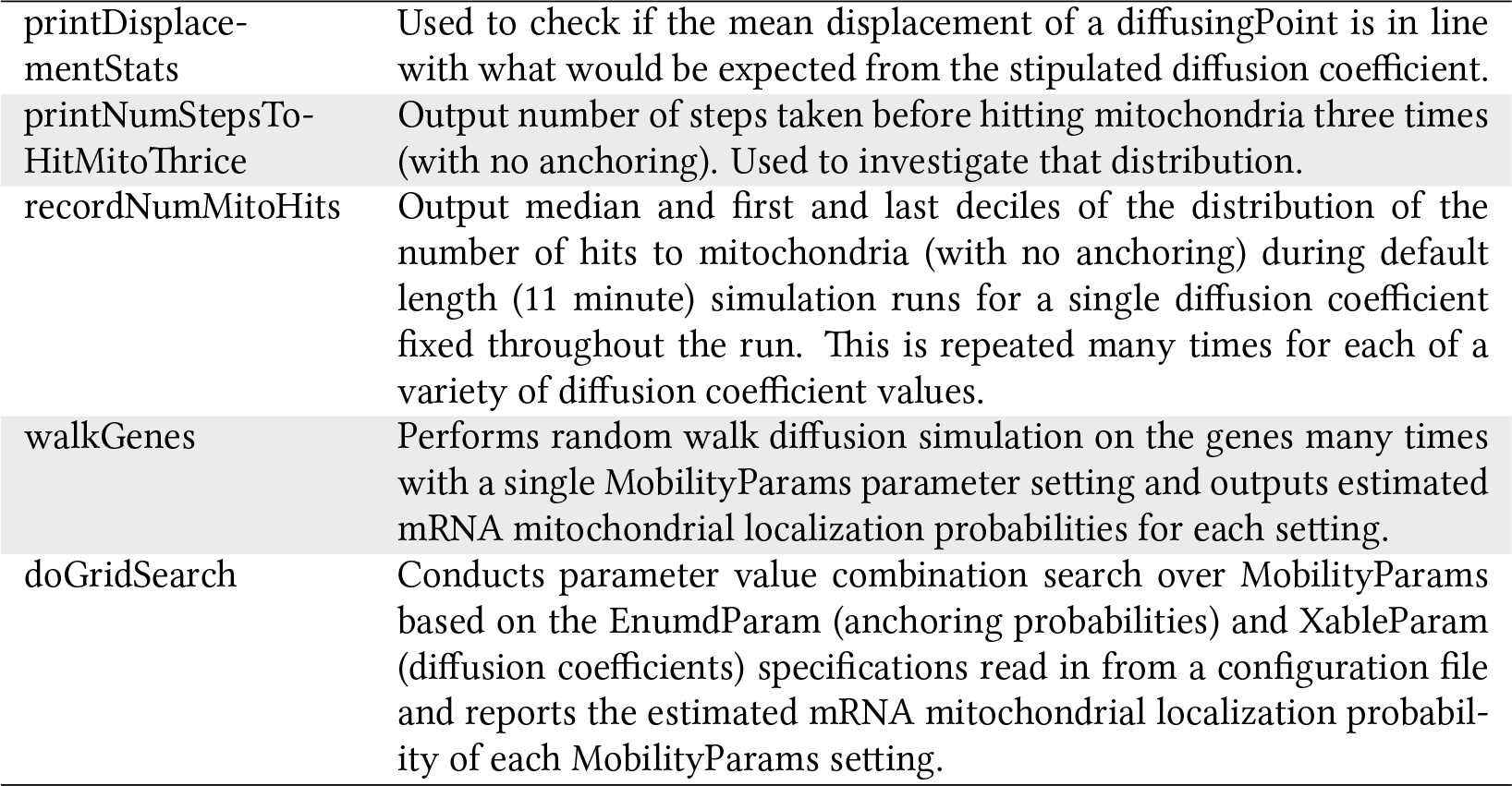
Executable programs used to generate the data in this manuscript.

**Table 5:** Results of parameter value grid search using the program doGridSearch with a constant time step size of 0.001 seconds. “free” denotes the state of mRNA before initiating translation (free of ribosomes). “Sim. MLP” denotes simulated mitochondrial localization probability estimated using 1000 random walks per parameter setting. Sim. vs. Measured MLP correlation and MAE are the Pearson’s correlation and MAE (mean absolute error) of the simulated mitochondrial localization probability compared to the probabilities derived from the experimental measurement of Williams et al. Rows in gray indicate parameter settings producing with a negative correlation; while rows in cyan indicate parameter settings with MAE of 0.07 or less. Of the cyan rows, the two in bold text indicate parameter settings further examined in this manuscript.

| Free<br>Diff. Coeff. | Translating<br>Diff. Coeff. | Free<br>anchor prob. | Translating<br>anchor prob. | Sim. MLP<br>mean | Sim. vs. Measured MLP<br>correlation | MAE |
| --- | --- | --- | --- | --- | --- | --- |
| 0.2 | 0.05 | 0 | 0.1 | 0.98 | -0.45 | 0.84 |
| 0.2 | 0.05 | 0 | 1 | 0.98 | -0.43 | 0.84 |
| 0.2 | 0.05 | 0.01 | 0.1 | 1.00 | -0.04 | 0.86 |
| 0.2 | 0.05 | 0.01 | 1 | 1.00 | -0.23 | 0.86 |
| 0.2 | 0.05 | 0.1 | 0.1 | 1.00 | 0.12 | 0.86 |
| 0.2 | 0.05 | 0.1 | 1 | 1.00 | 0.29 | 0.86 |
| 0.2 | 0.05 | 1 | 1 | 1.00 | 0.28 | 0.86 |
| 0.2 | 0.01 | 0 | 0.1 | 0.83 | -0.59 | 0.69 |
| 0.2 | 0.01 | 0 | 1 | 0.86 | -0.57 | 0.72 |
| 0.2 | 0.01 | 0.01 | 0.1 | 0.91 | 0.73 | 0.77 |
| 0.2 | 0.01 | 0.01 | 1 | 0.92 | 0.75 | 0.78 |
| 0.2 | 0.01 | 0.1 | 0.1 | 0.94 | 0.82 | 0.80 |
| 0.2 | 0.01 | 0.1 | 1 | 0.95 | 0.78 | 0.81 |
| 0.2 | 0.01 | 1 | 1 | 0.96 | 0.89 | 0.82 |
| 0.2 | 0.002 | 0 | 0.1 | 0.41 | -0.68 | 0.27 |
| 0.2 | 0.002 | 0 | 1 | 0.42 | -0.69 | 0.29 |
| 0.2 | 0.002 | 0.01 | 0.1 | 0.59 | 0.80 | 0.45 |
| 0.2 | 0.002 | 0.01 | 1 | 0.62 | 0.80 | 0.47 |
| 0.2 | 0.002 | 0.1 | 0.1 | 0.75 | 0.85 | 0.61 |
| 0.2 | 0.002 | 0.1 | 1 | 0.76 | 0.86 | 0.62 |
| 0.2 | 0.002 | 1 | 1 | 0.81 | 0.85 | 0.67 |
| 0.1 | 0.025 | 0 | 0.1 | 0.96 | -0.51 | 0.82 |
| 0.1 | 0.025 | 0 | 1 | 0.97 | -0.47 | 0.83 |
| 0.1 | 0.025 | 0.01 | 0.1 | 0.99 | 0.17 | 0.85 |
| 0.1 | 0.025 | 0.01 | 1 | 0.99 | 0.03 | 0.85 |
| 0.1 | 0.025 | 0.1 | 0.1 | 0.99 | 0.61 | 0.85 |
| 0.1 | 0.025 | 0.1 | 1 | 1.00 | 0.54 | 0.85 |
| 0.1 | 0.025 | 1 | 1 | 1.00 | 0.46 | 0.86 |
| 0.1 | 0.005 | 0 | 0.1 | 0.65 | -0.59 | 0.51 |
| 0.1 | 0.005 | 0 | 1 | 0.68 | -0.59 | 0.54 |
| 0.1 | 0.005 | 0.01 | 0.1 | 0.76 | 0.72 | 0.62 |
| 0.1 | 0.005 | 0.01 | 1 | 0.77 | 0.71 | 0.63 |
| 0.1 | 0.005 | 0.1 | 0.1 | 0.82 | 0.83 | 0.68 |
| 0.1 | 0.005 | 0.1 | 1 | 0.84 | 0.82 | 0.70 |
| 0.1 | 0.005 | 1 | 1 | 0.86 | 0.84 | 0.72 |
| 0.1 | 0.001 | 0 | 0.1 | 0.27 | -0.61 | 0.14 |
| 0.1 | 0.001 | 0 | 1 | 0.28 | -0.58 | 0.16 |
| 0.1 | 0.001 | 0.01 | 0.1 | 0.45 | 0.79 | 0.31 |
| 0.1 | 0.001 | 0.01 | 1 | 0.46 | 0.77 | 0.32 |
| 0.1 | 0.001 | 0.1 | 0.1 | 0.62 | 0.86 | 0.48 |
| 0.1 | 0.001 | 0.1 | 1 | 0.63 | 0.84 | 0.48 |
| 0.1 | 0.001 | 1 | 1 | 0.68 | 0.85 | 0.54 |
| 0.05 | 0.0125 | 0 | 0.1 | 0.88 | -0.56 | 0.74 |
| 0.05 | 0.0125 | 0 | 1 | 0.9 | -0.56 | 0.76 |
| 0.05 | 0.0125 | 0.01 | 0.1 | 0.92 | 0.15 | 0.78 |
| 0.05 | 0.0125 | 0.01 | 1 | 0.94 | 0.26 | 0.80 |
| 0.05 | 0.0125 | 0.1 | 0.1 | 0.94 | 0.73 | 0.80 |
| 0.05 | 0.0125 | 0.1 | 1 | 0.95 | 0.76 | 0.81 |
| 0.05 | 0.0125 | 1 | 1 | 0.96 | 0.69 | 0.81 |
| 0.05 | 0.0025 | 0 | 0.1 | 0.46 | -0.64 | 0.33 |
| 0.05 | 0.0025 | 0 | 1 | 0.49 | -0.60 | 0.35 |
| 0.05 | 0.0025 | 0.01 | 0.1 | 0.57 | 0.74 | 0.43 |
| 0.05 | 0.0025 | 0.01 | 1 | 0.59 | 0.72 | 0.45 |
| 0.05 | 0.0025 | 0.1 | 0.1 | 0.67 | 0.81 | 0.53 |
| 0.05 | 0.0025 | 0.1 | 1 | 0.68 | 0.79 | 0.54 |
| 0.05 | 0.0025 | 1 | 1 | 0.71 | 0.82 | 0.57 |
| 0.05 | 0.0005 | 0 | 0.1 | 0.18 | -0.58 | 0.09 |
| 0.05 | 0.0005 | 0 | 1 | 0.19 | -0.72 | 0.10 |
| 0.05 | 0.0005 | 0.01 | 0.1 | 0.34 | 0.76 | 0.20 |
| 0.05 | 0.0005 | 0.01 | 1 | 0.34 | 0.77 | 0.20 |
| 0.05 | 0.0005 | 0.1 | 0.1 | 0.49 | 0.83 | 0.35 |
| 0.05 | 0.0005 | 0.1 | 1 | 0.50 | 0.83 | 0.35 |
| 0.05 | 0.0005 | 1 | 1 | 0.54 | 0.84 | 0.40 |
| 0.025 | 0.00625 | 0 | 0.1 | 0.715 | -0.61 | 0.57 |
| 0.025 | 0.00625 | 0 | 1 | 0.742 | -0.60 | 0.60 |
| 0.025 | 0.00625 | 0.01 | 0.1 | 0.765 | 0.40 | 0.62 |
| 0.025 | 0.00625 | 0.01 | 1 | 0.792 | 0.02 | 0.65 |
| 0.025 | 0.00625 | 0.1 | 0.1 | 0.805 | 0.78 | 0.66 |
| 0.025 | 0.00625 | 0.1 | 1 | 0.828 | 0.70 | 0.69 |
| 0.025 | 0.00625 | 1 | 1 | 0.835 | 0.72 | 0.69 |
| 0.025 | 0.00125 | 0 | 0.1 | 0.313 | -0.72 | 0.19 |
| 0.025 | 0.00125 | 0 | 1 | 0.327 | -0.64 | 0.20 |
| 0.025 | 0.00125 | 0.01 | 0.1 | 0.416 | 0.75 | 0.28 |
| 0.025 | 0.00125 | 0.01 | 1 | 0.426 | 0.69 | 0.29 |
| 0.025 | 0.00125 | 0.1 | 0.1 | 0.509 | 0.78 | 0.37 |
| 0.025 | 0.00125 | 0.1 | 1 | 0.522 | 0.77 | 0.38 |
| 0.025 | 0.00125 | 1 | 1 | 0.539 | 0.80 | 0.40 |
| 0.025 | 0.00025 | 0 | 0.1 | 0.119 | -0.68 | 0.08 |
| 0.025 | 0.00025 | 0 | 1 | 0.125 | -0.72 | 0.08 |
| 0.025 | 0.00025 | 0.01 | 0.1 | 0.246 | 0.73 | 0.11 |
| 0.025 | 0.00025 | 0.01 | 1 | 0.252 | 0.71 | 0.12 |
| 0.025 | 0.00025 | 0.1 | 0.1 | 0.373 | 0.80 | 0.23 |
| 0.025 | 0.00025 | 0.1 | 1 | 0.375 | 0.80 | 0.23 |
| 0.025 | 0.00025 | 1 | 1 | 0.408 | 0.80 | 0.27 |
| 0.0125 | 0.00313 | 0 | 0.1 | 0.523 | -0.68 | 0.38 |
| 0.0125 | 0.00313 | 0 | 1 | 0.551 | -0.64 | 0.41 |
| 0.0125 | 0.00313 | 0.01 | 0.1 | 0.578 | 0.36 | 0.44 |
| 0.0125 | 0.00313 | 0.01 | 1 | 0.602 | 0.44 | 0.46 |
| 0.0125 | 0.00313 | 0.1 | 0.1 | 0.626 | 0.70 | 0.49 |
| 0.0125 | 0.00313 | 0.1 | 1 | 0.641 | 0.70 | 0.50 |
| 0.0125 | 0.00313 | 1 | 1 | 0.656 | 0.76 | 0.52 |
| 0.0125 | 0.000625 | 0 | 0.1 | 0.216 | -0.67 | 0.11 |
| 0.0125 | 0.000625 | 0 | 1 | 0.219 | -0.71 | 0.11 |
| 0.0125 | 0.000625 | 0.01 | 0.1 | 0.297 | 0.66 | 0.16 |
| 0.0125 | 0.000625 | 0.01 | 1 | 0.302 | 0.68 | 0.16 |
| 0.0125 | 0.000625 | 0.1 | 0.1 | 0.374 | 0.77 | 0.23 |
| 0.0125 | 0.000625 | 0.1 | 1 | 0.376 | 0.76 | 0.24 |
| 0.0125 | 0.000625 | 1 | 1 | 0.399 | 0.77 | 0.26 |
| 0.0125 | 0.000125 | 0 | 0.1 | 0.080 | -0.63 | 0.09 |
| 0.0125 | 0.000125 | 0 | 1 | 0.082 | -0.75 | 0.09 |
| 0.0125 | 0.000125 | 0.01 | 0.1 | 0.174 | 0.73 | 0.06 |
| 0.0125 | 0.000125 | 0.01 | 1 | 0.175 | 0.70 | 0.06 |
| 0.0125 | 0.000125 | 0.1 | 0.1 | 0.274 | 0.80 | 0.13 |
| 0.0125 | 0.000125 | 0.1 | 1 | 0.27 | 0.77 | 0.13 |
| 0.0125 | 0.000125 | 1 | 1 | 0.296 | 0.79 | 0.15 |
| 0.00625 | 0.00156 | 0 | 0.1 | 0.37 | -0.72 | 0.24 |
| 0.00625 | 0.00156 | 0 | 1 | 0.389 | -0.69 | 0.26 |
| 0.00625 | 0.00156 | 0.01 | 0.1 | 0.407 | 0.36 | 0.27 |
| 0.00625 | 0.00156 | 0.01 | 1 | 0.426 | 0.36 | 0.29 |
| 0.00625 | 0.00156 | 0.1 | 0.1 | 0.447 | 0.67 | 0.31 |
| 0.00625 | 0.00156 | 0.1 | 1 | 0.463 | 0.69 | 0.32 |
| 0.00625 | 0.00156 | 1 | 1 | 0.476 | 0.71 | 0.34 |
| 0.00625 | 0.000313 | 0 | 0.1 | 0.146 | -0.53 | 0.08 |
| 0.00625 | 0.000313 | 0 | 1 | 0.151 | -0.67 | 0.09 |
| 0.00625 | 0.000313 | 0.01 | 0.1 | 0.202 | 0.63 | 0.08 |
| 0.00625 | 0.000313 | 0.01 | 1 | 0.214 | 0.72 | 0.09 |
| 0.00625 | 0.000313 | 0.1 | 0.1 | 0.26 | 0.74 | 0.12 |
| 0.00625 | 0.000313 | 0.1 | 1 | 0.266 | 0.73 | 0.13 |
| 0.00625 | 0.000313 | 1 | 1 | 0.277 | 0.75 | 0.14 |
| 0.00625 | 6.25e-05 | 0 | 0.1 | 0.052 | -0.53 | 0.10 |
| 0.00625 | 6.25e-05 | 0 | 1 | 0.053 | -0.66 | 0.10 |
| 0.00625 | 6.25e-05 | 0.01 | 0.1 | 0.12 | 0.73 | 0.05 |
| **0.00625** | **6.25e-05** | **0.01** | **1** | **0.12** | **0.74** | **0.05** |
| 0.00625 | 6.25e-05 | 0.1 | 0.1 | 0.185 | 0.78 | 0.06 |
| 0.00625 | 6.25e-05 | 0.1 | 1 | 0.185 | 0.80 | 0.06 |
| 0.00625 | 6.25e-05 | 1 | 1 | 0.201 | 0.78 | 0.07 |
| 0.00313 | 0.000781 | 0 | 0.1 | 0.25 | -0.71 | 0.14 |
| 0.00313 | 0.000781 | 0 | 1 | 0.27 | -0.67 | 0.15 |
| 0.00313 | 0.000781 | 0.01 | 0.1 | 0.29 | 0.46 | 0.15 |
| 0.00313 | 0.000781 | 0.01 | 1 | 0.30 | 0.39 | 0.16 |
| 0.00313 | 0.000781 | 0.1 | 0.1 | 0.32 | 0.71 | 0.17 |
| 0.00313 | 0.000781 | 0.1 | 1 | 0.33 | 0.69 | 0.19 |
| 0.00313 | 0.000781 | 1 | 1 | 0.33 | 0.71 | 0.19 |
| 0.00313 | 0.000156 | 0 | 0.1 | 0.10 | -0.49 | 0.08 |
| 0.00313 | 0.000156 | 0 | 1 | 0.1 | -0.56 | 0.08 |
| 0.00313 | 0.000156 | 0.01 | 0.1 | 0.14 | 0.71 | 0.05 |
| 0.00313 | 0.000156 | 0.01 | 1 | 0.14 | 0.68 | 0.05 |
| 0.00313 | 0.000156 | 0.1 | 0.1 | 0.17 | 0.79 | 0.05 |
| 0.00313 | 0.000156 | 0.1 | 1 | 0.18 | 0.77 | 0.06 |
| 0.00313 | 0.000156 | 1 | 1 | 0.18 | 0.78 | 0.06 |
| 0.00313 | 3.13e-05 | 0 | 0.1 | 0.03 | -0.10 | 0.11 |
| 0.00313 | 3.13e-05 | 0 | 1 | 0.04 | 0.03 | 0.11 |
| 0.00313 | 3.13e-05 | 0.01 | 0.1 | 0.08 | 0.76 | 0.06 |
| 0.00313 | 3.13e-05 | 0.01 | 1 | 0.08 | 0.72 | 0.06 |
| 0.00313 | 3.13e-05 | 0.1 | 0.1 | 0.12 | 0.77 | 0.05 |
| 0.00313 | 3.13e-05 | 0.1 | 1 | 0.12 | 0.76 | 0.05 |
| 0.00313 | 3.13e-05 | 1 | 1 | 0.13 | 0.77 | 0.05 |
| 0.00156 | 0.000391 | 0 | 0.1 | 0.17 | -0.57 | 0.09 |
| 0.00156 | 0.000391 | 0 | 1 | 0.18 | -0.72 | 0.09 |
| 0.00156 | 0.000391 | 0.01 | 0.1 | 0.20 | 0.56 | 0.08 |
| 0.00156 | 0.000391 | 0.01 | 1 | 0.21 | 0.41 | 0.09 |
| 0.00156 | 0.000391 | 0.1 | 0.1 | 0.22 | 0.73 | 0.09 |
| 0.00156 | 0.000391 | 0.1 | 1 | 0.22 | 0.67 | 0.09 |
| 0.00156 | 0.000391 | 1 | 1 | 0.22 | 0.76 | 0.09 |
| 0.00156 | 7.81e-05 | 0 | 0.1 | 0.06 | 0.07 | 0.09 |
| 0.00156 | 7.81e-05 | 0 | 1 | 0.06 | -0.30 | 0.09 |
| 0.00156 | 7.81e-05 | 0.01 | 0.1 | 0.09 | 0.69 | 0.07 |
| 0.00156 | 7.81e-05 | 0.01 | 1 | 0.09 | 0.70 | 0.06 |
| 0.00156 | 7.81e-05 | 0.1 | 0.1 | 0.11 | 0.72 | 0.05 |
| 0.00156 | 7.81e-05 | 0.1 | 1 | 0.11 | 0.73 | 0.05 |
| **0.00156** | **7.81e-05** | **1** | **1** | **0.12** | **0.78** | **0.04** |
| 0.00156 | 1.56e-05 | 0 | 0.1 | 0.02 | 0.09 | 0.12 |
| 0.00156 | 1.56e-05 | 0 | 1 | 0.02 | 0.07 | 0.12 |
| 0.00156 | 1.56e-05 | 0.01 | 0.1 | 0.05 | 0.74 | 0.09 |
| 0.00156 | 1.56e-05 | 0.01 | 1 | 0.05 | 0.74 | 0.09 |
| 0.00156 | 1.56e-05 | 0.1 | 0.1 | 0.08 | 0.73 | 0.06 |
| 0.00156 | 1.56e-05 | 0.1 | 1 | 0.08 | 0.73 | 0.06 |
| 0.00156 | 1.56e-05 | 1 | 1 | 0.08 | 0.76 | 0.06 |

### Technical Notes on Implementation

The choice of spheres to represent all objects with volume simplified collision detection, since computing the distance between a point and the center of a sphere is sufficient to determine if the point is inside the sphere.

Random placement of mitochondria is done sequentially by rejection sampling; the center of a mitochondrion is tentatively randomly placed inside a cube encompassing the virtual cell by independently selecting *x*, *y* and *z* coordinates according to a uniform distribution and accepting that tentative position if the mitochondrion is completely in the cytosol and not overlapping the nucleus or any previously placed mitochondria, otherwise the random placement is repeated until a viable position for the mitochondrion is obtained.

In our simulation scheme, the mRNA diffusion into the cytosol starts at the surface of the nucleus. Due to symmetry, any point on the surface of the nucleus should yield the same results. For convenience the software always starts the mRNA at the point (1,0,0).

### Software Execution Time and Hardware Requirements

In this study we were not focused on optimizing running time, and for example did not implement multi-threading. However the computations presented in this manuscript were manageable. For example, using the executable `walkGenes` to run the simulation 10,000 times with the time step size fixed at 0.001s required approximately 3 hours on a laptop PC running Linux.

### Reproducibility/Extensibility of this document

We made an extra effort to make this document reproducible and extensible. In particular, all data, code and drawings used to produce the figures here are made available as supplementary files. In principle, anyone familiar with using `LaTeX` should not only be able to reproduce the figures in this document, but also customize them. Readers wishing to do so should start by looking at the `.tex` document source and `makefile` supplementary files.

## 10 Discussion

### Possible Future Extensions to the Simulation

Several extensions could be made to incrementally improve the simulation:

- The anchor probability could be made a function of the time step length to address the scale-effect seen in the Params2 parameter setting of figure 10.
- Gene-specific mRNA half-life values could be used instead of using 11 minutes for all genes.
- The anchor probability and diffusion coefficient of translating mRNA could be adjusted for number of ribosomes.

The modular design of the software would make these improvements simple to add. However, we do not expect that those improvements would significantly alter the conclusions drawn here, so we leave them for potential future work.

## 11 Conclusions

We conducted a mRNA diffusion simulation study to more thoroughly investigate the results of an initial simulation study. Our main conclusions are that if a change in mobility upon translation initiation is a significant factor in explaining the negative correlation between mRNA fast translation initiation and mitochondrial localization then:

- Even mRNA which are not yet translating when approaching a mitochondrial surface must have a non-zero probability of anchoring there.
- The effective diffusion coefficient of mRNA must be significantly higher (20–100x) before initiating translation than afterwards.

## 12 Acknowledgements

This study borrows very heavily on ideas from an initial simulation study conceived of and performed by Thomas Poulsen. Some of the data used in this follow-up study was initially gathered by Kenichiro Imai and Thomas Poulsen. Martin Frith read parts of this manuscript and gave helpful feedback.

## Appendix: Comparison with Initial Study

### Methodology

Although based on the initial study, the methodology used here differs from the initial study [1] in several ways as summarized in table 6. Perhaps the most important difference is that this study allows for semi-quantitative comparison between measurements of mitochondrial localization (the sequencing based mRNA localization measurements [5] handled on a linear scale) as opposed to the initial study which compares simulation results to microarray derived results [4] on a logarithmic scale.

**Table 6:** Comparison of initial and follow-up mRNA diffusion simulation study. “DC” denotes diffusion coefficient. “free mRNA” are mRNA molecules which have not yet initiated translation.

| Study | Initial | Follow-up |
| --- | --- | --- |
| **Measured MLP source** | Microarray based | Sequencing Based |
| **Results comparison scale** | Logarithmic | Linear |
| **mRNA mobility** | DC & Entrapment | Effective DC |
| **mRNA start at** | Center of cell | Surface of nucleus |
| **When free mRNA Hit Mito.** | Usually initiate translation | Have some chance to anchor |
| **Parameter values** | Some variations tried | Grid Search over Combinations |
| **Software design** | Integrated | Modular |

### Conclusions

The two main conclusions of this follow-up study are qualitatively consistent with the initial study. Below we repeat the main conclusions of this study in the context of comparison with the initial study.

### Non-translating mRNA must be able to anchor to mitochondria

The first conclusion is that mRNA which are not yet translating when approaching a mitochondrial surface must have a non-zero probability of anchoring. The initial study does not included explicit anchoring of non-translating mRNAs, but translation is automatically initiated when non-translating mRNA molecules encounter mitochondria and given a chance to anchor in the same time step as well. This follow-up study is agnostic about whether the anchoring mechanism of mRNA which approach mitochondria in a not-yet-translating state would be via the mRNA itself or the effect of an increased probability of translation. However we note that in the former case, whatever causes the mRNA to anchor to mitochondria would somehow have to be specific for genes coding for mitochondrially localizing proteins.

### Non-translating mRNA must have much higher mobility

The second conclusion is that the effective diffusion coefficient of mRNA must be significantly higher (20–100x) before initiating translation than afterwards. To compare this conclusion with the initial study, the difference between how mRNA mobility was modeled in that study (four total parameters: a diffusion coefficient and an entrapment probability for non-translating and translating mRNAs, respectively) and this follow-up study (where mobility effects are bundled into an effective diffusion coefficient) must be considered. Since the effective diffusion of the follow-up study conceptually includes possible periods of reduced mobility, it would be expected to be numerically smaller than the diffusion coefficient in the initial study. Also, since in the initial simulation study translating mRNAs had a much higher probability of entrapment than non-translating mRNAs, the ratio (non-translating versus translating) of effective diffusion constant in this follow-up study should be greater than ratio used in the initial simulation study. This is indeed the case for the effective diffusion coefficients with good fit to the data. For example (in units of *µ*m^2^/s) Params1 and Params2 have (non-translating: translating) effective diffusion coefficients of (0.0015625 : 7.8125e-05) and (0.00625 : 6.25e-05) respectively, while the diffusion coefficients used for the initial study were (0.123 : 0.03) for non-translating and translating mRNA respectively.

## References

[1] Thomas M. Poulsen, Kenichiro Imai, Martin C. Frith, and Paul Horton. Hallmarks of slow translation initiation revealed in mitochondrially localizing mRNA sequences. 2019. bioRxiv: 614255.

[2] Premal Shah, Yang Ding, Malwina Niemczyk, Grzegorz Kudla, and Joshua B. Plotkin. Rate-limiting steps in yeast protein translation. Cell, 153(7):1589–1601, 2013.

[3] Marlena Siwiak and Piotr Zielenkiewicz. A comprehensive, quantitative, and genome-wide model of translation. PLoS Comput. Biol., 6(7):1–15, 2010.

[4] Julien Sylvestre, Stéphane Vialette, Marisol Corral-Debrinski, and Claude Jacq. Long mRNAs coding for yeast mitochondrial proteins of prokaryotic origin preferentially localize to the vicinity of mitochondria. Genome Biol., 4(7):R44, 2003.

[5] Christopher C. Williams, Calvin H. Jan, and Jonathan S. Weissman. Targeting and plasticity of mitochondrial proteins revealed by proximity-specific ribosome profiling. Science, 7(346):748–751, 2014.

