## Supplementary material for "A Simulation Study of mRNA Diffusion and Mitochondrial Localization": Document source and data files: perl5tab

#!/usr/bin/perl
#  Author: Paul Horton
#  Copyright (C) 2016, 2017, 2018  Paul B. Horton, All rights reserved.
#
#  License: Anyone may use this software under the GNU Public License GPLv3.
#  For discussion of other licenses please contact the author, Paul Horton.
#
#  Description: an extenstion of perl autosplit mode.
#               Designed to ease one-liner handling of tab separated fields data streams.
#
# ────────────────────  Variable Naming and Use Conventions  ────────────────────
#  Subroutines visible from user scripts should start with '_'
#  UNLESS they are intended to be available to users, e.g. ih().
#
use utf8::all;
use Carp;
use Encode;
use Hash::Util qw(lock_keys lock_keys_plus unlock_keys lock_hash);
use List::Util qw(first max maxstr min minstr reduce shuffle pairkeys pairmap pairvalues pairs);
use List::MoreUtils qw(:all);
use POSIX qw(ceil floor);
use Scalar::Util qw(looks_like_number dualvar);
use Scalar::Util::Numeric qw(isfloat isint isneg);
use Statistics::Basic;
use Statistics::Distributions;
use Statistics::RankCorrelation;
use Unicode::Collate;
use Data::Lock qw(dlock dunlock);

$VERSION= '0.002_001';  eval $VERSION;
use strict;
use warnings;
no warnings  'experimental::lexical_subs';
use feature qw(lexical_subs say state);


my %numRE_=  (
    plain   =>   qr/^[+-]?   (?:  [0-9]+ (?:[.][0-9]*)?  |  [.][0-9]+  )   (?:[eE][+-]?[0-9]+)?  $/x,

    InfOK   =>   qr/^[+-]?
      (?:
        (?:inf)  |
        (?:  [0-9]+ (?:[.][0-9]*)?  |  [.][0-9]+  )   (?:E[+-]?[0-9]+)?
      )$/ix,
    NanOK   =>   qr/^[+-]?
      (?:
        (?:nan)  |
        (?:  [0-9]+ (?:[.][0-9]*)?  |  [.][0-9]+  )   (?:E[+-]?[0-9]+)?
      )$/ix,
    InfNanOK=>   qr/^[+-]?
      (?: 
        (?:inf)  |
        (?:nan)  |
        (?:  [0-9]+ (?:[.][0-9]*)?  |  [.][0-9]+  )   (?:E[+-]?[0-9]+)?
      )$/ix
    );
lock_hash %numRE_;

my $numRE_;


#────────────────────   Some subs which do not depend on global vars  ────────────────────


# ────────────────────  Utilities  ────────────────────
sub Dumper{
    eval  'use Data::Dumper';
    goto &Data::Dumper::Dumper;
}


#  Return true iff $_[0] is a Reference to a List of Lists.
#  Loose in the sense that it accepts undef elements, such as [ [3,4], undef, ...]
sub _LoLRefP_loose{
    @_   or   _dieProg(  '_LoL_P called with no arguments'  );
    ref $_[0] eq 'ARRAY'   or   return undef;
    for(   @{ $_[0]}   ){
        return undef  if defined && ref ne 'ARRAY';
    }
    return 't';
}

# Return one minus the size of longest component list pointed to by $_[0]
sub _LoLRef_maxListSize{
    my $maxSize=  -1;
    for(   @{ $_[0]}   ){
        $maxSize=  @$_   if defined && @$_ > $maxSize;
    }
    return  $maxSize;
}


#  fillHoles( $fillVal, \@matrixWannabeLoL )
#  @matrixWannabe should be a list of lists.
#  fillHoles makes @matrixWannabe into a rectangular shape and fills any undef values with $fillVal.
#  If $fillVal is omitted missingMark() is used as the value to fill.
#  If there is demand, interface may be expanded to take multiple @matrixWannabeLoL
sub fillholes{  goto &fillHoles  }
sub fillHoles{
    @_   or   _diell(
        'fillHoles called with no arguments.'=>
        'fillHolesが引数無しで呼び出された。');
    my $fillVal=  missingMark();
    my $firstRefIdx=  firstidx {ref} @_;
    $firstRefIdx > -1   or   _diell(
        'fillHoles called without any array reference argument'=>
        'fillHolesの引数にarray参照子がなかった。');
    $firstRefIdx < +2   or   _diell(
        'fillHoles expects at most one non-reference argument but got '. 1+$firstRefIdx .'.'=>
        'fillHolesは多くとも１個の非参照子引数を受け付けるが、'. 1+$firstRefIdx .'を受け取った。');
    $fillVal=  shift   if $firstRefIdx == 1;

    _LoLRefP_loose( $_[0] )   or   _diell(
        'fillHoles expects any reference argument to refer to a list of lists, but argument number '. $firstRefIdx .'is a reference to something else.'=>
        'fillHolesの参照子引数はすべてリストへのリスト（「LoL」、例えば、[[3,4],[1,3,5]]）への参照子でなければならないが、第' .$firstRefIdx. 'の引数はLoLと異なる型のデータへの参照子。');

    my $maxListSize=   _LoLRef_maxListSize $_[0];
    for  my $listRf(  @{ $_[0]}  ){
        if(  defined $listRf ){
            $$listRf[$_] //= $fillVal   for 0..$maxListSize-1;
        }
        else{
            $listRf=  [($fillVal) x $maxListSize];
        }
    }
}#fillHoles


#  push2( @f, @g, @val )
#  Shortcut for  push @f, $val[0]; push @g, $val[1]
sub push2(\@\@;@){
    @_ == 4   or   _diell(
        'push2 expected 4 arguments, but got '.@_=>
        'push2の引数は４個あるはずだが、'.@_.'個を受け取った。');
    push @{$_[0]}, $_[2];
    push @{$_[1]}, $_[3];
}


#  push3( @f, @g, @h, @val )
#  Like push2, but with three arrays to copy into.
sub push3(\@\@\@;@){
    @_ == 6   or   _diell(
        'push3 expected 6 arguments, but got '.@_=>
        'push3の引数は６個あるはずだが、'.@_.'個を受け取った。');
    push @{$_[0]}, $_[3];
    push @{$_[1]}, $_[4];
    push @{$_[2]}, $_[5];
}


#  Like builtin ::push, but first argument should be an array reference.
sub pushref{
    push @{$_[0]}, @_[1..$#_];
}


#  List List::MoreUtils::part, but returns indices
sub partidx(&@){
    my $code=  shift;
    my @retVal;
    for  my $i(  0..$#_  ){
        $_=  $_[$i];
        my $part=  &$code;
        defined $part   or
            _diell("Working with element $i, partIdxs expected a non-negative integer partition number but got undef"=>
                   "${i}番目の要素の処理中に、非負整数の分割番号を期待したが、undefに出くわした。");
        $part =~ /\D/   and
            _diell("Working with element $i, partIdxs expected a non-negative integer partition number but got '$part'"=>
                   "${i}番目の要素の処理中に、非負整数の分割番号を期待したが、'${part}'に出くわした。");

        push @{$retVal[$part]}, $i;
    }
    return @retVal;
}


#Return true if @_ contains no empty strings
sub nomt{  all {length} @_  }

#Return non-empty strings in @_.
#named for the construct:   say @x  if an @x;
sub an{  grep {defined && length} @_  }


{ state $printfWantsNumRx=  qr{ % [-+.#0-9]* l? [defgoux] }x;

  #Return strings in @_ sprintf formated according to FORMAT.
  #When FORMAT expects a number, non-numbers are passed through as is.
  sub form{
      wantarray   or  goto &_formScalar;
      @_ > 1   or   _diell(  'form() expected at least two arguments but got '.@_=>'form()はふたつ以上の引数が必要であるが、'.@_.'個しか受け取らなかった。'  );
      my $format=  shift;
      if(  $format=~  /$printfWantsNumRx/  ){
          map  {looks_like_number $_?  sprintf $format, $_  :  $_}    @_;}
      else{
          map  {                       sprintf $format, $_       }    @_;}
  }

  #Version of form when called in scalar context.
  sub _formScalar{
      defined  wantarray   or
          _diell(  'form() called in a void context'  =>
                   'void文脈でform()が呼ばれた。'  );
      @_ == 2   or
          _diell(  'form() called in scalar context should have 2 arguments, but called with '.@_  =>
                   'スカラ文脈ではform()はふたつの引数で呼ばなければならないが、'.@_.'を受け取った。'  );

      my $format=  shift;
      if(  $format=~  /$printfWantsNumRx/  ){
          looks_like_number $_[0]?  sprintf $format, $_[0]  :  $_[0];
      }
      else{
          ;                         sprintf $format, $_[0];
      }
  }

  #Modify @_ sprintf formated according to FORMAT.
  #When FORMAT expects a number, non-numbers left unchanged.
  sub reform{
      @_ > 1   or   _diell(  'reform() expected at least two arguments but got '.@_=>'reform()はふたつ以上の引数が必要であるが、'.@_.'個しか受け取らなかった。'  );
      my $format=  shift;
      if(  $format=~  /$printfWantsNumRx/  ){
          $_=  looks_like_number $_?  sprintf $format, $_  :  $_   for @_;}
      else{
          $_=                         sprintf $format, $_          for @_;}
  }
}


#Like say, but does nothing if @STRING is empty or contains any empty strings or undef values.
sub speak{    say @_   if  @_  and  all {length} @_    }

#Like say, but replaces undef arguments with the missing value mark.
sub dit{    say   map  {$_ // missingMark()}  @_    }

#Defined and EQual.  If any arguments are undefined return undef.  Otherwise return 't' if all arguments are string equal and '' if not.
sub deq{
    defined $_[0]  or  return _diell(  'deq called with no arguments'=>
                                       'deqが引数無しで呼び出された。'  );
    for(  1..$#_  ){
        defined  $_[$_]   or  return undef;
        $_[0] eq $_[$_]   or  return '';
    }
    return 't';
}


sub numP(_);  #Declared here, but defined later after defining %opt_.

#doNum( SUB, val... ).
#If all vals look numerical, run SUB(val...) and forward its return value,
#otherwise return the missing value mark.
sub doNum(&@){
    my $sub=  shift;
    ref $sub eq 'CODE'  or  _diell(  'doNum expected code reference (i.e. sub{...}) for first argument',
                                     'doNumの最初の引数はコード参照(つまりsub{...})でなければならない。');
    @_   or   _diell(  "doNum expected 2 or more arguments, but got only one.  Should look like 'donum( sub{CODE}, VAL )'.",
                       "doNumはふたつ以上の引数が必要であるが、ひとつしか受け取らなかった。donumは'donum( sub{CODE}, VAL )'のように使う。");
    
    return missingMark()    if  any {!numP} @_;

    return &$sub( @_ )   for @_;   #To alias $_ to $_[0] in SUB.
}
sub donum(&@){  goto &doNum  }


#  bigger( STRING1, STRING2 )
#  Return true iff NUM1 would be a preferable value to NUM2 when looking for big values.
#  For special numbers the preference being  1. inf, 2. normal numbers, 3. nan, 4. -nan 5. -inf
sub bigger{
    defined $_[0]   and  $_[0]=~ /$numRE_/    or  return;
    defined $_[1]                             or  return 't';

    if(  $_[0]=~ /\d/  ){
        if(  $_[1]=~ /\d/      ){   return  $_[0] > $_[1]      }
        if(  $_[1]=~ /^inf$/i  ){   return                     }
        return  't';   # $_[1] ∈ {-inf, nan, -nan}
    }

    # $_[0] ∈ {inf, -inf, nan, -nan}
    if(  $_[1]=~ /\d/  ){
        if(  $_[0]=~ /^inf$/i  ){   return  't'                }
        return;  # $_[0] ∈ {-inf, nan, -nan}
    }

    # Both $_[0] and $_[1] are ∈ {inf, -inf, nan, -nan}.
    if(  $_[0]=~ /^inf$/i  )     {   return   $_[1]!~ /^inf$/i   }
    if(  $_[0]=~ /^nan$/i  )     {   return(  $_[1]=~ /^-inf$/i  or  $_[1]=~ /^-nan$/i  )}
    if(  $_[0]=~ /^-nan$/i )     {   return   $_[1]=~ /^-inf$/i  }
    return;   # $_[0] is -inf
}#bigger


#  lesser( STRING1, STRING2 )
#  Return true iff NUM1 would be a preferable value to NUM2 when looking for small values.
#  For special numbers the preference being  1. inf, 2. normal numbers, 3. nan, 4. -nan 5. -inf
sub lesser($$){
    defined $_[0]   and  $_[0]=~ /$numRE_/    or  return;
    defined $_[1]                             or  return 't';

    if(  $_[0]=~ /\d/  ){
        if(  $_[1]=~ /\d/      ){   return  $_[0] < $_[1]     }
        if(  $_[1]=~ /^-inf/   ){   return                    }
        return  't';   # $_[1] ∈ {-inf, nan, -nan}
    }

    # $_[0] ∈ {inf, -inf, nan, -nan}
    if(  $_[1]=~ /\d/  ){
        if(  $_[0]=~ /^-inf$/i ){   return  't'               }
        return;  # $_[0] ∈ {inf, nan, -nan}
    }

    # Both $_[0] and $_[1] are ∈ {inf, -inf, nan, -nan}.
    if(  $_[0]=~ /^-inf$/i )    {   return  $_[1]!~ /^-inf$/i  }
    if(  $_[0]=~ /^-nan$/i )    {   return( $_[1]=~ /^inf$/i  or  $_[1]=~ /^nan$/i  )}
    if(  $_[0]=~ /^nan$/i  )    {   return  $_[1]=~ /^inf$/i   }
    return;   # $_[0] is inf
}#lesser


#Set first argument to maximum value of all arguments, including itself.
#undef values are treated as less than any defined value.
#defined values which do not look like numbers are skipped.
sub bemax{
    for (@_[1..$#_]){
        $_[0]= $_    if  bigger $_, $_[0];
    }
}

#Like bemax.
sub bemin{
    for (@_[1..$#_]){
        $_[0]= $_    if  lesser $_, $_[0];
    }
}


#Like bemax.
sub beminmax{
    for (@_[2..$#_]){
        $_[0]= $_    if  lesser $_, $_[0];
        $_[1]= $_    if  bigger $_, $_[1];
    }
}


#SNset( %S, PATHNAME )
#Construct hash set %S to contain lines of PATH mapped to serial numbers as its elements.
#Values of elements are set to 1.. in their order in PATH, repeats overwriting.
sub SNset(  \% $  ){
    my $path=  $_[1];

    my @item;
    if(   $path eq '-'  or  $path =~ /^stdin$/i   ){
        chomp(  @item= <STDIN>  );}
    else{
        -f $path   or   _diell(  "in SNset; could not find file '$path'",
                                 "SNset実行中でファイル'${path}'が見つからなかった。"  );
        open  my $in, '<', $path   or   _diell(  "in set; failed to open file '$path'.  $!",
                                                 "set実行中でファイル'${path}'が開けなかった。"  .  decode_utf8 $!);
        chomp(  @item= <$in>    );
    }
    my $c;
    $_[0]->{$_}= ++$c  for @item;
}

sub  snset{  goto &SNset  };


# ──────────────────────────────  Global Variables  ──────────────────────────────
#  Variables intentionally visible to user
my @F;           #Fields of the current row, subject to ditto substitution under the -D|--ditto option..
my @Fin;         #F of the current row as read in.
my @FinPrev;     #Fields of previously read row as input.
my @H;           #To hold head fields unless $opt_{noHeadP}.


{ state $NR=  state $NRT=  0;
  sub NR (){  $NR  }
  sub NRT(){  $NRT }
  sub _incrNR(){   ++$NR   }
  sub _incrNRs {   ++$NR;   ++$NRT  if $_[0]   }
  sub _rezeroNRs(){  $NR= $NRT= 0  }
}


#  Print $msg and usage info to STDERR and STDOUT.
#  Or just to STDERR, if both STDOUT and STDERR are directed to the same inode.
sub _dieUsage{
    my $msg=  _ll(@_);
    $msg =  _ll('perltab command line parsing error; ', 'perltabコマンド行処理エラー、') . $msg . "\n";
    $msg.=      "Usage: perltab [opts...] tabSeparatedFieldsInputFile...\n       perltab -h";

    say STDERR $msg;
    say STDOUT $msg    if  (stat STDERR)[0] != (stat STDOUT)[0]  or  (stat STDERR)[1] != (stat STDOUT)[1];

    exit 64;
}


{ my $compilingScript_;
  my $scriptDesc_;

  #  Try compiling script after doctoring with perltab shortcuts.  MODIFIES FIRST ARG.
  #  Regarding placement of this subroutine within this file,
  #  USER GLOBALS SHOULD BE IN SCOPE, to avoid spurious warnings such as 'Possible unintended interpolation... of @F'
  sub doctorScript{
      ($scriptDesc_, my $msgStart_)=  @_[1,2];

      for(  $_[0]  ){
          s{ ^\s* | \s*$ }  {}gx;  #Remove leading and trailing white space.
          s{\bF\(  ( [^"',(){}\$@]+ )  \)}  {F(qw($1))}gx;
          s{\bN\(  ( [^"',(){}\$@]+ )  \)}  {N(qw($1))}gx;
          $_   or   $_=  q{0;};   #Make sure all defined values of %script_ look true in boolean context.
      }
      $compilingScript_= $_[0];
      local $SIG{__WARN__}=  local $SIG{__DIE__}=  \&compileErrorHandler;
      eval  "no strict 'vars';  no indirect;  use warnings;  sub{ $compilingScript_ }";
      substr( $compilingScript_, 1024 )= '...'   if length $compilingScript_ > 1024;
      _dieUsage $msgStart_._ll(" but the string '$compilingScript_' produced error '$@' when compiling."=>
                               "文字列'${compilingScript_}'のコンパイル時にエラー:\n" . decode_utf8($@) . 'が発生した。')
          if $@;
  }


  #Print string in @_ to STDERR and also STDOUT unless both are connected to a terminal.
  sub _say2{
      for(  @_  ){
          say STDERR;
          say STDOUT "STDOUT: $_"   if  (stat STDERR)[0] != (stat STDOUT)[0]  or  (stat STDERR)[1] != (stat STDOUT)[1];
      }
  }


  #  Use eval to test if $script appears to be well-formed Perl.
  #  Die with warning based on error message $errMsg if it fails.
  #  $scriptDesc_ is used for diagnostic messages.
  sub compileErrorHandler{
      my $errMsg=  $_[0];

      (my $lineNum)=  $errMsg=~/ at \(eval \d+\) line (\d+)/;
      my @scriptLine=  split "\n", $compilingScript_;


      #  ──────────  Print information about error  ──────────
      if(  defined $lineNum  ){
          my $scriptLine  =  $scriptLine[$lineNum-1];

          $errMsg=~  s| \(eval \d+\)||;
          if(  @scriptLine < 2  ){
              _say2 _ll("Trouble compiling $scriptDesc_ '$compilingScript_'. ",
                        "${scriptDesc_}スクリプト'${compilingScript_}'にコンパイル・エラー。")
                  . $errMsg;
              exit 64;                                                          #PROGRAM EXIT POINT
          }

          _say2 _ll("${errMsg}\n\nWhen compiling ${scriptDesc_} script, at line ${lineNum}.",
                    "${errMsg}\n\n${scriptDesc_}スクリプト、コンパイル時、${lineNum}行目。");

          #  ─────  Print part of script near error  ─────
          my $begLineNo  =  max        0     , $lineNum-7;
          my $endLineNo  =  min  $#scriptLine, $lineNum+7;

          $begLineNo == 0   or   say  '    ...';

          _say2(  $_ == $lineNum-1?   #Count from zero.
                  "--> $scriptLine[$_]"  :
                  "    $scriptLine[$_]"  )    for  $begLineNo .. $endLineNo;

          $endLineNo == $#scriptLine   or   say  '    ...';
      }
      else{   #No line number in $errMsg.
          _say2 _ll("perltab error encountered when compiling $scriptDesc_ script:",
                    "perltab ${scriptDesc_}スクリプトのコンパイル時にエラーが発生した。")
              ,substr $errMsg, 0, 1024;
      }

      exit 64;
  }#compileErrorHandler

}


my %script_;
{# ───────────────  Run scripts and manage associated state  ───────────────
  my $scriptType_;  #Current script type.

  sub _runScript{
      $script_{ $_[0]}   or   return;
      $scriptType_=  shift;

      local $SIG{__DIE__ }=  local $SIG{__WARN__}=  \&_runtimeErrorHandler;

      no strict 'vars';  no indirect;  no warnings qw(once redefine reserved);  use warnings FATAL => qw(printf uninitialized);
      my @retVal=  eval $script_{$scriptType_};
      $@   and   _diell("Error in $scriptType_ script: $@"=>"${scriptType_}スクリプトにエラー、$@");
      return  wantarray?  @retVal  :  $retVal[0];
  }


  #  Called when $scriptType_ script has generated a runtime exception.
  #  Print information in error message $errMsg, part of the script and then exit.
  sub _runtimeErrorHandler{
      chomp(  my $errMsg=  $_[0]  );

      (my $lineNum)=  $errMsg=~/ at \(eval \d+\) line (\d+)/;
      my @scriptLine=  split "\n", $script_{$scriptType_};

      if(  $scriptType_=~ /ROW/  ){
          $errMsg=  _ll('While reading second pass -in2 input,' => '-in2で指定された入力の読み込み時、') . $errMsg
              if $script_{ROW2};
          $errMsg=  _lln("While reading input row number ${\NR}," => "${\NR}目の行を処理しようとした際、") . $errMsg;
      }

      #  ──────────  Print information about error  ──────────
      if(  defined $lineNum  ){
          dlock my $scriptLine=  $scriptLine[$lineNum-1];

          if(   my ($num1,$num2)=  ($errMsg =~ /Number found where operator expected at.*"(\d+) (\d+)"/)   ){
              $errMsg .= _ll("\nTry replacing '$num1 $num2' with '$num1, $num2'?",
                             "\n'$num1 ${num2}'を'$num1, ${num2}'に修正すると良いかも知れない。");
          }
          
          $errMsg=~  s| \(eval \d+\)||;
          if(  @scriptLine < 2  ){
              $errMsg= _ll("Trouble running $scriptType_ script '$script_{$scriptType_}'.  ",
                           "${scriptType_}スクリプト'$script_{$scriptType_}'の実行エラー。") . $errMsg;
              _say2 $errMsg;   exit 64;                                       #PROGRAM EXIT POINT
          }

          _say2 _ll("${errMsg}\n\n${scriptType_} script, line ${lineNum}.",
                    "${errMsg}\n\n${scriptType_}スクリプト、${lineNum}行目。");

          #  ─────  Print part of script near error  ─────
          my $begLineNo  =  max        0     , $lineNum-7;
          my $endLineNo  =  min  $#scriptLine, $lineNum+7;

          $begLineNo == 0   or   say  '    ...';

          _say2(  $_ == $lineNum-1?   #Count from zero.
                  "--> $scriptLine[$_]"  :
                  "    $scriptLine[$_]"  )    for  $begLineNo .. $endLineNo;

          $endLineNo == $#scriptLine   or   _say2  '    ...';
      }
      else{   #No line number in $errMsg.
          _say2  _ll("perltab error encountered when executing $scriptType_ script:",
                     "perltab ${scriptType_}スクリプトを実行中にエラーが発生した。")
              ,substr $errMsg, 0, 1024;
      }

      exit -1;
  }#_runtimeErrorHandler

}# ───────────────  Run scripts and manage associated state  ───────────────


{#──────────   Message Language Selection Related  ──────────

  state $langI=  0;
  $langI=  1   if $ENV{LANG} =~ /^j[ap].*[.]utf-?8$/i;

  sub _lang(){  qw(ENG JPN)[$langI]  }

  #Select string based on output language.
  sub _ll {  my $msg= $_[$langI] // $_[0];  $msg       }
  sub _lln{  my $msg= $_[$langI] // $_[0];  $msg."\n"  }

  sub _lln_concat{
    # Receive flattened list (msgLang1, msgLang2,... extraMsgLang1, extraMsgLang2,...)
    # Concatenate newline, then extra message modifying @_
    @_ % 2   and   die  'Programming error, expected even length list.';

    my @half2=  splice @_, @_/2;
    $_[$_] .=  "\n" . $half2[$_]   for 0..$#_
  }

}


#  Substitute the string 'undef' for undef values in @_.
sub _stringifyUndef{   defined   or  $_= 'undef'   for @_   }


#Tried using state sub dieProg{...},  but that crashes on v5.18.2 and v5.20.2.
sub _dieProg{   say STDERR 'perltab internal programming error: ', @_;  exit 1   }

my $callerForDieMsg_;
sub _diell{
    if(  grep /CALLER/, @_  ){
        length $callerForDieMsg_   or   _dieProg  '$callerForDieMsg_ was on defined when _diell called with arguments ', @_;
        map  {s/CALLER/$callerForDieMsg_/g}  @_;
    }
    die  _ll(@_) . "\n"
}


{#────────────────────  Help Doc Related Code   ────────────────────

  #Return language of line $_[0].
  state sub lineLang{
      return 'JPN'    if  $_[0]  =~  /(\p{Hiragana}|\p{Katakana})/;
      return '---'    if  $_[0]  =~  /^  [^-]/ ;                   #Lines starting with two spaces (followed by anything but: -, for options table).
      return 'ENG'    if  $_[0]  =~  /   [a-zA-Z]+.*   [.,:] $/x;  #Lines ending in {.,:} considered English.
      return 'ENG'    if  $_[0]  =~  /^\S[a-zA-Z]+.*       ; $/x;  #Or ending in ';', if flush with the left margin.
      return 'ENG'    if  $_[0]  =~  /   [a-zA-Z]+.* [.] [)] $/x;  #Lines ending in  ).   considered English.
      return '---';  #verbatim line.
  }


  my %wrappedTab;   #  $wrappedTab{TEXT} == Size of left hand pad to add to lines 2,3...of TEXT.


  state sub sayDocLines{
      @_ == 1   or   _dieProg  'wrong number of args';
      chomp(  my $docLines=  shift  );

      state $termWidth=  `tput cols`;

      state $text;   state $leftPad;   state $lang;
      state $wRE=  qr{(?:  \p{Han} | \p{inHiragana} | \p{inKatakana} | \p{InCJKSymbolsAndPunctuation} | \p{InHalfwidth_and_Fullwidth_Forms}   )}x;

      #BEG──────────  sayDocLines internal subs  ──────────

      my sub width($){   #Return display width of $_[0]
          my $w=()=  $_[0]=~ /($wRE)/g;       
          return    $w  +  length $_[0];
      }

      #Return max position in $text such that
      #the proportional font display width of substr( $text, 0, POS) is ≦ $width
      my sub textPosAtWidth{
          dlock my( $width  )=  @_;
          die  "invalid negative width: '$width'"   if $width < 0;
          length $text   or   return 0;

          my( $strPos, $disPos )=  (0,0);
          while(   $strPos < length $text   and   $disPos < $width   ){
              $disPos+=  substr( $text,$strPos,1 )=~  $wRE;
              $disPos++;   $strPos++;
          }
          $strPos--     if  $disPos > $width;   #While loop overran by one char position?
          return  $strPos;
      }


      #Parameters for heuristic pseudo-semantic line breaking.
      my %breakRE;  my %breakScore;

      my @breakers=(
          'space1',     qr/(\s)/,          => 1.00,
          'commaSpace', qr/(\s)(?=,)/      => 1.05,
          'space2',     qr/(\s\s+)/        => 1.25,
          'space3',     qr/(\s\s\s+)/      => 1.30,
          'fullStop',   qr/(\s\s+)(?=[.])/ => 1.30,
          );

      my $it= natatime 3, @breakers;
      while(  my @val= $it->()  ){
          my $patName=  shift @val;
          $breakRE   {$patName}=  shift @val;
          $breakScore{$patName}=  shift @val;
      }


      #Return position of next recommended line break in $text, of undef if none found.
      my sub bestBreak{
          if(  $lang eq 'JPN'  ){
              width $text > $termWidth-2   or   return  width $text;

              my $breakPos=  textPosAtWidth $termWidth;
              --$breakPos  while
                  $breakPos > 0
                  and substr( $text, $breakPos, 1   )=~ qr/[])、。]/                       #Bad line start
                  ||  substr( $text, $breakPos-1, 1 )=~ qr/[-[(（「参一\#=_a-zA-Z0-9\$@]/  #Bad line end
                  ;
              return $breakPos;
          }
          else{
              length $text > $termWidth-1   or   return  length $text;
              dlock my $leftPadWidth=  length $leftPad;
              dlock my $widthSansPad=  $termWidth- $leftPadWidth;
              my $rtext=    reverse   substr $text, $leftPadWidth, $widthSansPad;
              
              #Break more in the middle if the following line will be short.
              my $shortEnough=   min  $widthSansPad/2,  length($text) - $widthSansPad;

              my $curBestScore= -1;   my $curBestBreak=  undef;
              for  my $patName  (keys %breakRE){
                  if(  $rtext=~  $breakRE{$patName}  ){
                      my $break=  $termWidth - $+[0];
                      next    if $break < 12;
                      my $score=   min($break, $shortEnough) * $breakScore{$patName};
                      ($curBestBreak, $curBestScore)=  ($break, $score)    if $score > $curBestScore;
                  }
              }
              return  $curBestBreak;
          }
      }#bestBreak.
      #END──────────  sayDocLines internal subs  ──────────


      #Do simple case first to avoid edge case of split (empty string).
      if(   width($docLines) < $termWidth  ){
          say $docLines;  return;                             #EXIT sayDocLines
      }

    DOC_LINE:
      for(  split /\n/, $docLines, -1  ){
          $text= $_;
          $lang=  lineLang $text;
          if(  width $text  >  $termWidth
               and
               #Only wrap verbatim lines in a tabular environment.
               $lang ne '---' ||  exists $wrappedTab{$text}
              ){
              ($leftPad)=  exists $wrappedTab{$text}?  $wrappedTab{$text}  :  $text=~ /\A(\s*)/;
              width $leftPad  < $termWidth   or   _dieProg  'In doc, leading white space too long';

              #Output initial line.
              defined(  my $break=  bestBreak  )   or   say($text),  next DOC_LINE;
              say  substr $text, 0, $break;
              substr $text, 0, $break, '';

              #Output subsequence lines, left padded with $leftPad;
              while(  length $text  ){
                  $text=~ s/\A(\s*)/$leftPad/;
                  defined(  $break=  bestBreak  )   or  say($text), last;
                  say  substr $text, 0, $break;
                  substr $text, 0, $break, '';
              }
          }
          else{
              say $text;
          }
      }#for docLines.
  }#sayDocLines


  my @doc;  #Documentation lines read from <DATA>, filtered for language.
  my @toc;  #To hold documentation table of contents.
  my %kw_section;  #Map keyword to section.


  #Parse documentation in DATA at bottom of this file, to fill in @doc, %kw_section.
  state sub parseDoc(){
      my %kw_lineNum;  #Map keywords to their documentation starting line number.
      my %kw_openP;  #True for keyword specs which have been opened by not closed.
      my $xtermP=  $ENV{TERM} eq 'xterm';
      my $B=  "\e[1m";   #Start  bold
      my $b=  "\e[0m";   #Reset (bold)

      my(  $curLine, $prevLine, $curHead  )=  '';
      my $lineNum=  my $prevSec_lineNum=  0;

      my $curTab= undef;

      my sub dieDoc{
          say STDERR  "In documentation line #$lineNum: $curLine";
          say STDERR  @_;
          exit 1;
      };

    LINE:
      while(   my $line=  <DATA>   ){
          next LINE                        if  $line=~ /^#/;
          chomp $line;

          $prevLine= $curLine;  $curLine= $line;

          if(  $line=~ /^(=settab +)==/  ){
              defined $curTab  and   dieDoc  'tab already set';
              $curTab=  $+[1];                                                           next LINE;
          }
          if(  $line=~ /^=unsettab/  ){
              defined $curTab   or   dieDoc  'tab already unset';
              $curTab=  undef;                                                           next LINE;
          }
          if(   (my $kwSpec)=  $line=~ /^=kw\s*(.*)$/   ){
              length $kwSpec   or   dieDoc  '=kw line with empty keyword list';
              for  my $kw  (split '\|', $kwSpec){
                  exists $kw_openP{$kw}    and   dieDoc  "Multiple starts for keyword '$kw'";
                  $kw_openP{$kw}= 't';
                  $kw_lineNum{$kw}= $lineNum;
              }
              ;                                                                          next LINE;
          }

          if(   (my $kwSpec)=   $line=~ /^=wk\s*(.*)/   ){
              length $kwSpec   or   dieDoc  '=wk line with empty keyword list';
              for  my $kw  (split '\|', $kwSpec){
                  $kw_openP{$kw}   or   dieDoc  "keyword '$kw' has end before any start";
                  delete $kw_openP{$kw};
                  $kw_section{$kw} .=   join  "\n",  @doc[ $kw_lineNum{$kw} .. $lineNum-1 ], '', '';
              }
              ;                                                                          next LINE;
          }

          if(   (my $nextHead)  =  $line =~ /^=head1\s+(.+)/   ){
              #When in 'ENG', hide all but English.
              $nextHead  =~  s/ \| .* $//x    if _lang eq 'ENG';

              if(  $curHead  ){   #Process record which ended one line ago.
                  push  @toc,   "    $curHead";
                  #head1 headings are automatically treated as keywords.
                  for  my $kw  (split '\|', $curHead){
                      $kw_section{$kw}=   join  "\n",  @doc[$prevSec_lineNum..$lineNum-1];
                  }
              }

              if(  $nextHead =~ /END_DOC/   ){
                  dieDoc  "Documentation for keyword '$_' opened but never closed"   for keys %kw_openP;
                  return;                                                          #SUBROUTINE EXIT
              }

              $curHead=  $nextHead
                  or   dieDoc  "Head '$curHead' evaluates to false, which will confuse parsing.";

              $prevSec_lineNum=  $lineNum;
              $line=  $xtermP?  "$B$curHead$b" : $curHead;
              $line=  '        ' . $line;
          }#=head1
          elsif(   (my $head2)=  $line =~ /^=head2\s+(.+)$/   ){
              $line=   "\n    " . _ll(  split '\|', $head2  );
          }
          else{#  Not a =head line
              #If $line starts with ditto mark; substitute portion of previous line into it.
              if(  (my $dittoPrefix)=  $line =~ /^( \| \s*)/x   ){
                  my $dittoLen=  length $dittoPrefix;
                  dieDoc  "dittoLen:$dittoLen longer than previous line"    if $dittoLen > length $prevLine;
                  substr  $line,  0,  $dittoLen,  substr $prevLine, 0, $dittoLen;
              }
              #Skip lines not appropriate for _lang language.
              $_ eq _lang  or  $_ eq '---'  or  next LINE    for lineLang $line;        #LOOP FLOW
          }#  if =head1  else =head2...
          
          $lineNum++;
          push @doc, $line;
          $wrappedTab{$line}=  (' ' x $curTab)   if defined $curTab;

      }#while  $line = shift @data

      dieDoc  'Documentation data did not end in "=head1  END_DOC"?';
  }#parseDoc

  #Process help option.
  sub _printHelp_thenExit{
      `tput cols` >= 70
          or   die _lln('To use the help facility, please widen your terminal to at least 70 chars.',
                        'ヘルプ機能を使うのに端末の幅を70(つまり漢字35字分)以上にしてください。');
      parseDoc;
      my $qry=   shift  //  'help';

      if(   $qry=~ /^(all|man)$/   ){
          sayDocLines $_   for @doc;                                    exit 0;

      }
      if(   $qry=~ /^(kw)$/i       ){
          my @kw=  keys %kw_section;
          @kw=  grep {/^-?[[:ascii:]]+$/} @kw   if _lang eq 'ENG';
          say  for sort @kw;                                            exit 0;
      }
      if(   $qry=~ /^(toc)$/i      ){
          say  for @toc;                                                exit 0;
      }
      for  my $prefRx  (qr/^$qry$/, qr/(?i:^$qry$)/, qr/^$qry/, qr/(?i:^$qry)/){
          my @match=  grep /$prefRx/, sort keys %kw_section   or   next;
          if(  @match == 1  ){
              my $kw=  $match[0];
              sayDocLines _ll("Documentation for '$kw'", "'${kw}'についての説明");
              sayDocLines  $kw_section{$kw};                             exit 0;  
          }
          say _ll("No documentation for '$qry'.  Perhaps you were looking for on of the following keywords?"=>
                  "${qry}についての説明文はない。以下のキーワードの説明文を求めているのでしょうか？");
          say  'candidates are:';   say  for @match;                    exit 0;
      }
      say _ll("Sorry, no help available for '$qry', try 'perltab -h kw for list of keywords",
              "'${qry}'についての説明がありません。'perltab -h kw'でキーワードのリストは表示できる。");
      ;                                                                 exit 0;
  }#_printHelp_thenExit
}#Documentation


# ━━━━━━━━━━  BEG: Assign and check command line options  ━━━━━━━━━━
@ARGV   or   _dieUsage( 'expected options and arguments on command line, but got none.',
                        'コマンド行にオプションと引数を期待したが、何もない。');

my %opt_;   #To map option names to values, obtained either via command line or top of script file.

{ my %optSpecs=  qw(
BEGIN             b|begin=*
BEGIN2           b2|begin2=*
countP            c|count
DATA              d|data-script=*
DATA2            d2|data-script2=*
defaultDittoMarkP D|ditto
dittoMark           ditto-mark=
extraFieldsOK     X|extra-fields-OK
ROWS              e|rows-script=*
ROWS2            e2|rows-script2=*
scriptFile        f|script-file=
inputFieldSep     F|field-sep=
grepP             g|grep
help             h!|help:HELP
HEAD              H|head-script=*
HEAD2            H2|head-script2=*
skipInfP            skipinf
missingMark       m|missing-mark=
modules           M|module=
skipNanP            skipnan
noHeadP           N|no-head
outputFile        o|output-file=
outputFieldSep   OF|output-field-sep=
printEachRowP     p|print-each-row
printScriptP      n|print-script
SORT_KEY          s|sort=*
SORT_KEY2        s2|sort2=*
grepQuietP        q|grep-quiet
testP             test
usageP            usage
printVersionP     V|version
END               z|end=*
END2             z2|end2=*
);
#h|help, s|sort are handled in a customized way.
#Reserved for possible future implementation
#inPlaceP         i|in-place   like sed -i
  lock_hash  %optSpecs;
  lock_keys_plus  %opt_,  keys %optSpecs;

  my(  $bundleFlag, $bundleOpt, %default, %needVal_optS, %scriptOptS, %toOptName   );

  #Extract option specification from $optSpecs and set %opt;
  while(   my($optName, $spec)=  each %optSpecs   ){

      $opt_{$optName} //= undef;   #Ensure that %opt_ has an entry for all options.

      if(     $spec =~ s/=[*]/=/   ){     $scriptOptS{$optName}= undef   }

      if(     $spec =~ s/=//       ){   $needVal_optS{$optName}= undef   }
      elsif(  $spec =~ s/:(\S*)//  ){        $default{$optName}=  $1     }

      for my $alias(  split '\|', $spec  ){
          if(   $alias !~ s/!$//  and  length $alias == 1   ){   #spec 'h!|help:HELP', is not bundle-able
              $bundleOpt  .= $alias;
              $bundleFlag .= $alias    unless exists $needVal_optS{$optName}
          }
          exists $toOptName{$alias}   and   _dieProg  "tried to overwrite existing option alias '$alias'";
          $toOptName{$alias}=  $optName;
      }
  }#WHILE  ($optName, $spec)=  each %optSpec
  dlock $bundleFlag;  dlock $bundleOpt;  lock_hash %default;  lock_hash %needVal_optS;  lock_hash %toOptName;  lock_hash %scriptOptS;


  #  Set scripts and return information to prepend to @ARGV from contents of file with pathname passed as $_[0].
  my sub parseScriptFile{
      $opt_{scriptFile}=  shift;
      $opt_{scriptFile}=~ /[.][ct]sv$/
          and   _dieUsage("filename '$opt_{scriptFile}' looks like a filename of input data file, but is positioned as a script filename."=>
                          "ファイル名'$opt_{scriptFile}'はスクリプト・ファイル名の位置にあるが、入力データファイルのファイル名に見える。");

      -e $opt_{scriptFile}   or   _dieUsage(  "Script file '$opt_{scriptFile}' does not exist."=>
                                              "スクリプト・ファイル'$opt_{scriptFile}'は存在しない。"  );
      -f $opt_{scriptFile}   or   _dieUsage(  "Script file '$opt_{scriptFile}' is not a plain file."=>
                                              "スクリプト・ファイル'$opt_{scriptFile}'は普通のファイルではない。"  );
      -r $opt_{scriptFile}   or   _dieUsage(  "Script file '$opt_{scriptFile}' is not readable."=>
                                              "スクリプト・ファイル'$opt_{scriptFile}'の読込み権限がない。"  );
      open  my $scriptFile, '<', $opt_{scriptFile};
      my $scripts=  do{ local $/; <$scriptFile> };
      my @ARGVfromFile;
      @ARGVfromFile=  split ' ', $1    if  $scripts=~ /\A#perltab(.*)/;

      require Text::Balanced;
      for  my $scriptType(  qw(BEGIN BEGIN2 HEAD DATA DATA2 ROWS ROWS2 END SORT_KEY SORT_KEY2)  ){
          if(my(               $upToBlockLabel,          $rest        )=
             (  $scripts=~  / ^( .*? ) $scriptType \s* ({ .* ) /xs   )
              ){

              $scripts=  $upToBlockLabel . $rest
                  if   my $block=  Text::Balanced::extract_codeblock( $rest, '{}' );
        
              _dieUsage("script only allowed one $scriptType block"=>
                        "スクリプトに${$scriptType}ブロックが重複している。")
                  if   $scripts=~  / $scriptType \s* { /x;

              doctorScript( $block, "BLOCK $scriptType",
                            _ll("When parsing script file '$opt_{scriptFile}' for $scriptType script,"=>
                                "スクリプトファイル'$opt_{scriptFile}'から${scriptType}スクリプトとして抽出した時、"));

              $opt_{$scriptType}=  $block;
          }
      }
      return @ARGVfromFile;
  }#my sub parseScriptFile


  my $ARGVforPrintScript= '';
  my $OPT_VALseparator= ' ';
  my $nextOPT=  sub{  @ARGV   or   return '';
                      do{   shift @ARGV; return ''   }if  $ARGV[0] eq '--';
                      do{   return    shift @ARGV    }if  $ARGV[0]=~ /\A-/;
                      return ''};

  while(   my $arg=  &$nextOPT()   ){
      if(   $arg=~  /^( -h|--help )$/x                                      ){
          _printHelp_thenExit( @ARGV )                                  ;next;}

      if(   $arg=~  /^( -f|--script-file )$/x  ){
          my $pathname=  shift @ARGV   or
              _dieUsage(  "option '${arg}' needs a following argument but none found."=>
                          "オプション'${arg}'の後に来るはずの引数が見つからない。"  );
          unshift @ARGV, parseScriptFile $pathname                     ;next;}
      
      #Ditto and another flag bundled together
      if(   $arg=~  /^-D[des]/                                              ){
          $opt_{dittoMark}= '';
          unshift @ARGV,  '-' . substr $arg, 2                          ;next;}

      state %seenValS;
      if(   $arg=~  /^(-[^=]+)  =  (.+)$/x                                  ){
          $OPT_VALseparator=  '=';
          unshift @ARGV, $1, $2;  $seenValS{$2}= undef                  ;next;}

      if(   $arg=~  /^(-[$bundleFlag]+) ([bdes]2)$/x                        ){
          unshift @ARGV,  $1, "-$2"                                     ;next;}

      if(   $arg=~  /^-([$bundleFlag]+ [$bundleOpt])$/x                     ){
          unshift @ARGV,  map {"-$_"} split //,$1                       ;next;}

      my $optStem=  $arg=~ s/^--?//r;
      if(   $arg=~ /^-s|^--sort/   ){
          ($optStem, my $sortSpec)=  $optStem=~ /^ (sort2?|s2?) (.*) /x;
          _parseSortSpecsStr(   $arg,  $sortSpec,  $optStem=~ /(2?)$/   );
          $ARGVforPrintScript.= " $arg";
      }
      if(   !exists $toOptName{$optStem}   ){
          if(   $arg  eq  '-in2'  ){
              _dieUsage(  "'-in2' encountered before any input file for the first pass.",
                          "'-in2'の前に一回目の入力読込みファイルが見当たらない。");
          }else{
              if(   $arg=~ /^-de$/  or  $arg=~ /^-ed$/   ){
                  _dieUsage(  "Only one of '-d' and '-e' script options can be given at once.",
                              "'-d'と'-e'のスクリプトオプションを同時に指定することはできない。");}
              else{
                  _dieUsage(  "there is no option '$arg'.",
                              "'${arg}'というオプションは存在しない。");
              }
          }
      }
      my $optName=   $toOptName{$optStem};
      if(   exists $needVal_optS{$optName}   ){
          my $val=   shift @ARGV //  _dieUsage(  "No value given for option '$arg'"=>
                                                 "オプション'${arg}'の値が指定されていない。");
          if(  exists $scriptOptS{$optName}  ){
              if(  $opt_{scriptFile}  ){
                  _dieUsage(  "Tried to stipulate script with '$arg', but a script file has already been stipulated via -f"=>
                              "'{$arg}'でスクリプトを指定しようとしているが、スクリプトは既に-f|--script-fileのスクリプト・ファイルで指定している。");
              }

              doctorScript( $val, $arg,
                            _ll("option '$arg' expects a script as its argument, "=>
                                "オプション'${arg}'の引数はスクリプトのはずであるが、"))
          }

          _dieUsage(  "'$val' looks like an option, but is positioned like a value for option '$arg'.  If '$val' really is the value for option $arg, use '$arg=$val'."=>
                      "'${val}'は自分自身がオプションに見えるが、コマンド行での位置を考慮すると'${arg}'の値にも見える。もし'${val}'は本当に'${arg}'の値なら、'$arg=${val}'で指定できる。")
              if  $val=~ /^-/  and  !exists $seenValS{$val};

          _dieUsage(  "Value for option ${arg} defined more than once."=>
                      "オプション${arg}を二度以上指定しようとしている。"  )    if  defined $opt_{$optName};
          $opt_{$optName}=  $val;
          $ARGVforPrintScript.=  " $arg$OPT_VALseparator$val"   if !exists $scriptOptS{$optName};
      }elsif(   exists $default{$optName}   ){
          my $val= 
              (   @ARGV   and   $ARGV[0]!~ /^-/  ||  exists $seenValS{$ARGV[0]}   )
              ?   shift @ARGV  :  $default{$optName};
          $opt_{$optName}=  $val;
          $ARGVforPrintScript.= " $arg$OPT_VALseparator$val";
      }else{#   Flag option.
          $opt_{$optName}=  't';
          $ARGVforPrintScript.= " $arg"   if $arg ne '-n';
      }
      $OPT_VALseparator=  ' ';
  }#WHILE  @ARGV

  $opt_{inputFieldSep}  //=  "\t";  #Defaults to tab even when -F not given as a flag.
  $opt_{outputFieldSep}  //=  $opt_{inputFieldSep};
  lock_hash %opt_;


  if(  $opt_{printVersionP}  ){
      say  "perltab $::VERSION   Perl Tab   Copyright (C) 2016,2017 Paul Horton\n";
      exit 0}                                                                       #EXIT PROGRAM

  if(  $opt_{printScriptP}  ){
      say '#perltab' . $ARGVforPrintScript . "\n";
      for  my $type  (  qw(BEGIN BEGIN2 HEAD HEAD2 DATA ROWS DATA2 ROWS2 END END2 SORT_KEY SORT_KEY2)  ){
          $opt_{$type}   or   next;
          say  "${type}{";
          say  $opt_{$type}=~ s/^(.+)/  $1/gr;
          say  '}';
      }
      exit 0;
  }

  $numRE_= $numRE_{$opt_{skipInfP}?  ($opt_{skipNanP}? 'plain' : 'NanOK'   )
                       :             ($opt_{skipNanP}? 'InfOK' : 'InfNanOK')  };


  pop @ARGV   if  @ARGV  and  $ARGV[0] eq '--';
  
  @ARGV   or   !-t *STDIN   or   $opt_{printEachRowP}   or   $opt_{BEGIN}
  or   _dieUsage( 'expected more command line arguments, perhaps you forgot the input file.'=>
                  'コマンド行の引数が足りない。入力ファイルを書き忘れた？');

  @ARGV   or   $opt_{BEGIN} && !$opt_{ROWS}   or   @ARGV=  ('-');

  dlock \@ARGV;


  #──────────  Check for contradictory options  ──────────
  _dieUsage('-p option cannot be combined with grepish options {-c,-g,-q}.',
            '-pオプションはgrep様のオプション{-c,-g,-q}との重複指定はできない。')
      if   $opt_{printEachRowP}  and  $opt_{countP} || $opt_{grepP} || $opt_{grepQuietP};

  if(  $opt_{SORT_KEY}   ){
      _dieUsage("$_ script not allowed with SORT_KEY script","-s|--sortスクリプトを指定すると${_}スクリプトは使用できない")
          for  grep {$opt_{$_}}  qw(HEAD HEAD2 DATA DATA2 SORT_KEY2);
  }
  if(  $opt_{SORT_KEY2}  ){
      _dieUsage("$_ script not allowed with SORT_KEY2 script","-s|--sortスクリプトを指定すると${_}スクリプトは使用できない")
          for  grep {$opt_{$_}}  qw(HEAD2 DATA2);
  }


}#END ━━━━━  Assign and check command line options  ━━━━━


# ──────────  Preload modules  ──────────
if(  $opt_{modules}  ){
    for my $module(  split ','=>$opt_{modules}  ){
        eval  "use $module";
        $@   and   _diell(  "Error when loading module: $module; $@\n"=>
                            "モジュール'{$module}'読み込み時にエラー${@}が発生した。");
    }
}


#──────────  Set up HEAD and DATA scripts  ──────────
%script_=  map {$_ => $opt_{$_} || undef}  qw(BEGIN BEGIN2 HEAD HEAD2 DATA ROWS DATA2 ROWS2 END END2 SORT_KEY SORT_KEY2);
lock_keys %script_;

if(  !$opt_{noHeadP}  ){
    $script_{HEAD} .=  ";\n$script_{ROWS}"    if  $script_{ROWS}  &&  $script_{HEAD};
    $script_{HEAD2} =      $script_{ROWS2}    if  $script_{ROWS2};                   
    $script_{HEAD}  //= $script_{ROWS};
    $script_{HEAD2} //= $script_{ROWS2};

}
$script_{DATA}  //= $script_{ROWS};
$script_{DATA2} //= $script_{ROWS2};


doctorScript(  $script_{$_}, _ll("$_ script constructed by combination"=>"組み合わせた${_}スクリプト"),
               _ll("When attempting to construct $_ script by combining two scripts, "=>"${_}スクリプトとしてふたつのスクリプトの組み合わせたところ、"))
    for  grep {$script_{$_}}  qw(HEAD DATA HEAD2 DATA2);
undef &doctorScript;

@script_{'SORT_KEY','SORT_KEY2'}=  @opt_{'SORT_KEY','SORT_KEY2'};
lock_hash %script_;


#━━━━━━━━━━   Set up globals dependent on %opt_  ━━━━━━━━━━
sub dittoMark(){#String used to mark ditto'ed values.
    state $dittoMark=   $opt_{dittoMark}  //(  $opt_{defaultDittoMarkP}? '' : undef  );
}

sub missingMark(){
    state $missingMark=   $opt_{missingMark}  //  '';
}

looks_like_number missingMark    and
    _dieUsage(  "missingMark cannot be set to '${\missingMark}' because it looks like a number."=>
                "missingMarkの値に数字'${\missingMark}'は指定できない。"  );


#━━━━━━━━━━  Check Input Filenames  ━━━━━━━━━━━━━━━━━━━━
{ my @pathname=   grep !/^-in2$/,  @ARGV;
  dlock \@pathname;

  2 > grep /^-$/, @pathname   or   _dieUsage( 'STDIN listed as input file more than once'=>
                                              '複数箇所に標準入力を入力と指定した。' );
  
  require Digest::MD5  if $opt_{testP};
  for  my $pathname  (@pathname){
      my $numDashes= 0;
      if(  $pathname eq '-' ){
          _dieUsage(  'STDIN stipulated for input, but is attached to a terminal.'=>
                      '標準入力を入力と指定したが、その入力先は端末になっている。'  )
              if -t *STDIN;
      }
      else{
          unless( -e $pathname ){
              my @msg= ("Data input file '${pathname}' does not exist."=>
                        "データ入力ファイル'${pathname}'は存在しない。");
              if(  _looksLikeScriptP( ${pathname} ) ){
                  _lln_concat( @msg,
                               'Perhaps that argument was intended to be a script, but you forgot to place -e (or -d etc.) before it?'=>
                               'もしかしたらスクリプトのつもりでしたがその前に-e (や-d等)を入れるの忘れたのでしょうか？');
              }
              _dieUsage(  @msg  );
          }

          -f $pathname   or   _dieUsage(  "Data input file '${pathname}' is not a plain file."=>
                                          "データ入力ファイル'${pathname}'は普通のファイルではない。"  );
          -r $pathname   or   _dieUsage(  "Data input file '${pathname}' is not readable."=>
                                          "データ入力ファイル'${pathname}'の読込み権限がない。"  );
          if(  $opt_{testP}  ){
              open my $inputFile, '<', $pathname   or   _diell("could not open input data file '$pathname', $!"=>
                                                               "入力データファイル'${pathname}が開けなかった。" . decode_utf8 $!);
              binmode $inputFile;
              say "MD5:$pathname\t", Digest::MD5->new->addfile($inputFile)->b64digest;
              close $inputFile;
            }
        }
  }#foreach $pathname

}#checkInputFilenames


if(  $opt_{outputFile}  ){
    -t STDOUT   or
        _dieUsage("'$opt_{outputFile}' parsed as an output filename, STDOUT has also been redirected."=>
                  "'$opt_{outputFile}'は出力ファイルのファイル名として指定されているが、標準出力も端末以外の出力先となっている。");
    open  my $outStream, '>', $opt_{outputFile}   or
        _dieUsage("could not open output file '$opt_{outputFile}', $!"=>
                  "出力ファイル'$opt_{outFile}'が開けなかった。" . decode_utf8 $!);
    select $outStream;   #To make user script print and say statements default to $outStream.
}


#String looks more like a script than a pathname?
sub _looksLikeScriptP{  $_[0] =~ /\s[@\$]\S/   }

sub _warn{
    select(   (select(STDERR), $|=1)[0]    );
    $|= 1;  print'';
    say STDERR  'perltab' . _ll(' warning: '=>'警告：') . _ll(@_);
    $|= 0;
}


# ────────────────────  Numish Related Utilities  ────────────────────
#Return true iff $_[0] is numish.  $_ used if argument omitted.
sub numP(_){    return 't'   if  defined $_[0] && $_[0]=~ /$numRE_/;
                return;
}


#Return $_[0] if it looks like a number, otherwise return 0.  $_ used if argument omitted.
sub num0(_){  numP $_[0] ? $_[0] : 0  }

#Return $_[0] if it looks like a number, otherwise return 1.  $_ used if argument omitted.
sub num1(_){  numP $_[0] ? $_[0] : 1  }

#Return true if all elements of @_ look like numbers.
sub allNum{   all  {numP $_}  @_   }
sub allnum{   all  {numP $_}  @_   }

#Filter non-number elements out of list
sub nums{   grep  {numP $_}  @_   }


#━━━━━━━━━━━━━━━━━━━━  Main loop  ━━━━━━━━━━━━━━━━━━━━
state sub openAndProcessInputFiles;
state sub openAndProcessInputFiles_forSorting;

dlock state $noncommentLineRegex=  qr/^[^#\n]/;   #Caveat: not sure if \n in character class will work for dos newlines.  PH20180823

dlock state $curPass= '';
state sub curPass(){  $curPass  }

{ 
  $, =  $opt_{outputFieldSep};
  _runScript 'BEGIN';
    
  my @inFilename1=  my @inFilename2=  @ARGV;
  dlock my $make2passesP=   $script_{DATA2}  ||  $script_{ROWS2}  ||  $script_{SORT_KEY2};
  dlock my $in2idx=  firstidx  {$_ eq '-in2'}  @ARGV;
  if(   $make2passesP   ){
      if(  $in2idx != -1  ){
          @inFilename1=  @ARGV[0   ..  $in2idx-1];
          @inFilename2=  @ARGV[$in2idx+1..$#ARGV];
      }

      any {$_ eq '-'} @inFilename1   and   any {$_ eq '-'} @inFilename2
          and   _dieUsage(  'standard out cannot be read in both input passes',
                            '標準入力が両方の入力読み込みpassに指定されている。');
  }else{
      $in2idx == -1   or   _dieUsage(  '-in2 flag given without any second pass script (e.g. the -e2 or -d2 options)'=>
                                       '-in2オプションを指定したが、２回目読み込み時のスクリプト(例えば、-e2又は-d2のオプション)は指定していない');
  }

  if(  $script_{SORT_KEY}   ){   openAndProcessInputFiles_forSorting  @inFilename1   }
  else                       {   openAndProcessInputFiles             @inFilename1   }
  _runScript 'END';
  
  $make2passesP   or   last;

  dunlock $curPass;  $curPass= '2';  dlock $curPass;
  _rezeroNRs;
  _runScript 'BEGIN2';

  if(  $script_{SORT_KEY2}  ){   openAndProcessInputFiles_forSorting  @inFilename2   }
  else                       {   openAndProcessInputFiles             @inFilename2   }

  _runScript 'END2';
}


say   NRT         if $opt_{countP};
exit (NRT? 0:1)   if $opt_{countP} || $opt_{grepP} || $opt_{grepQuietP};
exit 0;  #Return success to shell.
#━━━━━━━━━━━━━━━━━━━━━━━━━  END MAIN PROGRAM  ━━━━━━━━━━━━━━━━━━━━━━━━━


#  Return list of labels in @H similar to $query, or empty list if none found.
#  For convenient of printing, quotes are added.  e.g.  ('weight', 'height')
state sub candMistypedLabel{

    #Return edit distance of $s1, $s2
    #Character pairs, such as 'a' to 'A', that match except for case are given distance 0.1
    #Uses memory proportional to the length of the shorter string.
    my sub editDistance{
        @_ == 2   or   _dieProg;
        my( $s1, $s2 )=  sort {length $b <=> length $a}  @_;
        my( $l1, $l2 )=  map {length} ($s1, $s2);
        
        my @T=(   [0..$l2],
                  []
            );    #Dynamic programming table;

        my( $cur, $prv )  =  (0, 1);
        for  my $i  (1..$l1){
            $prv= $cur;  $cur= 1-$cur;#  $i % 2
            my $c1=  substr $s1, $i-1, 1;
            $T [$cur] [0]=  $i;   #Cost of skipping length $i prefix of string $s1.
            for  my $j  (1..$l2){
                my $c2=  substr $s2, $j-1, 1;
                my $d=    $c1  eq     $c2?   0
                    :  fc $c1  eq  fc $c2?   0.1
                    :                        1;
                $T[$cur][$j]  =  min(  $d +  $T [$prv] [$j-1],
                                       1  +  $T [$prv] [$j  ],
                                       1  +  $T [$cur] [$j-1]);
            }
        }
        return  $T [$cur] [$l2];
    }#sub editDistance


    #Main part of candMistypedLabel 
    my $query=  shift;
    my %dist =  map  {$_ => editDistance $query, $_ }  @H;
    my $minDist=  min values %dist;

    my @cand=  grep  {$dist{$_} == $minDist}  @H;

    grep  {$minDist < 0.5* min  map {length}  $query, $_}   @cand;


}#candMistypedLabel


#━━━━━━━━━━━━━━━━━━━━  User Functions  ━━━━━━━━━━━━━━━━━━━━
my $callerNF_;   #Temporary store for N() or F() caller name for error messages.

#────────────────────  Accessors  ────────────────────
my %N;

sub N{
    $callerNF_=  'N';
    goto &_N;
}

sub _N{
    my @retVal;
  ARG:
    for  my $query  (@_){
        if(  length ref($query)  ){
            _diell(  "Function ${callerNF_}() expected scalar column name argument but got reference to ".ref($query).'.'=>
                     "関数${callerNF_}()はスカラの列名を期待していたが、".ref($query).'への参照子を受け取った。'  );
        }
        length $query   or   _diell('Empty string given as column label.'=>
                                    '列ラベルとして空文字列を受け渡された。');

        exists $N{$query}   and       push( @retVal,  $N{$query} ),  next ARG;

        #  Code copy from _columnLabelPrefixMatches to avoid spending the time needed for a function call.
        for  my $prefRx  (qr/^$query$/, qr/(?i:^$query$)/, qr/^$query/, qr/(?i:^$query)/){
            my @match=  grep /$prefRx/, @H   or   next;
            if(  @match == 1  ){
                push  @retVal,  $N{$query}= $N{ $match[0]};         next ARG;
            }
        }

        _dieAllegedColumnLabelNotFound( $query, '' );
    }

    @_   or   _diell('${callerNF_}() called with no arguments.'=>'${callerNF_}()が引数無しで呼ばれた。');
    return  wantarray?  @retVal  :  $retVal[0];

}#sub _N


sub  F : lvalue{  $callerNF_=  'F';  return  @F[_N(@_)]   }


#  Return labels which prefix match $query.
#  Side-effect:  found labels are cached in %N to speed up future lookups.
sub _columnLabelPrefixMatches{
    my( $query )=  @_;

    my @match= ();
    for  my $prefRx  (qr/^$query$/, qr/(?i:^$query$)/, qr/^$query/, qr/(?i:^$query)/){
        if(   @match=  grep /$prefRx/, @H   ){
            $N{$query}=  $N{ $match[0]};    last;
        }
    }
    return @match;
}


#  $msgPrefix should be a phrase which can stand by itself, such as 'Error when calling insAft.'
#  It will be output as is (i.e. in whatever language it happens to be in).
sub _dieAllegedColumnLabelNotFound{
    my( $query, $msgPrefix )=  @_;

    my @match=  _columnLabelPrefixMatches( $query );

    @match == 1   and   _dieProg  "Nothing wrong with allegedly mistaken column label '$query'";

    if(  @match > 1  ){
        _diell("${msgPrefix}String '$query' matches prefices of column labels: '$match[0]', '$match[1]'..."=>
               "${msgPrefix}文字列'${query}'は複数の列ラベル'$match[0]', '$match[1]'...の前置詞である。");
    }
    
    # $query does not prefix match any column labels.
    my @cand=  candMistypedLabel $query;
    if   (  @cand==1  ){
        _diell("${msgPrefix}No column matches string '$query', perhaps you intended '$cand[0]?'"=>
               "文字列'${query}'に一致する列ラベルが見つからない。ラベル$cand[0]のつもりだったのでしょうか？");}
    elsif(  @cand> 1  ){
        _diell("${msgPrefix}No column matches string '$query', perhaps you meant one of {@cand}?"=>
               "文字列'${query}'に一致する列ラベルが見つからない。ラベル{@cand}のどれのつもりだったのでしょうか？");}
    else{
        _diell("${msgPrefix}No column matches string '$query'."=>
               "文字列'${query}'に一致する列ラベルが見つからない。");
    }
}#_dieAllegedColumnLabelNotFound


#stanza() returns dualvar yielding the stanza number in a numerical context.
#In a string context it includes trailing newlines so an otherwise empty string comment is "\n", while no comment at all is ''.
my $stanza;
my $stanzaPrintedP;

sub stanza(){   $stanza    }
state sub clearStanza(){   $stanza=  dualvar( $stanza, '' );
                            $stanzaPrintedP=  undef                      }

state sub zeroStanza() {   $stanza=  dualvar( 0, '');
                            $stanzaPrintedP=  undef                      }

state sub setStanza    {   $stanza=  dualvar ++$stanza,  join '', @_;
                            $stanzaPrintedP=  undef                      }

state sub printStanza_ifNotYetPrinted(){
    return    if  $stanzaPrintedP;
    print $stanza;
    $stanzaPrintedP=  't';
}


#━━━━━━━━━━  Core perltab subs  ━━━━━━━━━━
{ state $ih; 
  sub ih (){  $ih  }   #True iff current line is the head line.

  #  Read rows and run row script on contents of $inputFile
  #  $inputFile may be a reference to STDIN.
  #
  #  Named with 'file' instead of 'stream' to remind us that _processInputFile is called separately for each input file.
  #  e.g.
  #    % perltab -e SCRIPT in1.tsv in2.tsv in3.tsv   -->  Calls _processInputFile 3 times.
  state sub processInputFile{
      my(  $inputFile, $inputName   )  =  (@_);

      state $maxNumColsToExpect;
      state $prevPass= -1;
      state @Hin;
      state %Nin;
      state $head= undef;

      my $curPassFinalP=   curPass eq '2'  ||  !$script_{DATA2} && !$script_{HEAD2};
      my $printEachRowP=  $opt_{printEachRowP} && $curPassFinalP;


      goto START_PROCESSING_DATA_LINES  if $opt_{noHeadP};

      
      # ──────────────────────────────  Read and Process Head Line  ──────────────────────────────
      my $curRow;
      my @commentAtTop;
      while(   $curRow= <$inputFile>   ){
          chomp $curRow;   last if $curRow=~ /$noncommentLineRegex/;
          push @commentAtTop, $curRow;
      }
      defined $curRow   or   return;                                          #EMPTY INPUT SUB EXIT

      if(  $prevPass eq curPass  ){
          $head eq $curRow   or
              _diell("Input error; line at top of input file $inputName: '$curRow'\nnot equal to previously seen head line: '$head'", 
                     "入力エラー、ファイル${inputName}の先頭行: '${curRow}'\nは前に見たラベル行: '${head}'と一致しない。");
      }
      else{
          $prevPass= curPass;
          zeroStanza;
          $head= $curRow;  $ih= 1;

          dunlock \@Hin;
          @H= @Hin= @F= @Fin=  split $opt_{inputFieldSep}, $curRow, -1;
          dlock \@Hin;

          $maxNumColsToExpect=  @Hin;
          unlock_keys %N;  undef %N;
          for  my $i  (0..$#H){
              if(  exists $N{ $H[$i]}  ){
                  _diell(  "Input error: on first input line; label '$H[$i]' in column $i duplicates previous column label.
If the input has no column label head line you may use the '-N|--no-head' option."=>
                           "入力エラー、入力の１行目${i}番目の列ラベル'$H[$i]'はその前にある列ラベルと重複している。
もし入力に列ラベルがなければオプション'-N|--nohead'を使ってください。"  );
              }
              $N{ $H[$i]}=  $i;
          }
          %Nin=  %N;

          if(  $script_{'HEAD'.curPass}  ){
              my $scriptRetVal=  _runScript 'HEAD'.curPass;
              _incrNRs $scriptRetVal;
              if(   $scriptRetVal  &&  $curPassFinalP   ){
                  exit 0    if $opt_{grepQuietP};                               #PROGRAM EXIT POINT
                  if(  $opt_{grepP}  ){
                      say  for @commentAtTop;   say  $head;
                  }
              }
          }                  

          ($_=  firstidx  {!defined}  @F)  == -1   or   _diell
              "When attempting to output head line, \$F[$_] was undefined."=>
              "先頭行を出力しようとした時に、\$F[$_]はundef値だった。";
          
          do{   say for @commentAtTop;  say @F   }if  $printEachRowP;
          @FinPrev= @Fin;
      }# if/else $head already defined (from a preceding input file)


    START_PROCESSING_DATA_LINES:
      if(   $prevPass ne curPass   ){### assert:  $opt_{noHeadP}
          $prevPass=  curPass;
          zeroStanza;
      }

      $ih= 0;   my @comment;
      my $atTopOfStanzaP=  't';
      while(  <$inputFile>   ){
          #Handle comments.
          if(  !/$noncommentLineRegex/   ){
              push(  @comment,  $_);
              $atTopOfStanzaP=  't'    if  /^$/;
              next;
          }

          #Input is a non-comment line...
          chomp(  $curRow= $_);

          if(  @comment  &&  $atTopOfStanzaP){
              setStanza(  @comment);
              @comment=  ();
          }
          $atTopOfStanzaP=  undef;

          @Fin=   split  $opt_{inputFieldSep},  $curRow,  -1;

          $maxNumColsToExpect //=  @Fin;
          if(  @Fin > $maxNumColsToExpect   ){
              my $p= $maxNumColsToExpect;  my $n= @Fin;
              $opt_{extraFieldsOK}   or
                  _diell("Error unequal number of columns.  A previous row had $p columns, but on line $.: '$curRow' has $n columns.",
                         "列の本数が合わない。前の行では${p}本の列があったが、${.}行目の現在行には${n}列しかない。");
          }

          #Pad trailing columns with the missingMark if necessary.
          @Fin >= $maxNumColsToExpect   or   push  @Fin,  (missingMark) x  ($maxNumColsToExpect-@Fin);
              
          @F= @Fin;

          if(  defined dittoMark  ){
              for( 0..$#F ){
                  $F[$_]= $Fin[$_]=  $FinPrev[$_]     if  $F[$_] eq dittoMark;
              }
          }

          
          my $printCurRowP=  $printEachRowP;

          #Reinstate @H if necessary (perhaps should add  if $HmodifiedP  condition).
          @H= @Hin,  %N= %Nin    if  !$opt_{noHeadP};
          my $scriptRetVal=  _runScript 'DATA'.curPass;
          _incrNRs $scriptRetVal;
          if(   $scriptRetVal  &&  $curPassFinalP   ){
              exit 0    if $opt_{grepQuietP};                                   #PROGRAM EXIT POINT
              if(  $opt_{grepP}  ){
                  #Print head if it has not been printed yet  (NRT == 1)
                  #And will not be via the say @F below     (not in head)
                  if(   NRT == 1   and   !$opt_{noHeadP}   ){
                      say  for @commentAtTop;   say  $head;
                  }
                  $printCurRowP= 't';
              }
          }

          if(   $printCurRowP  ){
              printStanza_ifNotYetPrinted;
              if(  defined dittoMark  ){
                  state @FprevOut;   ### assert: $finalPassP_
                  if(  @FprevOut  ){
                      @F == @FprevOut   or  _diell("When printing, number of columns changed between formerly printed row '@FprevOut' and current row '@F'",
                                                   "行を出力しようとしたら、前に出力した行'@{FprevOut}'の列数と現在行'@{F}'の列数が一致しない。");
                      for( 0..$#F ){
                          if(  $F[$_] eq $FprevOut[$_]  ){    $F[$_]  =  dittoMark   }
                          else{                           $FprevOut[$_]  =  $F[$_]   }
                      }
                  }
                  else{
                      @FprevOut= @F;
                  }
              }#defined dittoMark

              $_ //= missingMark()  for @F;
              say @F;
          }

          @FinPrev= @Fin;
      }#  while(  $curRow = <INPUT_STREAM>  )
      
  }#processInputFile


  #Make one pass through input files.
  sub openAndProcessInputFiles{
      for my $filename (@_){
          if(  $filename eq '-'  ){
              processInputFile  *STDIN, '*STDIN'}
          else{
              open my $inputFile, '<', $filename   or   _diell("could not open input data file '$filename', $!",
                                                               "入力データファイル'${filename}が開けなかった。" . decode_utf8 $!);
              processInputFile  $inputFile, $filename;
          }
      }
  }#openAndProcessInputFiles

}#state $ih.


#────────────────────  Column Delete/Move/Insert functions  ────────────────────
{ state $callerName;   #For error messages.  Name of (trouble causing) function called by the user.

  state sub setCallerName{
      return   if $opt_{noHeadP};
      $callerName=  shift;
      state %seenCallS;
      if(  ih  ){
          $seenCallS{$callerName}{curPass()}=  undef;}
      else{
          return   if exists $seenCallS{$callerName}{curPass()};
          die _lln("$callerName must be called on head row because it needs to modify it, perhaps you used -d${\curPass}, when you meant -e${\curPass}?",
                   "${callerName}は先頭行を変更する為、先頭行に実行する必要がある。-e${\curPass}のつもりで間違って-d${\curPass}を使った？" );
      }
  }          


  # idx_of(  arg  )
  #  Returns positive numerical index into @F corresponding to $arg.
  #  $arg may be a label or positive or negative numerical index.
  #  Dies with error message listing caller, if $arg is invalid.
  #  Assumes @N, @F are up-to-date.
  state sub idx_of{
      @_   or   _dieProg  'idx_of called without arguments';

      my $arg=  $_[0];
      defined $arg   or   _dieProg  'idx_of called with undefined argument';

      return  $N{$arg}   if exists $N{$arg};                                         #LOOP EXIT POINT

      if(   $arg=~  /^[+-]? [0-9]+ $/x   ){
          if(  $arg < 0  ){   return $arg+@F   if $arg > -@F   }                     #LOOP EXIT POINT
          else            {   return $arg      if $arg <  @F   }                     #LOOP EXIT POINT

          my $callerName=  (caller 1)[3];   $callerName=~ s/^main:://;
          _diell(  "In $callerName; \$F[$arg] out of range.\n",
                   "${callerName}にて範囲外の添字\$F[$arg]\n"  );
      }

      my $callerName=  (caller 1)[3];   $callerName=~ s/^main:://;
      length $arg   or   _diell(  "In $callerName; empty string given as a column label.\n",
                                  "${callerName}は列ラベル引数として空文字列を受け渡された。\n"  );

      my @prefixMatch=  _columnLabelPrefixMatches( $arg );
      if(  @prefixMatch== 1  ){
          if(  exists $N{$arg}  ){
              return  $N{$arg};
          }else{
              _dieProg  "_columnLabelPrefixMatches should have cached column of '$arg' in %N...";
          }
      }

      _dieAllegedColumnLabelNotFound(  $arg,
                                       _ll  "In $callerName; " => "${callerName}で、"
                                       );
  }#idx_of()


  #Delete fields @_ from @F.  Fields may be given by its label or an integer in [0,$#F].
  sub del{
      @_   or   return;  #Deletion of empty list is noop.                             #SUB EXIT POINT
      setCallerName 'del';

      my @idx=  reverse sort  map{idx_of $_}  @_;
      my @label=  @H[@idx];
      while(  @idx  ){
          my $idx=  shift @idx;  my $label=  shift @label;

          @F=  @F[0..$idx-1, $idx+1..$#F];
          @H=  @H[0..$idx-1, $idx+1..$#H];
      }

      undef %N;  @N{@H}= 0..$#H;
  }
  


  #Internal function doing the work of insAft and insBef.
  #Those functions ensure validity of the first argument POS only.
  #insBef must check validity of the rest of its arguments.
  state sub insBef_{
      my $pos=  shift;
      @_   or   return;    #insert empty list is noop.                                #SUB EXIT POINT

      my @data;
      if(  $opt_{noHeadP}  ){
          @data=   map  {ref $_ eq 'CODE'?  &$_ : $_}  @_;
          splice  @F,    $pos,  0,  @data;
          return;                                                                     #SUB EXIT POINT
      }

      # ──────────  Else input has head line  ──────────
      if(  @_ % 2  ){
          _stringifyUndef @_;
          die _ll("In $callerName; -N|--no-head option not given, so $callerName expected an even number of (<LABEL, DATA_GEN> pair forming) trailing args, but got (@_)",
                  "${callerName}で、-N|--no-headオプションをしていないので、${callerName}は(<LABEL, DATA_GEN>の２組リストとなる)偶数の後尾引数を期待していたが、(@_)を受け取った。");
      }
      
      my @label=   splice  @_,  0,  @_/2;

      if(  ih  ){
          for(  @label  ){
              exists $N{$_}  &&  die _lln("In $callerName; Tried to insert duplicate column label '$_'",
                                          "${callerName}で、重複するラベル'${_}'を挿入しようとした。")
          }
          @data=  @label;}
      else{
          @data=   map  {ref $_ eq 'CODE'?  &$_ : $_}  @_;
      }        

      splice  @H,  $pos, 0, @label;
      splice  @F,  $pos, 0, @data;

      undef %N;  @N{@H}= 0..$#H;
  }#insBef_(pos)


  #  insAft(  PRE,  LABEL1,...,LABELn,  DATA_GEN1,...DATA_GENn  )
  #  Insert new column(s) into table after current column PRE.
  #
  #  PRE may be given by label or as an integer in [0,$#F] or (counting from final column) [-1..-@F].
  #  DATA_GEN can be either data or a sub reference to generate data.  See help documentation for details.
  #
  #  Calls insBef_ to do its work.
  sub insAft{
      @_   or   die _ll("$callerName called with empty argument list.", "引数無しで${callerName}を呼び出した。");
      setCallerName 'insAft';

      my $pre =idx_of shift;   unshift @_, $pre+1;   goto  &insBef_;
  }


  #  insBef(  SUC,  LABEL1,...,LABELn,  DATA_GEN1,...DATA_GENn  )
  #  Insert new column(s) into table before current column SUC.
  #  See insAft for details.
  #
  #  Calls insBef_ to do its work.
  sub insBef{
      setCallerName 'insBef';
      @_   or   die _ll("$callerName called with empty argument list.", "引数無しで${callerName}を呼び出した。");

      my $suc= idx_of shift;   unshift @_, $suc;     goto  &insBef_;
  }


  #  Internal function doing the work of movAft and movBef.
  #  Those functions ensure validity of the first argument POS only.
  #  movTo must check validity of the rest of its arguments.
  #  
  #  Implementation attempts to optimize move of contiguous ascending blocks of columns.
  #    e.g. when executing;  movTo( 20, 0..9 )
  #    The first 10 columns are moved as a block to the left of the 20th column.
  state sub movTo{
      my $pos=  shift;
      @_   or   return;    #Moving an empty list of columns is a noop.                #SUB EXIT POINT

      my @colsToMov;
      my @FtoMov;   #To hold the contents of @F to be moved.

      my( $curCol, $prvCol, $runBegCol, $runLen );
      if(  $opt_{noHeadP}  ){
          @colsToMov=   @_;

          #Push cols to move onto @colsToMov, one run (e.g., (6,7,8)) at a time.
          #Code takes advantage of entry conditions: 1) @colsToMov is not empty,  2) @H,@F do not contain undef.
          $prvCol   =  shift @colsToMov;
          $runBegCol=  $prvCol;
          while(  defined ($curCol = shift @colsToMov)  ){
              if(   $curCol  !=  $prvCol+1   ){   #If at end of run:
                  #Splice current run of indices in @colsToMov out of @H,@F, using undef's as place holders to keep coords valid.
                  $runLen=  1 + $prvCol - $runBegCol;
                  push  @FtoMov,   splice @F, $runBegCol, $runLen, (undef) x $runLen;
                  $runBegCol= $curCol;
              }
              $prvCol= $curCol;

          }
          #Push final run of indices in @colsToMov into @FtoMov.
          $runLen=  1 + $prvCol - $runBegCol;
          push  @FtoMov,   splice  @F,  $runBegCol,  $runLen,  (undef) x $runLen;

          #Splice columns to move back into @F.
          splice  @F,  $pos,  0,  @FtoMov;

          #Cleanup placeholders.
          @F=  grep {defined} @F;

          return;                                                                     #SUB EXIT POINT
      }#if $opt_{noHeadP}

#Navigation-->                                                                           sub movTo
      if(  ih  ){
          @colsToMov=   map  {idx_of $_}  @_;
          my %seen;
          for(  @colsToMov  ){
              _diell("In $callerName; column to move '$_' given twice"=>
                     "${callerName}で、移動すべき列'${_}'は重複している。"  )
                  if $seen{$_}++;
          }
      }
      else{
          #Not in head, so args have already been checked for validity.
          @colsToMov=   map  {exists $N{$_}?  $N{$_}  : $_}   @_;
      }

      my @HtoMov;   #To hold the contents of @H to be moved.

      #Push cols to move onto @colsToMov, one run (e.g., (6,7,8)) at a time.
      #Code takes advantage of entry conditions: 1) @colsToMov is not empty,  2) @H,@F do not contain undef.
      $runBegCol=  $prvCol=  shift @colsToMov;
      while(  defined ($curCol= shift @colsToMov)  ){
          if(   $curCol  !=  $prvCol+1   ){   #If at end of run:
              #Splice current run of indices in @colsToMov out of @H,@F, using undef's as place holders to keep coords valid.
              $runLen=  1 + $prvCol - $runBegCol;
              push  @HtoMov,   splice @H, $runBegCol, $runLen, (undef) x $runLen;
              push  @FtoMov,   splice @F, $runBegCol, $runLen, (undef) x $runLen;
              $runBegCol = $curCol;
          }
          $prvCol = $curCol;

      }
      #Push final run of indices in @colsToMov into @HtoMov,@FtoMov.
      $runLen=  1 + $prvCol - $runBegCol;
      push  @HtoMov,   splice  @H,  $runBegCol,  $runLen,  (undef) x $runLen;
      push  @FtoMov,   splice  @F,  $runBegCol,  $runLen,  (undef) x $runLen;

      #Splice columns to move back into @H,@F.
      splice  @H,  $pos,  0,  @HtoMov;
      splice  @F,  $pos,  0,  @FtoMov;

      #Cleanup placeholders.
      @H=  grep {defined} @H;
      @F=  grep {defined} @F;

      undef %N;  @N{@H}= 0..$#H;
  }#movTo(pos)


  #  movAft(  PRE,  @colsToMov   )
  #  Move columns @colsToMov after current column PRE.
  #
  #  PRE may be given by label or as an integer in [0,$#F] or (counting from final column) [-1..-@F].
  #  See the help documentation for details.
  #
  #  Calls movTo to do its work.
  sub movAft{
      setCallerName 'movAft';
      @_   or   die _ll("$callerName called with empty argument list.", "引数無しで${callerName}を呼び出した。");

      my $pre= idx_of shift;   unshift @_, $pre+1;   goto  &movTo;
  }

  #  movAft(  SUC,  @colsToMov   )
  #  Move columns @colsToMov before current column SUC.
  #
  #  SUC may be given by label or as an integer in [0,$#F] or (counting from final column) [-1..-@F].
  #  See the help documentation for details.
  #
  #  Calls movTo to do its work.
  sub movBef{
      setCallerName 'movBef';
      @_   or   die _ll("$callerName called with empty argument list.", "引数無しで${callerName}を呼び出した。");

      my $suc= idx_of shift;   unshift @_, $suc;     goto  &movTo;
  }


  #  swap( F1,F2,...,Fn, G1,G2,...,Gn )
  #  Swaps fields F1 and G1, then swaps fields F2 and G2, ...
  sub swap{
      @_    or   return;   #Move of empty list is noop.
      @_ %2   and   _dieProg  "In swap; expected even number of arguments, but got ".@_.'.';
      setCallerName 'swap';

      my $mid=  @_ / 2;
      while(  @_  ){
          my $idx1=  idx_of  shift;
          my $idx2=  idx_of  splice @_, --$mid, 1;
          @H[ $idx1, $idx2 ]=  @H[ $idx2, $idx1 ];
          @F[ $idx1, $idx2 ]=  @F[ $idx2, $idx1 ];
      }

      undef %N;  @N{@H}= 0..$#H;
  }#swap

}#state $callerName, %calledFromHeadP; sub idx_of(arg)


#Forgive likely misappelations.
sub  insaft{  goto &insAft  }   sub  insbef{  goto &insBef  }
sub  movaft{  goto &movAft  }   sub  movbef{  goto &movBef  }
sub moveAft{  goto &movAft  }   sub moveBef{  goto &movBef  }


#If in xterm, return $_[0] converted to bold
#Otherwise    return $_[0] as is.
sub _toBold{
    state $xtermP=  $ENV{TERM} =~ /^xterm/;
    return  $xtermP? "\e[1m$_[0]\e[0m"  : $_[0];
}


#────────────────────  Statistics Routines  ────────────────────

my @stats_data;  #Not visible to user.  Considering making this a PDL instead of a Perl array PH:20171120.

my $stats_numSigDig;  #Max number of significant digits in @stats_data, for prettier output.

my(  $stats_seenInfPlus, $stats_seenInfMnus, $stats_seenNaN  );

my $specialNumRE=  qr/^  (?: inf|nan )  \z/i;


#Analogous to @stats_data but used by functions such as pairwise correlation functions expecting pairs of values.
#Considering making this a PDL instead of a Perl array PH:20171120.
my @stats_pairData;  
my $stats_pairData_size;   #Number of pairs.

my $stats2_numSigDig;  #Max number of significant digits in @stats_dataData, for prettier output.


#  Push stats data onto @stats_pairData.
#  In future may switch to have this construct a PDL.
sub pushStats{
    (@_ % 2)   and   _diell
        'In the current implementation, pushStats expects exactly two arguments, but got ' .@_  =>
        '現状の実装ではちょうどふたつの引数を期待するが、' .@_. '個を受け取った。';

    goto &_pushStats2;
}


sub _pushStats2{
    defined $_[0]   or   _diell
        'pushStats called with undef value for first argument.'=>
        'pushStatsの第一引数としてundefを受け取った。';

    defined $_[1]   or   _diell
        'pushStats called with undef value for second argument.'=>
        'pushStatsの第二引数としてundefを受け取った。';

    if(  numP $_[0]  ){
        numP $_[1]   or   _diell
            "pushStats called with list in which pair the" .@stats_pairData. "th pair: (@_) is mixed numeric and non-numeric"=>
            "pushStatsが呼び出された引数リストの" .@stats_pairData. "番目の２組(@_)に数値と数値でない値が混在している";

        bemax  $stats2_numSigDig,  _numSigDigits( $_[0] ), _numSigDigits( $_[1] );
        push @stats_pairData, @_;
        ++$stats_pairData_size;
    }
    else{
        numP $_[1]   and   _diell
            "pushStats called with list in which pair the" .@stats_pairData. "th pair: (@_) is mixed numeric and non-numeric"=>
            "pushStatsが呼び出された引数リストの" .@stats_pairData. "番目の２組(@_)に数値と数値でない値が混在している";
        return;   #Pairs for which both values are non-numerical are quietly skipped.
    }
}


#───────────  Related to number of digits to display  ──────────

#Return number of digits in $_[0] which are not simply place holders.
#Always returns at least one.
sub _numSigDigits(_){
    dlock my $val=  shift;

    my $intPart;  my $frcPart=  '';
    if(  $val=~ /[.]/  ){
        ($intPart)=  $val=~ /^[+-]?   ( [0-9]* ) [.]/x;
        ($frcPart)=  $val=~ /     [.] ( [0-9]* )    /x;
    }
    else{
        ($intPart)=  $val=~ /^[+-]?([0-9]*)/;
    }

    if(  $frcPart  ){
        if(  !$intPart  ){
            my $sigPart=  $frcPart=~/^0*([1-9][0-9]*)/?  $1  :  '';
            return   max  1,  length $sigPart;
        }
        return  length( $intPart )  +  length( $frcPart );
    }
    else{
        $intPart=~ s/[+-]?E[0-9]+\z//i;   #Strip off exponent if present.
        $intPart=~ s/^(0+)\d//;           #Strip off any lead padding zeros.
        my $intPartRevd=   reverse $intPart;
        ( my $sigPart )=(  $intPartRevd=~ /^0*(\d+)\z/  )
            or   _dieProg  "Regex failed to find significant digits part of value $val, with intPart '$intPart, intPartRevd= '$intPartRevd'";
        return  length $sigPart;
    }
}#_numSigDigits


#  (Re)set stats @stats_data and related variables to reflect content of @_.
sub _stats_set_data{
    my $caller=  shift;
    {   my $undefIdx=  firstidx {!defined} @_;
        $undefIdx == -1   or
            _diell("${caller} called with an undef in place of ".(1+$undefIdx).'th data value'=>
                   "${caller}に渡された".(1+$undefIdx).'番目のデータ値はundefである。')
    }

    @stats_data=  ();
    $stats_seenNaN= $stats_seenInfPlus= $stats_seenNaN=  '';
    $stats_numSigDig=  1;

    @_   or   return;   #Called with empty list.  (Should not happen?)

    my $dataRf=  \@_;
    if(  ref $_[0] eq "ARRAY"  ){
        @_ == 1   or   _diell
            "${caller} received arguments following the first array reference argument."=>
            "${caller}の引数に問題があった。最初のARRAY参照子引数の後に余分な引数を受け取った。";
        $dataRf= $_[0];
    }

    for(  @$dataRf  ){
        /$numRE_/   or  next;
        if   (     /^[+-]? nan$/ix  ){   $stats_seenNaN=     $_   }
        elsif(     /^[+] ? inf$/ix  ){   $stats_seenInfPlus= $_   }
        elsif(     /^  - ? inf$/ix  ){   $stats_seenInfMnus= $_   }
        else{
            my $numSigDig=  _numSigDigits;
            $stats_numSigDig=  $numSigDig    if $numSigDig > $stats_numSigDig;
        }
        push @stats_data, $_;
    }

}


#  (Re)set stats @stats_data and related variables to reflect content of @_.
sub _stats_set_pairData{
    my $caller=  shift;

    @_   or   _diell
        "${caller} passed empty list"=>
        "${caller}が引数無しで呼び出された。";

    (@_ % 2)   and   _diell
        "${caller} expects even sized list but got a list of size " .@_. '.'=>
        "${caller}の引数個数" .@_. 'は偶数個ではなかった。';

    {   my $undefIdx=  firstidx {!defined} @_;
        $undefIdx == -1   or   _diell
            "${caller} called with an undef in place of ".(1+$undefIdx).'th data value'=>
            "${caller}に渡された".(1+$undefIdx).'番目のデータ値はundefである。';
    }

    @stats_pairData=  ();

    @_   or   return;   #Called with empty list.  (Should not happen?)

    $stats2_numSigDig=  1;

    if(  ref $_[0] eq "ARRAY"  ){
        _diell
            "${caller} called with array reference as first argument but that case is not yet implemented"=>
            "${caller}の第一引数がarray参照子でしたが、その呼び方はまだ実装されていない。";
    }

    my $numSigDig;
    for  my $pair  (pairs @_){
        my( $v1, $v2 )=  @$pair;
        if(  numP $v1  ){
            numP $v2   or   _diell
                "${caller} called with list in which pair the" .@stats_pairData. "th pair: ($v1,$v2) is mixed numeric and non-numeric"=>
                "${caller}が呼び出された引数リストの" .@stats_pairData. "番目の２組($v1,$v2)に数値と数値でない値が混在している";

            $numSigDig=  _numSigDigits $v1;
            defined $numSigDig  or  die  "numSigDiv of $v1 is undef!!";
            $stats2_numSigDig= $numSigDig    if  $numSigDig > $stats2_numSigDig;
            $numSigDig=  _numSigDigits $v2;
            defined $numSigDig  or  die  "numSigDiv of $v2 is undef!!";
            $stats2_numSigDig= $numSigDig    if  $numSigDig > $stats2_numSigDig;

            push @stats_pairData, @$pair;
        }
        else{
            numP $v2   and   _diell
                "${caller} called with list in which pair the" .@stats_pairData. "th pair: ($v1,$v2) is mixed numeric and non-numeric"=>
                "${caller}が呼び出された引数リストの" .@stats_pairData. "番目の２組($v1,$v2)に数値と数値でない値が混在している";
            next;   #Pairs for which both values are non-numerical are quietly skipped.
        }
    }#  for (pairs @_)

    $stats_pairData_size=  @stats_pairData / 2;
}


# reformat string part of numerical value to have a similar number of non-placeholder digits as the input data.
sub _stats_reformatNumstring{
    dualvar(  $_[0],
              sprintf  '%.*g', max( 4, $stats_numSigDig ),  $_[0]  )
}

sub _stats_reformatNumstrings{   map {_stats_reformatNumstring $_}  @_   }


sub _stats2_reformatNumstring{
    dualvar(  $_[0],
              sprintf  '%.*g', max( 4, $stats2_numSigDig ),  $_[0]  )
}


    
#  Return sum of numbers in @stats_data
sub _sum(){
    return                      if  !@stats_data;
    return  $stats_seenNaN      if  $stats_seenNaN;
    return  'NaN'               if  $stats_seenInfPlus && $stats_seenInfMnus;
    return  $stats_seenInfPlus  if  $stats_seenInfPlus;
    return  $stats_seenInfMnus  if  $stats_seenInfMnus;

    List::Util::sum @stats_data;
}


#  Return sum of numbers in @_
sub sum{   
    _stats_set_data 'sum', @_   if @_;

    @stats_data   or   _diell
        'tried to compute sum of list with no numbers.'=>
        '数値を含んでないリストの和を求めようとした。';

    return _stats_reformatNumstring _sum;
}


sub _mean(){
    my $sum=  _sum;
    return  $sum / @stats_data    if  $sum=~ $numRE_{plain};
    return  $sum;    # NaN, -inf, etc.
}


#  Arithmetic mean of numerical values in @_
sub mean{
    _stats_set_data 'mean', @_   if @_;

    @stats_data   or   _diell
        'tried to compute mean of list with no numbers.'=>
        '数値を含んでいないリストの平均値を求めようとした。';

    return _stats_reformatNumstring _mean;
}


sub _geomean(){
    return                      if  !@stats_data;
    return  $stats_seenNaN      if  $stats_seenNaN;
    return  'NaN'               if  $stats_seenInfPlus && $stats_seenInfMnus;
    return  $stats_seenInfPlus  if  $stats_seenInfPlus;
    return  $stats_seenInfMnus  if  $stats_seenInfMnus;

    my $retVal=  1;
    my $rootExp=  1 / @stats_data;

    for  my $val  (@stats_data){
        return 'NaN'   if $val < 0;
        $retVal *=  $val**$rootExp;
    }

    return $retVal;
}


#  Geometric mean of numerical values in @_
sub geomean{
    _stats_set_data 'geomean', @_   if @_;

    @stats_data   or   _diell
        'tried to compute geometic mean of list with no numbers.'=>
        '数値を含んでいないリストの幾何平均値を求めようとした。';

    return _stats_reformatNumstring _geomean
}
    

sub _stdev(){
    return                      if !@stats_data;
    return  $stats_seenNaN      if $stats_seenNaN;
    return  'NaN'               if @stats_data < 2;
    return  $stats_seenInfPlus  if $stats_seenInfPlus;
    return  'Inf'               if $stats_seenInfMnus;

    my $mean=  _mean;

    my $ssDev= 0.0;
    for (@stats_data){
        my $dev=  $_ - $mean;
        $ssDev+=  $dev * $dev;
    }

    return   sqrt( $ssDev )  /  (@stats_data - 1);
}


#  Sample standard deviation of numerical values in @_
sub stdev{
    _stats_set_data 'stdev', @_   if @_;
        
    @stats_data   or   _diell
        'tried to compute stdev of list with no numbers.'=>
        '数値を含んでないリストの標準偏差を求めようとした。';

    return _stats_reformatNumstring _stdev;
}


#  Return array holding counts of $@_ in $q equal width intervals
#  Still experimenting with interface...
sub _histc{
    my $caller=  shift;
    @_    or   _diell
        "${caller} needs at least one argument"=>
        "${caller}は少なくともひとつの引数が必要";

    my $q=  shift;
    $q =~ /^[1-9]\d*$/   or   _diell
        "${caller} expected first argument to a positive integer but got '$q'."=>
        "${caller}の第一引数は正整数でなければならないが、'${q}'を受け取った。";

    return (1)   if $q==1;

    if(  @_  ){
        ref $_[0] eq 'ARRAY'   or   _diell
            "currently ${caller} requires second argument (if present) to be an array reference, but got '${_[0]}'"=>
            "現状の実装では${caller}に第二引数を与える場合、そのタイプはarray参照子でなければならないが、'${_[0]}'を受け取った。";
    }

    _stats_set_data( $caller, @_  )    if @_;

    @stats_data   or   _diell
        'tried to compute ${caller} of list with no numbers.'=>
        '数値を含んでないリストに${caller}を呼んだ。';

    my(  $min, $max  )=  minmax @stats_data;
    $min=~ /inf/ || $max=~ /inf/   and   _diell
        "${caller} encountered infinite value"=>
        "${caller}は-inf値に出くわした。";

    my $span=  $max-$min;

    my @count=  (0) x ($q+1);

    for  my $val  (@stats_data){
        next   if  $val=~  /nan/i;
        ++$count[$q*($val-$min)/$span];
    }

    # Now $count[-1] is special, holding only values exactly equal to $max, so merge it with $count[-2].
    $count[-2]+= pop @count;

    return @count;
}


#  Return array holding frequencies of @_ in $q equal width intervals
#  Still experimenting with interface...
sub histc{
    unshift  @_,  'histc';
    goto  &_histc;
}


#  Return array holding frequencies of @_ in $q equal width intervals
#  Still experimenting with interface...
sub histf{
    unshift  @_,  'histf';
    my @freq=  _histc @_;
    my $sum=  sum @freq;;
    $_ /= $sum  for @freq;
    return @freq;
}


#  Return mean absolute error of paired data in @_
sub MAE{
    _stats_set_pairData 'MAE', @_   if @_;

    @stats_pairData    or      _diell
        'MAE() called with no numerical arguments to work with.'=>
        'MAE()が呼び出されたが、扱うべき数値は与えられていない。';

    my $sum= 0.0;
    for  my $pair  (pairs @stats_pairData){
        my( $v1, $v2 )=  @$pair;
        $sum+=  abs( $v1 - $v2 );
    }
    
    return  _stats2_reformatNumstring(  $sum / $stats_pairData_size  );
}


#  Return Pearson's linear correlation of paired data in @_
sub corrP{
    _stats_set_pairData 'corrP', @_   if @_;

    @stats_pairData    or      _diell
        'corrP() called with no numerical arguments to work with.'=>
        'corrP()が呼び出されたが、扱うべき数値は与えられていない。';

    @stats_pairData > 3   or   _diell
        'corrP() requires at least two pairs of numbers, but got only one pair.'=>
        'corrP()はふたつ以上の数値２組を必要とするがひとつしか受け取っていない。';


    my( $mean0, $mean1 )=  (0, 0);
    for(  pairs @stats_pairData  ){
        $mean0 +=  $$_[0];
        $mean1 +=  $$_[1];
    }
    {  my $n=  @stats_pairData / 2;
       $mean0 /= $n;
       $mean1 /= $n;
    }


    my( $diffInnerProd, $diffSqrd0, $diffSqrd1 ) =  (0, 0, 0);
    for  (pairs @stats_pairData){
        my $d0=  ($$_[0]-$mean0);
        my $d1=  ($$_[1]-$mean1);
        $diffInnerProd+=  $d0 * $d1;
        $diffSqrd0+=      $d0 * $d0;
        $diffSqrd1+=      $d1 * $d1;
    }
    
    return  _stats2_reformatNumstring(  $diffInnerProd / sqrt( $diffSqrd0 ) / sqrt $diffSqrd1  );
}


#  Return string summarizing pairwise correlations of input data.
#  UNDER CONSTRUCTION
#  Still thinking about the best interface and implementation for this.
sub corrs{
    @_ > 1   or   _diell
        'corrs expected at least two arguments, but got '.(@_? 'only one':'none')  =>
        'corrsはふたつ以上の引数が必要であるが、１個'.(@_? 'しか':'も').'受け取らなかった';

    
    if(  any {ref ne 'ARRAY'} @_  ){
        my $badIdx=  firstidx {ref ne 'ARRAY'} @_;
        my $argType=  ref $_[$badIdx];
        _diell(
            "corrs expected each of its arguments ARG to be an array reference, but ARG[$badIdx] is ".(length $argType? "a reference to $argType":'not a reference.')  =>
            "corrsの各引数ARGはarray参照子でなければならないが、ARG[$badIdx]は".(length $argType? "${argType}参照子である。":'参照子ではない。')
            );
    }


    for (0..$#_-1){
        @{$_[$_]} == @{$_[1+$_]}    or
            _diell(  "corrs expected references to equal size arrays, but encountered unequal sizes: size ARG[$_]=" .@{$_[$_]}. ' ≠ size ARG[' .(1+$_). ']=' .@{$_[1+$_]}  =>
                     "corrsの引数はすべて同じ大きさのarrayへの参照子でなければならないが、size ARG[$_]=" .@{$_[$_]}. ' ≠ size ARG[' .(1+$_). ']=' .@{$_[1+$_]}  );
    }
            
    my $retVal;

    $retVal=  'dummy return value';

    # Statistics::Basic::correlation( $X, $Y );
    # my $r=  Statistics::Basic::correlation( $X, $Y );
    # my $n=  0+@$X;
    # my $t=  $r * sqrt(  ($n-2) / (1-$r*$r)  )  if $r*$r < 1.0;


    # my $pVal = 'NaN';
    # $pVal = defined $t? 2 * Statistics::Distributions::tprob( $n-1, abs $t ) : 'NaN';


    # my $rankCorr  =  Statistics::RankCorrelation->new( $X, $Y );
    # print    "n: $n";
    # printf    ' r:%5.3f',   $r;
    # printf ' pVal:%5.3f',   $pVal;
    # printf    ' τ:%5.3f',   $rankCorr->kendall();
    # printf    " ρ:%5.3f\n", $rankCorr->spearman();

    return $retVal;
}#sub corrs


#  Return indicies into @stats_data for boundaries of $q bins
#  Sorts @stats_data as side effect.
sub _binBoundaryIdxs{
    my $arg1=  shift;

    @stats_data=  sort {$a<=>$b} @stats_data;

    if   (  $arg1 eq '-w'  ){
        goto &_binBoundaryIdxs_uniformRange;
    }
    elsif(  $arg1 eq '-o' ){
        goto &_binBoundaryIdxs_uniformOccupancy;
    }
    else{
        _diell(  "unknown binning spec '$arg1', supported specs are {-r,-o}."=>"データを区間に分割する仕様には{-r,-o}がある。'${arg1}'は無効な仕様'"  );
    }
}
    


#  Return indicies into @stats_data for boundaries of $q uniform size ranges
sub _binBoundaryIdxs_uniformRange{
    my $q=  shift;

    my(  $min, $max  )=  minmax @stats_data;
    
    my $range=  $max - $min;
    my $binWidth= $range / $q-1;

    return    map  {$min + $binWidth*$_}   1 .. $q-1;
}


#  Return indicies into @stats_data for boundaries of $q uniformly occupied bins.
#  Side effect: numerically sorts @stats_data.
sub _binBoundaryIdxs_uniformOccupancy{
    my $q=  shift;

    @stats_data=  sort {$a<=>$b} @stats_data;


    my @retVal;


    #Assign indices with propotional intervals over range [0,$#stats_data], since @stats_data is sorted, this gives approximate uniform occupancy boundary indices.
    #The rest of the code below is to make the bins more uniformly occupied in the case of ties (runs of the same value in the sorted data).
    my $prevBound=  -1;  #Uninitialized.
    for  my $iProp  (map  {$#stats_data*$_/$q}  1..$q-1){
        my $iLw=  my $iUp=  ceil $iProp;
        my $valProp=  $stats_data[$iProp];

        $iUp++   while  $iUp < $#stats_data   and   $stats_data[$iUp+1] == $stats_data[$iUp];
        $iUp++      if  $iUp < $#stats_data;

        $iLw--   while  $iLw > 0              and   $stats_data[$iLw-1] == $stats_data[$iLw];
        $iLw--      if  $iLw > 0;

        #At this point $iLw is either the minimum index possible or is the greatest index with a value smaller than $valProp.
        #Symmetrically $iUp is either the maximum index possible or is the smallest index with a value greater than $valProp.


        my $newBound=   ($iUp-$iProp < $iProp-$iLw)
            ?  ( $stats_data[$iUp] + $valProp ) / 2
            :  ( $stats_data[$iLw] + $valProp ) / 2;
        
        #Push new boundary value if unique.
        push  @retVal,  $prevBound= $newBound    if $newBound != $prevBound;
    }

    return @retVal;
}# _binBoundaryIdxs_uniformOccupancy


sub binner{
    unshift @_, 'binner';
    goto &_binner;
}

#  Return reference to subroutine to do uniform occupancy binning into $q bins based on the data in @_[1...] (defaulting to @stats_data).
#  In the future other types of binning might be considered, e.g.  binner( -r, 4, @DATA ) might stipulate uniform range binning
sub _binner{
    my $caller=  shift;

    wantarray   and   _diell
        "${caller}() called in list context."=>
        "${caller}()がリスト文脈で呼ばれた。";

    @_   or   _diell
        "${caller}() called with no arguments."=>
        "${caller}()が引数無しで呼び出された。";

    my $q=  shift;

    _stats_set_data $caller, @_   if @_;

    @stats_data   or  _diell(  "${caller}() called without numerical values to work with."=>
                               "${caller}()が呼び出されたが、扱うべき数値は与えられていない。"  );

    my @bound= _binBoundaryIdxs $q;

    return
        sub{
            my $part=  firstidx {$_[0] <= $_}  @bound;
            $part > -1   or   $part= @bound;
            return $part;
    }
}#_binner


#  Return bin numbers of data in @_[1...] (defaulting to @stats_data), binned into $q bins with approximately uniform occupancy.
#  In the future other types of binning might be considered, e.g.  binner( -r, 4, @DATA ) might stipulate uniform range binning.
sub binned{
    wantarray   or   _diell
        'binned() called in non-list context.'=>
        'binned()が非リスト文脈で呼ばれた。';

    @_   or   _diell
        'binned() called with no arguments.'=>
        'binned()が引数無しで呼び出された。';

    my @dataCopy=  @_[1..$#_];

    my $binner=  _binner 'binned', @_;

    numP   and   $_=  $binner->($_)    for @dataCopy;

    return @dataCopy;
}
    
    


#  xqtiles($q,@num) returns the $q extended quantiles of @num (@num defaults to @stats_data),
#  where extended means the minimum and maximum value are included as the 0th and qth quantiles respectively.
sub xqtiles{
    unshift @_, 'xqtiles';
    goto &_qtiles;
}

#  qtiles($q,@num) returns the $q quantile of @num (@num defaults to @stats_data).
#  For example, when called with q=4, it will return 3 values.
sub qtiles{
    unshift @_, 'qtiles';
    goto &_qtiles;
}


#  _qtiles($caller,$q,@num) returns the $q quantile of @num (@num defaults to @stats_data).
#  When $caller eq 'xqtiles', the min and max values of @stats_data are also included in the returned list.
#  Side effect: numerically sorts @stats_data.
sub _qtiles{
    my $caller=  shift;
    
    @_   or   _diell
        '${caller}() called with no arguments.'=>
        '${caller}()が引数無しで呼び出された。';

    my $q=  shift;
    $q=~ /^[1-9]+[0-9]*/   or  _diell
        "${caller}() expected positive integer as its first argument but got '$q'"=>
        "${caller}()は最初の引数として整数値を期待したが、'${q}'を受け取った。";

    $q > 1   or   _diell
        "${caller}() first argument should be at least 2 but got '$q'"=>
        "${caller}() 第一引数は２≧でないといけないが、'${q}'を受け取った。";

    _stats_set_data $caller, @_   if @_;
        
    @stats_data   or   _diell
        '${caller}() called with no numerical arguments to work with.'=>
        '${caller}()が呼び出されたが、扱うべき数値は与えられていない。';

    $q < @stats_data   or   _warn
        "{caller}(${q},...) called with only ".@stats_data.' numbers.'=>
        "{caller}(${q},...)は".@stats_data.'個の数値引数しか受け取れなかった。';

    @stats_data=  sort {$a<=>$b} @stats_data;

    my $fin=  $#stats_data;

    my @qtile;
    for  my $t  (1..$q-1){
        my $quotient=  $t*$fin / $q;
        my $dw=  floor $quotient;
        my $up=  ceil  $quotient;
        my $r=   $quotient - $dw;
        push @qtile,  (1-$r) * $stats_data[$dw]  +  $r * $stats_data[$up];
    }

    if(  $caller eq 'xqtiles'  ){
        unshift  @qtile, $stats_data[ 0];
        push     @qtile, $stats_data[-1];
    }

    return _stats_reformatNumstrings @qtile;
}# _qtiles.


sub stats{
    _stats_set_data 'stats', @_   if @_;

    @stats_data    or      _diell
        'stats() called with no numerical arguments to work with.'=>
        'stats()が呼び出されたが、扱うべき数値は与えられていない。';

    my $stats=  'n:'. @stats_data;
    @stats_data > 0   or   return  $stats;

    $stats.=  '  μ' . mean;
    @stats_data > 1   or   return  $stats;

    $stats.=    '±' . stdev;
    @stats_data > 2   or   return $stats;

    my @q4tile=  xqtiles 4;

    $stats.=  "  Q4s:$q4tile[0] $q4tile[1] " . _toBold( $q4tile[2] ) . " $q4tile[3] $q4tile[4]";
    return $stats;
}


#────────────────────  Convenience Routines  ────────────────────
#  Return log base 2 or 10 of arguments.  Let non-numeric arguments pass through unchanged.

{
    state sub _log2(_){
        return $_[0]    if  !numP $_[0];

        isneg $_[0]   and    _diell
            "log2 called with negative argument '$_[0]', but the logarithm of a negative number is mathematically undefined."=>
            "log2が負の引数'$_[0]'で呼び出されたが、負の数の対数は数学的に定義されていない。";

        $_[0] == 0   and   _diell
            "log2 called with '$_[0]', but the logarithm of zero is mathematically undefined."=>
            "log2の引数に'$_[0]'が含まれていたが、０の対数は数学的に定義されていない。";

        #If $_[0] is a floating point number or an integer which is not a power of 2, use the standard log function to compute log2.
        return  log( $_[0] ) / log(2)    if   isfloat $_[0]  or  $_[0] & ($_[0] -1);

        #$_[0] is a positive power of 2, in this case compute and return integer value for log2.
        my( $x, $intRetVal )=  ( $_[0], 0 );

        ++$intRetVal   while $x >>= 1;
        return $intRetVal;
    }


    state sub _log10(_){
        return $_[0]    if  !numP $_[0];

        isneg $_[0]   and   _diell
            "log10 called with negative argument '$_[0]', but the logarithm of a negative number is mathematically undefined."=>
            "log10が負の引数'$_[0]'で呼び出されたが、負の数の対数は定義されていない。";

        $_[0] > 0   or   _diell
            'log10 called with 0, but the logarithm of a zero is mathematically undefined.'=>
            'log10の引数に0が含まれているが、０の対数は定義されていない。';

        #If $_[0] is a floating point number use the standard log function to compute log10.
        return  log( $_[0] ) / log(10)    if   isfloat $_[0];

        my $valStr=  sprintf( '%d', $_[0]  );

        return length $1   if $valStr=~ /^1(0+)$/;

        return  log( $_[0] ) / log(10);
    }


    sub log2(_){
        defined wantarray   or   _diell
            'log2 called in void context.'=>
            'log2がvoid文脈に呼び出された。';

        @_   or   unshift @_, $_;

        defined $_[0]   or   _diell
            'log2 called with undefined value.'=>
            'log2の引数にundef値で含まれていた。';

        wantarray   or   @_==1   or   _diell
            'log2 called in scalar context, but with multiple arguments.'=>
            'log2はスカラ文脈で呼び出されたが、返すべきリストに'.@_.'個の引数が含まれている。';

        my @retVal=  map  {_log2}  @_;
        return(  wantarray? @retVal : $retVal[0]  );
        _diell
            'log2 called in void context.'=>
            'log2がvoid文脈に呼び出された。';
    }


    sub log10(_){
        defined wantarray   or   _diell
            'log10 called in void context.'=>
            'log10がvoid文脈に呼び出された。';

        @_   or   unshift @_, $_;

        defined $_[0]   or   _diell
            'log10 called with undefined value.'=>
            'log10の引数にundef値で含まれていた。';

        wantarray   or   @_==1   or   _diell
            'log10 called in scalar context, but with multiple arguments.'=>
            'log10はスカラ文脈で呼び出されたが、返すべきリストに'.@_.'個の引数が含まれている。';

        my @retVal=  map  {_log10}  @_;
        return(  wantarray? @retVal : $retVal[0]  );
        _diell
            'log10 called in void context.'=>
            'log10がvoid文脈に呼び出された。';
    }
}


#────────────────────  Sorting Routines  ────────────────────
state sub processInputFile_forSorting;
state sub sortData_andPrint;

#Make pass through input files.
sub openAndProcessInputFiles_forSorting{

    for my $filename (@_){
        if(  $filename eq '-'  ){
            processInputFile_forSorting  *STDIN, '*STDIN'}
        else{
            open my $inputFile, '<', $filename   or   _diell("could not open input data file '$filename', $!"=>
                                                             "入力データファイル'${filename}が開けなかった。" . decode_utf8 $!);
            processInputFile_forSorting  $inputFile, $filename;
        }
    }

    sortData_andPrint();
}#openAndProcessInputFiles_forSorting


my @sort_row;
my @sort_keyVec;
my @sortPerm;  #To hold permutation of rows, such that sort_row[ @sortPerm ] equals the rows in sorted order.


#  Before sorting, the stanza consists of row numbers [oldBeg, oldFin].
my %stanza__oldBeg_comment;
my %stanza__oldBeg_oldFin;
my %stanza__newBeg_comment;

my $prevStanzaRN;

state sub configure_sort;
state sub sortKeylev_recurseOnTies;

sub sortData_andPrint{
    configure_sort;

    @sortPerm=  0 .. $#sort_row;
    sortKeylev_recurseOnTies   0,  0, 0+@sortPerm;
    dlock \@sortPerm;

    if(  defined $prevStanzaRN  ){
        $stanza__oldBeg_oldFin{$prevStanzaRN}=  $#sortPerm;

        my %newBoundaryS=  (0=>undef, 0+@sortPerm=>undef);
        my %newBeg_newEnd;

        #Place intact stanza comments into %stanza__newBeg_comment.
        for  my $i  (0..$#sortPerm){
            my $oldBeg=  $sortPerm[$i];
            exists $stanza__oldBeg_oldFin{$oldBeg}   or   next;

            #See if stanza is intact
            my $oldFin=  $stanza__oldBeg_oldFin{$oldBeg};

            my( $newBeg, $newFin)=  ($i,$i);
            $newBeg--  while  $newBeg > 0           &&  $sortPerm[$newBeg-1] >= $oldBeg  &&  $sortPerm[$newBeg-1] <= $oldFin;
            $newFin++  while  $newFin < $#sortPerm  &&  $sortPerm[$newFin+1] >= $oldBeg  &&  $sortPerm[$newFin+1] <= $oldFin;
            if(  $newFin - $newBeg  ==  $oldFin - $oldBeg  ){
                $stanza__newBeg_comment{$newBeg}=
                    $stanza__oldBeg_comment{$oldBeg}   if exists $stanza__oldBeg_comment{$oldBeg};
                $newBoundaryS{$newBeg}=  undef;
                $newBeg_newEnd{$newBeg}=  $newFin+1;

            }
        }#for  $i (0..$#sortPerm)

        #Remove stanza comments which are not bracketed.
        while(   my( $newBeg, $newEnd )=  each %newBeg_newEnd   ){
            if(   !exists $newBoundaryS{$newBeg}  or
                  !exists $newBoundaryS{$newEnd}    ){
                delete $stanza__newBeg_comment{$newBeg};   #Because those rows are no longer bracketed.
            }
        }
    }


    say  @H    unless $opt_{noHeadP};

    #Print results of sort.
    if(  defined dittoMark  ){
        for  my $i  (0..$#sortPerm){
            my @F= my @Fverb=   split  $opt_{inputFieldSep},  $sort_row[ $sortPerm[$i]],  -1;
            $i  and  $F[$_] eq $FinPrev[$_]   and   $F[$_]= dittoMark    for 0..$#F;
            print  $stanza__newBeg_comment{$i}   if exists $stanza__newBeg_comment{$i};
            say @F;    @FinPrev= @Fverb;
        }}
    else{
        for  my $i  (0..$#sortPerm){
            print  $stanza__newBeg_comment{$i}   if  exists $stanza__newBeg_comment{$i};
            say  $sort_row[ $sortPerm[$i]];
        }
    }
}#sortData_andPrint


sub processInputFile_forSorting{
    my(  $inputFile, $inputName  )  =  (@_);

    state $head;
    state $prevPass= -1;

    goto START_PROCESSING_DATA_LINES_FOR_SORT  if $opt_{noHeadP};


    # ──────────────────────────────  Read and Process Head Line  ──────────────────────────────
    my $curRow;
    my @commentAtTop;
    while(   $curRow= <$inputFile>   ){
        chomp $curRow;   last   if $curRow =~ /$noncommentLineRegex/;
        push @commentAtTop, $curRow;
    }

    defined $curRow   or   return;                                              #EMPTY INPUT RETURN

    if(  $prevPass eq curPass  ){
        while(   $curRow= <$inputFile>   ){
            chomp $curRow;   last if $curRow=~ /$noncommentLineRegex/;   #Quietly skip comment lines.
        }
        $head eq $curRow   or
            _diell("Line at top of input file $inputName: '$curRow'\nnot equal to previously seen head line: '$head'", 
                   "入力ファイル${inputName}の先頭行: '${curRow}'\nは前に見たラベル行: '${head}'と一致しない。");
    }
    else{
        $prevPass= curPass;
        say  for @commentAtTop;
        zeroStanza;
        $prevStanzaRN= undef;
        undef %N;
        $head=  $curRow;
        @H =  split $opt_{inputFieldSep}, $curRow, -1;
        for  my $i  (0..$#H){
            my $label=  $H[$i];
            _diell(  "Input error; Duplicate label '$label'", "入力エラー、ラベル${label}が重複している。"  )    if exists $N{$label};
            $N{$label}=  $i;
        }
        _incrNR;
    }
    @FinPrev= (dittoMark x @H)   if defined dittoMark;


  START_PROCESSING_DATA_LINES_FOR_SORT:
    if(   $prevPass ne curPass   ){### assert:  $opt_{noHeadP}
        $prevStanzaRN=  undef;
        $prevPass=  curPass;
        zeroStanza;
    }

    my @comment;
    while(  <$inputFile>   ){
        push( @comment, $_ ), next   if  !/$noncommentLineRegex/;
        
        chomp(  $curRow= $_  );

        @F= @Fin=  split $opt_{inputFieldSep}, $curRow, -1;

        #Store row in sort_row
        if(  defined dittoMark  and  @FinPrev  ){
            $F[$_] eq dittoMark   and   $F[$_]= $FinPrev[$_]    for 0..$#F;
            push  @sort_row,  join $opt_{outputFieldSep}, @F;}
        else{
            push  @sort_row,  $curRow;
        }

        #Handle stanza comment.
        if(  @comment  ){
            setStanza  @comment;
            @comment= ();

            #Finish up saving info for previous stanza.
            $stanza__oldBeg_oldFin{$prevStanzaRN}=  $#sort_row-1
                if  defined $prevStanzaRN;

            $stanza__oldBeg_comment{$#sort_row}=  stanza;
            $prevStanzaRN=  $#sort_row;
        }


        #Compute and store sort keys.
        my @retVal=  _runScript 'SORT_KEY'.curPass;
        $sort_keyVec[$_][NR-1]=  $retVal[$_]    for 0..$#retVal;
        
        _incrNR;
        @FinPrev= @F;  #Might try (\@FinPrev, \@F)= (\@F, \@FinPrev)  If reference aliasing becomes standard.
    }#while reading data lines.

}#processInputFile_forSorting


my %sortCmpSpecs;


#Set $forPass entry of %sortCmpSpecs based on $sortSpecsStr.
sub _parseSortSpecsStr{
    dlock my $argBeingParsed= shift;
    dlock my $sortSpecsStr=   shift;
    dlock my $forPass=        shift;

    my sub dieSpec{
        _dieUsage( "\nWhen parsing command line arg '$argBeingParsed' as specification for sorting; " . $_[0],
                   "\nコマンド行の引数'${argBeingParsed}'をソートの指定として処理中に、"                . $_[1] );
    };

    $sortCmpSpecs{$forPass}= [];

    #Extract and upcase comma deliminated (possibly empty) sort specs.
    my @sortSpec=  "\U$sortSpecsStr," =~  /  (?: ([^,]*) ,)  /gx;

    for my $spec (@sortSpec){
        length $spec  or  $spec= 'AB01NM';  #Default spec.

        dlock my $atom=  qr/ 01 | 10 | AB | BA | M | N | U /x;
        $spec=~  /^ $atom{1,5} $/x   or
            dieSpec( "Invalid sort spec: '$spec', should look 1-5 slots each containing one of {01, AB, N, M, U}"=>
                     "並べ替え方の指定は１〜５個の{01, ab, N, M, U}の要素を連ねた文字列でなければならないが、受け取った文字列'${spec}'はこのパターンに合わない。");

        $spec =~ /01|10/   or   $spec !~ /N/   or
            dieSpec(  "Suspect sort spec: '$spec', 'N' stands for NaN (not a number) and should be accompanied with '01' or '10'"=>
                      "並べ替え方の指定'${spec}'を受け取ったが、'N'はNaN (not a number)を意味し、数値の並べ替えを指定する場合'01'か'10'も必要になる。"  );
            
        my( $badC )=  $spec=~ /([^01ABMNU])/;
        $badC   and   dieSpec( "encountered character '$badC', but only chars [01abMNU] are allowed in a sort comparison spec"=>
                               "文字'${badC}'に当ったが、並べ替え方の指定に[01abMN]以外の文字は許されない。");
        { my %freq;
          $freq{$_}++  for split //, $spec;
          $badC=   first{ 1 < $freq{$_} }  keys %freq;
        }
        $badC   and   dieSpec( "character '$badC' appears more than once, which is not allowed in a sort comparison spec"=>
                               "文字'${badC}'が２回以上に現れたが、並べ替え方の指定に文字の繰り返しは許されない");

        push  @{ $sortCmpSpecs{$forPass}},  $spec;
    }
}#_parseSortSpec


my @sortTyper;
my @sortPrecs;     #sortPrecs[i][j] to hold the precedence of type j when sorting on the ith key.

#  To understand why sortAlfT is ternary, but sortRevNumP is boolean,
#  note that when NUM is treated as a separate type by Typer, it is always supposed to be sorted (ascending or descending),
#  but since ALF is a fallback type sometimes ALF values should not be sorted,  for example a non-numerical string value under sort spec '01'.
my @sortAlfT;      #sortAlfT[i]=  {+1 for forward sorting, -1 for reverse sorting, 0 for no sorting.
my @sortRevNumP;   #True if numeric sorting for the ith key should be reversed.


my @sortPegUndefP; #$sortPegUndefP[KL] true iff undef should be pegged at key level KL.
my( $NUM, $NAN, $MIS, $UND, $ALF, $FIN );

sub configure_sort{
    dlock $UND= 0;  dlock $NUM= 1;   dlock $NAN= 2;   dlock $MIS= 3;   dlock $ALF= 4;   dlock $FIN= 4;

    my @cmpSpec=   @{ $sortCmpSpecs{curPass() }};   #Convenience copy.

    #Code below assumes the missingMark is not set to anything which looks_like_number.
    my %Typer=(
        UA   =>  sub{   #Only undef special.
            defined $_[0]            or  return $UND;
            ;                            return $ALF},
        A    =>  sub{   #undef pegged
            ;                            return $ALF},

        UMA  =>  sub{   #Undef and missing value mark special.
            defined $_[0]            or  return $UND;
            $_[0] eq missingMark    and  return $MIS;
            ;                            return $ALF},
        MA  =>   sub{   #undef pegged
            $_[0] eq missingMark    and  return $MIS;
            ;                            return $ALF},

        UA0  =>  sub{   #Numerical values special.
            defined $_[0]            or  return $UND;
            $_[0]=~ /^-?nan$/i      and  return $ALF;  #because looks_like_number accepts NaN.
            looks_like_number $_[0] and  return $NUM;
            ;                            return $ALF},
        A0  =>  sub{   #undef pegged
            $_[0]=~ /^-?nan$/i      and  return $ALF;  #because looks_like_number accepts NaN.
            looks_like_number $_[0] and  return $NUM;
            ;                            return $ALF},

        UNA0 =>  sub{   #Numerical values and NaN special.
            defined $_[0]            or  return $UND;
            $_[0]=~ /^-?nan$/i      and  return $NAN;
            looks_like_number $_[0] and  return $NUM;
            ;                            return $ALF},
        NA0  =>  sub{   #undef pegged
            defined $_[0]            or  return $UND;
            $_[0]=~ /^-?nan$/i      and  return $NAN;
            looks_like_number $_[0] and  return $NUM;
            ;                            return $ALF},

        UMA0 =>  sub{   #Missing value mark and numerical values (but not NaN) special.
            defined $_[0]            or  return $UND;
            $_[0] eq missingMark    and  return $MIS;
            $_[0]=~ /^-?nan$/i      and  return $ALF;  #because looks_like_number accepts NaN.
            looks_like_number $_[0] and  return $NUM;
            ;                            return $ALF},
        MA0 =>  sub{    #undef pegged
            $_[0] eq missingMark    and  return $MIS;
            $_[0]=~ /^-?nan$/i      and  return $ALF;  #because looks_like_number accepts NaN.
            looks_like_number $_[0] and  return $NUM;
            ;                            return $ALF},

        UNM0A=>  sub{   #Missing value mark, numerical values, and NaN are all special.
            defined $_[0]            or  return $UND;
            $_[0] eq missingMark    and  return $MIS;
            $_[0]=~ /^-?nan$/i      and  return $NAN;
            looks_like_number $_[0] and  return $NUM;
            ;                            return $ALF},
        NMA0=>   sub{   #undef pegged
            $_[0] eq missingMark    and  return $MIS;
            $_[0]=~ /^-?nan$/i      and  return $NAN;
            looks_like_number $_[0] and  return $NUM;
            ;                            return $ALF}
        );
    lock_hash %Typer;


    for  my $keyLev (0..$#sort_keyVec){
        my $spec=   $cmpSpec[  $keyLev > $#cmpSpec?  -1  :  $keyLev  ];

        $spec=~  /^[01ABMNU]+$/   or   _dieProg  "Only chars [01ABMNU] allowed in sort spec: '$spec'";

        $sortAlfT[$keyLev]=  0;
        $sortPrecs[$keyLev][$_]=  undef    for  0 .. $FIN;

        my $prec;  my $typerSpec;
        my $specRev=  reverse $spec;
        while(  length $specRev  ){
            my $c=  chop $specRev;
            $typerSpec.= $c;

            if   (  $c=~ /[01]/  ){   $sortPrecs[$keyLev][$NUM]=  $prec++;
                                      chop $specRev;
                                      $sortRevNumP[$keyLev]=  ($c eq '1');
            }
            elsif(  $c=~ /[AB]/  ){   $sortPrecs[$keyLev][$ALF]=  $prec++;
                                      chop $specRev;
                                      $sortAlfT[$keyLev]=  ($c eq 'A')?  +1  :  -1;
            }
            elsif(  $c eq 'N'  ){     $sortPrecs[$keyLev][$NAN]=  $prec++       }
            elsif(  $c eq 'M'  ){     $sortPrecs[$keyLev][$MIS]=  $prec++       }
            elsif(  $c eq 'U'  ){     $sortPrecs[$keyLev][$UND]=  $prec++       }
            else{   _dieProg  "bad char '$c'"   }
        }

        $sortPrecs[$keyLev][$ALF] //= $prec;

        $typerSpec=~ /[AB]/   or   $typerSpec.= 'A';   #non-numerical strings need to be their own type, even when not sorting them.
        $typerSpec=~ y/1B/0A/;  $typerSpec=  reverse  sort  split //, $typerSpec;   #Canonicalize typerSpec.
        $sortPegUndefP[$keyLev]=  $typerSpec!~ /U/;

        $sortTyper[$keyLev]=  $Typer{$typerSpec};
    }# for keyLev
}#configure_sort


#  Sort range [$wholeBeg, $wholeEnd) of @sortPerm for key level $kl,
#  When ties occur for numeric or alphabetical types, for each range of equal values, recurse to the next key level.
#  Recursion base case is either no more ties or the final key level.
sub sortKeylev_recurseOnTies{
    my( $kl,  #Key Level
        $wholeBeg, $wholeEnd )   = @_;
    state $collator=  Unicode::Collate->new();

    my $keys=  $sort_keyVec[$kl];   #with reference aliasing, perhaps could improve readability by something like \@key= $keyVec[$kl];

    #If pegging undef, move corresponding values of @sortPerm into @peggedPos.
    my @peggedPos;  #each element to hold the packed form of the row and permutation positions of one undef.
    if(  $sortPegUndefP[$kl]  ){
        defined $$keys[ $sortPerm[$_]]   or   push  @peggedPos,  pack 'LL', $_, $sortPerm[$_]
            for  $wholeBeg .. $wholeEnd-1;
        splice  @sortPerm, unpack('L'), 1   for reverse @peggedPos;
        $wholeEnd-= @peggedPos;
    }

    my $typer= $sortTyper[$kl];
    my $precs=  $sortPrecs[$kl];
    my @count= (0) x (1+$FIN);


    #Tally each type by precedence.
    for  my $j(  $wholeBeg .. $wholeEnd-1  ){
        my $type=   &$typer(  $$keys[ $sortPerm[$j] ]  );
        ++$count[  $$precs[ $type ]  ];
    }

    #Compute begs in order of precedence.
    my @beg;
    $beg[0 ]=  $wholeBeg;
    $beg[$_]=  $beg[$_-1] + $count[$_-1]   for 1..$FIN;

    #Initialize array to hold the positions in @sortPerm of the frontiers each type.
    my @grenze=  @beg;

    #Radix sort by type.
    for  my $blockPrec  (0..$FIN-1){
        my $elemPrec;
        for  my $j(  $grenze[$blockPrec] .. $beg[$blockPrec+1]-1   ){
            @sortPerm[  $j,  $grenze[$elemPrec]++   ]=   @sortPerm[  $grenze[$elemPrec],  $j  ]
                while   $blockPrec  !=  ($elemPrec=   $$precs[   &$typer($$keys[ $sortPerm[$j] ])   ]);
        }
    }
    
    #Reuse @grenze as arrary of ends.
    @grenze[0..$FIN-1]= @beg[1..$FIN];  $grenze[$FIN]= $wholeEnd;

    #Sort individual each type block on current key and recursively sort on next key level to break ties for equal value runs.
    for  my $blockPrec  (0..$FIN){
        my( $beg, $end )=   (  $beg[$blockPrec], $grenze[$blockPrec]  );

        next   if  $beg  >=  (my $fin= $end-1);                                          #Loop flow

        my $type=    &$typer(   $$keys[  $sortPerm[ $beg ]]   );
        if(   $type == $NAN   or  $type == $MIS   or  $type == $UND   ){
            if(  $kl == $#sort_keyVec  ){
                @sortPerm[$beg..$end-1]=  sort {$a<=>$b} @sortPerm[$beg..$end-1];}  #Make sort stable.
            else{
                sortKeylev_recurseOnTies   $kl+1,  $beg, $end;
            }
        }
        elsif(  $type eq $NUM  ){
            if(  $sortRevNumP[$kl]  ){  @sortPerm[$beg..$fin]=  sort  {$$keys[$b] <=> $$keys[$a]}   @sortPerm[$beg..$fin]   }
            else                     {  @sortPerm[$beg..$fin]=  sort  {$$keys[$a] <=> $$keys[$b]}   @sortPerm[$beg..$fin]   }

            for(   my $blockEnd= $end;  ($end= 1+$beg) <= $blockEnd;  $beg= $end   ){
                my $begVal=  $$keys[ $sortPerm[$beg] ];
                $end++     while   $end < $blockEnd   and   $$keys[ $sortPerm[$end] ]  ==  $begVal;
                if(  $end-$beg > 1  ){
                    if(  $kl == $#sort_keyVec  ){
                        @sortPerm[$beg..$end-1]=  sort {$a<=>$b} @sortPerm[$beg..$end-1];}  #Make sort stable.
                    else{
                        sortKeylev_recurseOnTies   $kl+1,  $beg, $end;
                    }
                }
            }
        }
        else{#  $type eq $ALF
            if(  $sortAlfT[$kl] == 0  ){
                @sortPerm[$beg..$end-1]=  sort {$a<=>$b} @sortPerm[$beg..$end-1];   #Restore order which radix sort may have changed.
                sortKeylev_recurseOnTies   $kl+1,  $beg, $end   if $kl < $#sort_keyVec;
                next;                                                                    #LOOP FLOW
            }
            $$keys[$_]=  $collator->getSortKey( $$keys[$_] )   for @sortPerm[$beg..$fin];
            ;  if(  $sortAlfT[$kl] == +1  ){
                @sortPerm[$beg..$fin]=   sort  {$$keys[$a] cmp $$keys[$b]}  @sortPerm[$beg..$fin];}
            elsif(  $sortAlfT[$kl] == -1  ){
                @sortPerm[$beg..$fin]=   sort  {$$keys[$b] cmp $$keys[$a]}  @sortPerm[$beg..$fin];
            }
            for(   my $blockEnd= $end;  ($end= 1+$beg) <= $blockEnd;  $beg= $end   ){
                my $begVal=  $$keys[ $sortPerm[$beg] ];
                $end++     while   $end < $blockEnd   and   $$keys[ $sortPerm[$end]]  eq  $begVal;
                if(  $end-$beg > 1  ){
                    if(  $kl == $#sort_keyVec  ){
                        @sortPerm[$beg..$end-1]=  sort {$a<=>$b} @sortPerm[$beg..$end-1];}  #Make sort stable.
                    else{
                        sortKeylev_recurseOnTies   $kl+1,  $beg, $end;
                    }
                }
            }
        }
    }#for $blockPrec

    if(  $sortPegUndefP[$kl]  ){   #Put positions pointing to undef back where they were.
        splice  @sortPerm, unpack('L'), 0,  unpack 'x4L'  for @peggedPos;
    }

}#sortKeylev_recurseOnTies


BEGIN{
    *CORE::GLOBAL::stat=  sub{ 
        _warn
            "stat command disabled.  Assuming you meant 'stats'."=>
            "statコマンドは使用不可になっている。'stats'のつもりだと仮定して進む。";
        goto &stats;
    }
}


__DATA__
=head1  NAME
 perltab -- perl tab

=head1  SYNOPSIS
 perltab  [opts] [TSV_FILE...]
 perltab  -h help


=head1  DESCRIPTION|説明
perltab is basically a fancy version of perl autosplit mode,
convenient for processing tab separated fields data.
perltabはperlのautosplitモードに毛の生えたような、タブ区切りファイルの単発な処理に便利なツール。


=head1  HELP|ヘルプ
=settab              ==
 perltab -h all|man  Prints entire documentation.
|                    説明書全体を表示。
 perltab -h kw       List keywords with documentation.
|                    説明書のキーワード・リストを表示。
 perltab -h opts     Describes command line options.
|                    コマンド行オプションを表示。
 perltab -h toc      Prints documentation table of contents.
|                    説明書の目録を表示。
 perltab -h KEYWORD  Prints part of documentation matching KEYWORD.
|                    説明書のKEYWORDに関連する部分を表示。
=unsettab
KEYWORD can be abbreviated; for example '-h star' instead of '-h starting'.  For Japanese, set environment variable LANG=ja_JP.utf8.
'-h starting'の代わりに'-h star'のようにKEYWORDを略しても良い。
'-h 始め'のように日本語のキーワードも略して使える。環境変数LANGをCなどに変更すれば、英語の説明文は表示される。


=head1  STARTING|始めに
To get started, make a sample input file.
まず、入力ファイルの例を作成し、

 % echo -e "FEATURE\tApple\tMikan\tPear\nWeight\t100\t100\t165\nCalorie\t52\t47\t57" > fruit.tsv

And try running perltab.
perltabを試してみる。

  % perltab fruit.tsv

This should do nothing if fruit.tsv is a proper tab separated field file.
fruit.tsvが各行の列数が揃っている正常なタブ区切りファイルでこのコマンドは何もしない。

Next try running perltab with some simple scripts.
次に簡単なスクリプトでperltabを試してみる。

  % perltab -e 'say "\@F: @F"'  fruit.tsv
  % perltab -H 'say "\@H: @H"' -d 'say "\@F: @F"'  fruit.tsv
  % perltab -e 'say $_,F($_) for @H'  fruit.tsv
  % perltab -H 'say $_,N($_) for @H'  fruit.tsv

Many more examples can be founds in EXAMPLES.
EXAMPLESにperltabの使用例が数多く挙げられている。


=head1  INPUT|入力
perltab assumes input files to be text files with fields (possibly empty) separated by single tabs.  Unless specified by command line option -N|--no-head, the first line of each input file is assumed to hold column labels.

An example input file:
入力ファイルの例、
  ------------------------------------
  FEATURE     Apple    Mikan    Pear
  Weight      100      100      165
  Calorie     52       47       57
  ====================================

The head line is:
先頭行は、
  'FEATURE     Apple    Mikan    Pear'

The data lines are:
データ行は、
  'Weight      100     100      165'
  'Calorie     52      47       57'

=kw missing|Missing values|欠損値
=head2  Missing values|欠損値

perltab defines functions, such as dit() and speak(), to facilitate files with missing values expressed as empty strings.
perltabは(空文字列で表した)欠損値を含むデータを扱いやすくするため、dit()やspeak()のような関数を提供している。
=wk missing|Missing values|欠損値

=head2  Mixed numerical and non-numerical values|数値と数値以外の値の混合

perltab defines functions, such as doNum, to facilitate handling columns with numberical entries mixed with non-numerical strings.
perltabはdoNumのような関数を提供することにより、数値と数値以外の値が混在する列を扱いやすくしている。

=kw ditto
=head2  ditto mode|同上モード

In ditto mode dittoMark strings are considered as to be like ditto marks, inheriting the value above them in their column.
同上モードでは入力にあるdittoMarkの文字列は欠損値ではなく、同じ列の上にある値の省略と見なさられる。

An example to illustrate ditto mode:
同上モードを説明するための入力例、
  <<  NEAsiaMegacities_ditto.tsv

  Population  Country    City
  36923000    Japan      Tokyo
  19342000               Osaka
  34000000    China      Shanghai
  25000000               Guangzhou
  24900000               Beijing
  23300000               Shenzhen
  25514000    Korea      Seoul


Option -D|--ditto affects both how input files are interpreted as read and the way they are ouput.
オプション-D|--dittoは入力の解釈と出力の形式に影響する。
-D|--dittoを指定すると同上モードはすべての列の入出力に適応される。

For example,
例えば、

 % perltab -pDd '$F[N(pop)]=2500_0000  if F(city) eq "Beijing"'

Makes the population of Beijing equal to Guangzhou producing:
は北京の人口と広州と等しくする為、以下の出力になる。

  Population  Country    City
  36923000    Japan      Tokyo
  19342000               Osaka
  34000000    China      Shanghai
  25000000               Guangzhou
                         Beijing
  23300000               Shenzhen
  25514000    Korea      Seoul


To convert a ditto format file to full format:
同上形式のファイルを普通の形式に変換するコマンド、
 % perltab -De 'say @F'  NEAsiaMegacities_ditto.tsv

To convert a full file to ditto format:
普通形式のファイルを同上形式に変換するコマンド、
 % perltab -pD  NEAsiaMegacities.tsv
=wk ditto


=head1  VARIABLES|変数
perltab provides the following column related variables:
perltabは列に関する以下の変数を提供する。

=kw @H
=settab        ==
 @H  $H[i] --> Label of ith column.
|              i番目の例ラベル
=wk @H
=kw @F
 @F  $F[i] --> Current value of ith column.  As in perl -a.
|              現在行のi番目の例の値。  perl -aと同様。
=wk @F
=kw @Fin
 @Fin          Copy of @F as read in.
|              読み込んだ時の@Fのコピー。
=wk @Fin
=kw prev|@FinPrev
 @FinPrev      @Fin from the previous row (initially the empty list).
|              ひとつ前の行の@Fin。初期値は空のリスト()。
=unsettab
=wk prev|@FinPrev


=head1  OPTIONS|オプション
=kw opts
=head2  Inline Script Options|コマンド行内スクリプト・オプション
=kw -b|--begin
=settab                      ==
  -b|--begin  SCRIPT         SCRIPT to execute before reading input.
|                            入力を読み込む前に実行するスクリプト。
=wk -b|--begin
=kw -b2|--begin2
 -b2|--begin2 SCRIPT         On second pass, SCRIPT to execute before reading input.
|                            ２回目の入力走査で、入力を読み込む前に実行するスクリプト。
=wk -b2|--begin2
=kw -d|--data-script         
  -d|--data-script  SCRIPT   SCRIPT to run on all but the header row.
|                            ラベル行(入力の先頭行)以外の行に対して実行するスクリプト。
=wk -d|--data-script         
=kw -d2|--data-script2
 -d2|--data-script2  SCRIPT  On second pass, SCRIPT to run on all but the header row.
|                            ２回目の入力走査で、ラベル行(入力の先頭行)以外の行に対して実行するスクリプト。
=wk -d2|--data-script2
=kw -e|--row-script
  -e|--row-script  SCRIPT    SCRIPT to run on each row.
|                            各行に対して実行するスクリプト。
=wk -e|--row-script
=kw -e2|--row-script2
 -e2|--row-script2 SCRIPT    On second pass, SCRIPT to run on each row.
|                            ２回目の入力走査で、各行に対して実行するスクリプト。
=wk -e2|--row-script2
=kw -H|--head-script
  -H|--head-script  SCRIPT   SCRIPT to run after reading head row, but before any data rows are read.
|                            先頭行を読み込んだ直後に実行するスクリプト。
=wk -H|--head-script
=kw -H2|--head-script2
 -H2|--head-script2 SCRIPT   On second pass, SCRIPT to run after reading head row, but before any data rows are read.
|                            ２回目の入力走査で、先頭行を読み込んだ直後に実行するスクリプト。
=wk -H2|--head-script2
=kw -s|--sort|ソート|整列|並べ替え
  -s|--sort[CMP]  SCRIPT     Sort rows by key(s) returned by SCRIPT for each row, according to CMP specification.  See -h sort for more.
|                            SCRIPTが行ごとに返すキーをCMPで指定した仕様で比較し、行を並べ替える。CMPの仕様について-h sortを参照。
=wk -s|--sort|ソート|整列|並べ替え
=kw -s2|--sort2|ソート|整列|並べ替え
 -s2|--sort2[CMP] SCRIPT     On second pass, sort rows by key(s) returned by SCRIPT for each row, according to CMP specification.  See -h sort for more.
|                            ２回目の入力走査で、スクリプトSCRIPTが行ごとに返すキーをCMPで指定した仕様で比較し、行を並べ替える。CMPの仕様について-h sortを参照。
=wk -s2|--sort2|ソート|整列|並べ替え
=kw -z|--end
  -z|--end= SCRIPT           SCRIPT to execute after processing all of the input.
|                            入力全体を処理した後に実行するスクリプト。
=wk -z|--end
=unsettab

=head2  Other Options|その他のオプション
=settab                      ==
=kw -c|--count               
  -c|--count                 Print number of rows for which the row script returned true (on final pass).
|                            (最終の入力走査で)行スクリプトが真を返した行数を出力する。
=wk -c|--count               
=kw ditto|-D|--ditto|ditto-mark
  -D|--ditto                 When an empty string column value is encountered, substitute value from previous row.
|                            入力ファイルに空文字列となっている列の値は上の行から取る。つまり空文字列を「同上」と見なす。
 --ditto-mark MARK           Like -D|ditto, but use MARK instead of empty string as ditto marker.
|                            -D|dittoと似ているか、入力ファイルにあるMARKに一致する値をその上の行から取る。
=wk ditto|-D|--ditto|ditto-mark
=kw -f|--script-file
  -f|--script-file FILE      Read scripts to execute in from FILE.
|                            実行すべきスクリプトをファイルFILEから読み込む。
=wk -f|--script-file
=kw -F|--field-sep|field-separator
  -F|--field-sep SEP         Input field separator SEP.  Default to tab.
|                            入力ファイルの列の区切り文字列SEP。デフォルト値はタブ。
=wk -F|--field-sep|field-separator
=kw -g|--grep
  -g|--grep                  grepMode: Print rows for which row script returns true (on final pass).
|                            grepMode: (最終の入力走査で)行スクリプトが真を返した行を出力する。
=wk -g|--grep
=kw -h|--help
  -h|--help [TOPIC]          Print help on TOPIC.  e.g. -h 'OPTIONS'. Default TOPIC is HELP.
|                            TOPICに関する説明を出力する。例えば、-h 'OPTIONS'。省略するとTOPICは'HELP'となる。
=wk -h|--help
=kw -N|--no-head|no-head|nohead
  -N|--no-head               Do not treat the first (non-comment) line of input files as a head line.
|                            入力ファイルの先頭を(列ラベルの行として)特別扱いしない。
=wk -N|--no-head|no-head|nohead
=kw --skipinf|inf|num|number|numeric|isnum|数値
 --skipinf                   Do not treat Inf values as numbers.  Affects behavior of numP() functions using numP() such as donum() and nums().
|                            Inf値を数値と見なさない。関数numP()及びdonum()やnums()のようなnumP()を使う関数に影響する。
=wk --skipinf|inf|num|number|numeric|isnum|数値
=kw -m|--missing-mark|missing|欠損値
  -m|--missing-mark          String used to mark used to mark missing, default to the empty string.
|                            欠損値の印となる文字列、デフォルト値は空文字列。
=wk -m|--missing-mark|missing|欠損値
=kw -M|--module
  -M|--module MODULES        MODULES to load before executing any scripts.
|                            スクリプトを実行する前にをロードしておくべきモジュール名。
=wk -M|--module
=kw --skipnan|nan|num|number|numeric|isnum|数値
 --skipnan                   Do not treat NaN values as numbers.  Affects behavior of numP() functions using numP() such as donum() and nums().
|                            NaN値を数値と見なさない。関数numP()及びdonum()やnums()のようなnumP()を使う関数に影響する。
=wk --skipnan|nan|num|number|numeric|isnum|数値
=kw -o|--output-file
  -o|--output-file FILE      Send output to file named FILE instead of STDOUT.
|                            標準出力に代わりにファイルFILEに出力する。
=wk -o|--output-file
=kw -OF|--OF|--output-field-sep
 -OF|--output-field-sep SEP  Output field separator SEP.  Defaults to the input field separator.
                             出力時の列の区切文字列SEP.  省略すると入力時の区切文字列は出力にも使われる。
=wk -OF|--OF|--output-field-sep
=kw -p|--print-each-row
  -p|--print-each-row        Print @F after executing (the final) row script on each row.  Any undef values in @F are quietly replaced with the missing value mark.
|                            (最終入力走査で)各行に対し行スクリプト実行後に@Fを出力する。  出力時に@Fにあるundef値は静か欠損値マーク(デフォルトは空文字列)に変えられる。
=wk -p|--print-each-row
=kw -n|--print-script
  -n|--print-script          Output script suitable for use with -f|--script-file.  See -h script-file.
|                            -f|--script-fileで使えるスクリプトを出力うる. -h script-fileを参照。
=wk -n|--print-script
=kw -q|--grep-quiet
  -q|--grep-quiet            perltab process returns 0 if -g option would have found any match rows.
|                            -gオプションの安静版。行スクリプトが真を返す入力行があれば、perltabプロセスはゼロを返す。
=wk -q|--grep-quiet
=kw -V|--version
  -V|--version               Print version number of perltab.
|                            perltabのバージョン番号を出力する。
=wk -V|--version
=kw -X|extra-fields|--extra-fields-OK|columns
  -X|--extra-fields-OK       Extra fields allowed in input lines.  Suppresses fatal error normally thrown when some input lines have more fields than expected.  The extra fields are included in @F; otherwise behavior under this option is not well defined.
|                            入力ファイルに余分な列(第１行より多い数の列)があってもエラーとして停止せずにその列を適当に@Fに入れて実行を続行する。ただし、列の数が合わないことにより他のエラーが発生する恐れはある。
=wk -X|extra-fields|--extra-fields-OK|columns
=unsettab
=wk opts


=head1  FUNCTIONS|関数
Most perltab functions fall into two categories: 1. accessors, 2. column manipulation functions, and 3. functions to help handle data of mixed type (i.e. numerical and missing value mark).
perltabが提供する関数の多くは、1.アクセサー的な関数、2. 列を動かしたりする関数  3.異なるタイプの値(数値と欠損値の標など)が混在する時に便利な関数。

=head2  Accessors|アクセサー的な関数
=kw ditto|dittoMark
=settab            ==
 sub dittoMark()   Value of string used to mark repeated values in ditto mode.  Defaults to the empty string.  See -h -D.
|                  同上モードで繰り返された値の代表として使われている文字列。デフォルト値は空文字列。-h -dを参照。
=wk ditto|dittoMark
=kw missingMark|missing|欠損値
 sub missingMark() Value of string used to represent missing values.  Defaults to the empty string.  See -h -m.
|                  欠損値を表す文字列。デフォルト値は空文字列。-h -mを参照。
=wk missingMark|missing|欠損値
=kw F
 sub F( LABELS )   Return value of column(s) given by LABELS.
|                  列ラベルLABELSの値を返す。
=wk F
=kw N
 sub N( LABELS )   Return column numbers of LABELS.
|                  列ラベルLABELの列番号を返す。
=unsettab
=wk N

=kw F|N
The relationship between @F, F() and N() is illustrated by the following equivalences;
F(), N(), @Fの関係は以下の等価式で表される。
   F( $H[i] )  ==  $F[   i      ]
   F( LABEL )  ==  $F[ N(LABEL) ]
where i is an index into @F, and LABEL is a column label.
ここで、iは@Fの添字となる整数で、LABELは列ラベルである。

For user convenience, scripts using F() and N() are 'doctored' are follows.
使い勝手を良くする為、F()とN()を含むスクリプトは自動的に以下の正規表現で書き直される。

  s{\bF\(  ( [^"',(){}\$@]+ )  \)} {F(qw($1))}gx;
  s{\bN\(  ( [^"',(){}\$@]+ )  \)} {N(qw($1))}gx;

For example, F(cat dog) and N(cat dog) are automatically converted to F(qw(cat dog)) and N(qw(cat dog)) respectively.
例えば、「F(cat dog)」や「N(cat dog)」は自動的に「F(qw(cat dog))」や「N(qw(cat dog))」のように書き換えられる。

=wk F|N
=kw label|列ラベル|ラベル|F|N
F() and N() also match unambiguous prefixes of the column labels, so one can replace this command:
F()とN()は列ラベルのみならず、(他のラベルと重複しない限り)ラベルの前置詞も受け入れるので、
 % perltab -e 'say N(height)'

with the shorter:
の代わりに、
 % perltab -e 'say N(hei)'

as long as no other column label starts with 'hei'.
などが使える。

Downcased unique prefixes are also matched.  Consider an input file with a column label 'FreddieMercury' and another labeled 'FredAstaire', but no other labels beginning with 'f' or 'F'.  The column 'FreddieMercury' could be abbreviated in many ways.  F(FreddieM), F(freddiem), F(fredd) are all okay.  F(freddieM), F(Freddiem) are not okay, because they are not prefixes of 'FreddieMercury' nor 'freddiemercury'.  F(Fred), F(fred) are not okay, because these also prefix match 'FredAstaire' or 'fredastaire'.
列ラベルの小文字化した前置詞も%Nに格納され、関数F()も同様の省略をサポートする。例えば、「FreddieMercury」と「FredAstaire」という列ラベルがあるが、他にFかfから始まる列ラベルがない入力ファイルの場合、FreddieMercuryはいくつかの省略の仕方が使える。F(FreddieM)、F(freddiem)、F(fredd)はぞれぞれ問題なく使える。しかし、F(freddieM)、F(Freddiem)は大文字が中途半端に入っており、「FreddieMercury」にも「freddiemercury」にも前置詞として一致しない。また、F(Fred), F(fred)は逆に「FreddieMercury」のみならず「FredAstaire」にも一致してしまうから使えない。
=wk label|列ラベル|ラベル|F|N


    Accessors (cont.)
    アクセサー的な関数(の続き)

=kw NR
=settab      ==
 NR          The number of times the row script has been called (on current pass).  Like Awk's NR, but counts from zero.
|            行スクリプトを呼び出している回数 (現行の入力呼び込みで)。 awk言語のNRと似ているが、ゼロから数える。
=wk NR
=kw NRT
 NRT         The number of rows (including initial label row if a script was run on it) for which the row script returned true (on current pass).
|            (現在の実行passで)行スクリプトが真を返した行数。Number Rows Trueの略。
=wk NRT

=kw stanza
 stanza     In numerical context returns the stanza number.  In string context returns the header comments lines for the current stanza.
|           数値文脈で呼ばれたら、スタンザ番号を返す。文字列文脈で呼ばれたら、スタンザの先端にあるコメント行を返す。
=unsettab

A stanza is a section of rows separated by blank lines.
スタンザとは空行で区切られた、連続した行のブロック。

=kw -g|--grep
 For example given input:
 例えば、以下の入力を与えられたら、

  ------------------------------------------------
  ;Opening comment followed by a blank line.

  FEATURE  apple   mikan   pear    peach    banana

  ; Numerical features
  weight   100     100     165     175      183
  cost     300     347     800     500      218
  height   5       3       5       4        7
  calorie  52      47      57      39       89

  ; Translations
  French   pomme   mikan   poire   pêche    banane
  German   Apfel   Mikan   Birne   Pfirsich Banane
  ================================================

Both:
で、

 % perltab -ge 'stanza == 2'  input.tsv

and:
又は、

 % perltab -ge 'stanza=~/Trans/'  input.tsv

Should yield:
の両方のコマンドはいづれも以下の出力が得られる。

  ------------------------------------------------
  ;Opening comment followed by a blank line.

  FEATURE  apple   mikan   pear    peach    banana

  ; Translations
  French   pomme   mikan   poire   pêche    banane
  German   Apfel   Mikan   Birne   Pfirsich Banane
  ================================================

By the way, under the -g|--grep option if any rows are printed the opening comments will be as well, and if any rows in a stanza are printed so will the stanza heading comments.  Continuing our example, the output of:
ちなみに-g|--grepオプション下ではひとつのデータ行でも出力されれば、入力の先頭コメントも表示される。また各節において、その節のひとつの行でも出力されれば節の上部コメントも表示される。

 % perltab -ge '$x= F(app); num $x and $x > 99'

  ------------------------------------------------
  ;Opening comment followed by a blank line.

  FEATURE  apple   mikan   pear    peach    banana

  ; Numerical features
  weight   100     100     165     175      183
  cost     300     347     800     500      218
  ================================================

If you do not like the comments, an easy way to remove them is:
このコメントを除きたければ、以下のコマンドは使える。

 % perltab -ge '...' |  sed -ne '/^[^;]/ p'

Or to remove only the opening comment at top:
或は入力先頭のコメントのみを除くには以下のコマンドが便利。

 % perltab -ge '...' |  sed -ne '/^[^;]/,$ p'
=wk -g|--grep
=wk stanza

=head2  Column Manipulation Functions|列を動かしたりする関数

=kw del|delete|削除
 sub del(  COLUMNS  )

 Delete COLUMNS from @F.  Columns may be given as labels or integers in [-$#F,$#F].
 列COLUMNSを@Fから削除する。列はラベル、又は[-$#F,$#F]内の整数で指定できる。

=settab                   == 
 del  0;                  #Delete first column.
|                         #一番目の列を削除
 del -2;                  #Delete penultimate column.
|                         #最後列の前の列を削除
 del qw(weight height);   #Delete columns 'weight' and 'height'.
|                         #'weight'と'height'の列を削除する
=unsettab
=wk del|delete|削除

=kw insert|挿入
=kw insaft
 sub insAft (  PRE, *columnInfo*  )
 sub insaft (  PRE, *columnInfo*  )

 Insert new column(s) into @F after column PRE.  PRE may be given by label or as an integer in [-$#F,$#F].

 新しい列をPREの後に加える。PREはラベル、又は[-$#F,$#F]内の整数で指定できる。

 In general (without the -N|--no-head option),
 一般に(-N|--no-head)オプションを指定していない場合、
 
 sub insAft(  PRE,  LABEL1, ...,  DATA_GEN1, ...  )
 sub insaft(  PRE,  LABEL1, ...,  DATA_GEN1, ...  )

 The non-head rows of the new colum LABEL1 are given by DATA_GEN1.  If DATA_GEN1 is a code reference it is executed and the return value used.  Any number of (LABEL, DATA_GEN) pairs may be stipulated.
 新しい列LABEL1の（ラベルを格納する先頭行以外）の内容はDATA_GEN1で指定する。DATA_GEN1がsubへの参照子である場合、DATA_GEN1を実行した返り値が列の内容となる。(LABEL, DATA_GEN)２組の数に特に制限はない。

 Insert BMI column before column 'weight'.  'ih' is used to skip the numerical computation on the head line.
 列'weight'の後に新しい'BMI'列を加える。ihは先頭行でBMIの数値計算を避ける為。

=settab    ==
 insAft(  'weight',  'BMI',  (ih or  $F{weight} / $F{height}**2)  );
=unsettab

 Or,
 又は、
=settab    ==
 insAft(  'weight',  'BMI'=> sub{ 10000*$F{weight} / $F{height}**2}  );
=unsettab

 Note that 'ih or' is not needed when DATA_GEN is a code reference.
 DATA_GENがsubへの参照子の場合、'ih or'は不要。

 The same, but robust to empty string missing values.
 同じ計算をするが、空の文字列などで表した欠損値に頑健。
=settab    ==
 insaft(  'weight',  'BMI'=> sub{ doNum{10000*$_[1]/$_[0]**2}, @F{height weight} }  );
=unsettab

 But when -N|--no-head option is stipulated,
 ただし、-N|--no-headオプションが指定された場合、

 insAft(  SUC,  DATA_GEN1, ...,  )
 insaft(  SUC,  DATA_GEN1, ...,  )

 In this case, no need to deal with labels or the head line.
 この場合、ラベルや先頭行の処理は不要。
=wk insaft

=kw insbef
 sub insBef(  SUC, *columnInfo*  )
 sub insbef(  SUC, *columnInfo*  )

 Insert new columns into @F before column SUC.  SUC may be given by label or as an integer in [-$#F,$#F].  See insAft for more details.
 新しい列をSUCの前に加える。SUCはラベル、又は[-$#F,$#F]内の整数で指定できる。詳細はinsAftを参照。
=wk insbef
=wk insert|挿入

=kw move|移動
=kw movaft
 sub movAft(  PRE,  @colsToMov   )
 Move columns @colsToMov after column PRE.  Columns may be given as labels of integers in [-$F#,$#F].
 列@colsToMovを列PREの後に移す。列はラベル、又は[-$#F,$#F]内の整数で指定できる。
=wk movaft

=kw movbef
 sub movBef(  SUC,  @colsToMov   )
 Move columns @colsToMov before column SUC.  Columns may be given as labels of integers in [-$F#,$#F].
 列@colsToMovを列SUCの前に移す。 列はラベル、又は[-$#F,$#F]内の整数で指定できる。
=wk movbef
=wk move|移動

=kw swap|入れ換え
 sub swap( F1,F2,...,Fn, G1,G2,...,Gn )
 Swaps columns F1 and G1, then columns fields F2 and G2,...   Columns may be given as labels of integers in [-$F#,$#F].
 列F1とG1を入れ換え、それから列F2とG2を入れ換え...。列はラベル、又は[-$#F,$#F]内の整数で指定できる。
=wk swap|入れ換え

=head2  Value Filters|値フィルター
=kw valuefilters|value filters|値フィルター
=kw deq
 sub deq( STRING1, STRING2... )
 Defined and EQual.  If any arguments are undefined return undef.  Otherwise return 't' if all arguments are string equal and '' if not.
 If all arguments are string equal return 't'; else if any argument is undefined return undef, otherwise return ''.
 Mnemonic: Defined and EQual.
 引数がすべて同じ文字列ならば't'を返す、そうでなければ引数にundefがあればundefを返し、なければ''を返す。覚え方、Defined and EQual。
=wk deq

=kw dit
 sub dit( @STRING )
 Like say( @STRING ), but replaces undef arguments with the missing value mark.  Mnemonic: French for say, starts with 'd' as does defined.
 普通はsay( @STRING )と同じであるが、値がundefの要素は欠損値の文字列として表示する。。
=wk dit

=kw speak
 sub speak( @STRING )
 Like say( @STRING ), but does nothing if @STRING is empty or contains any empty strings.  Mnemonic: Speeches should be grand with no short elements.
 普通はsay( @STRING )と同じであるが、@STRINGが空である、もしくは空の文字列を含んでいれば、speakは何もしない。
=wk speak

=kw form|format
 sub form( FORMAT, STRING1... )
 Return STRING1... sprintf formated according to FORMAT.  When FORMAT expects a number (e.g. '%3d'), non-numbers are passed through as is.
 文字列STRING1...にsprintf( FORMAT, STRING1 )をかけ、その結果を返す。FORMATが（例えば'%3d'）のような数値を前提にしている場合、数値以外の値にはsprintfをかけずにそのまま返す。
=wk form|format

=kw reform|format
 sub reform( FORMAT, STRING1... )
 Caution, modifies STRING1... according to sprintf format FORMAT.  When FORMAT expects a number (e.g. '%3d'), non-numerical STRINGs are left unchanged.
 注意、STRING1...の値を変えてしまう。文字列STRING1...をsprintf形式のFORMATに従って変更する。FORMATが（例えば'%3d'）のような数値を前提にしている場合、数値以外のSTRINGはそのまま残る。
=wk reform|format

=kw nomt|noempty|no empty|empty
 sub nomt( @STRING )
 Returns true if @_ contains no empty strings.
 @STRINGに空の文字列が含まれていなければ真を返す。
=wk nomt|noempty|no empty|empty

=kw an|any|non-empty
 sub an( @LIST )
 Returns non-empty strings in @LIST.
 Named for the construct:{  say @x  if an @x  }.
 @LIST中の非空文字列を返す。
 「an」という名称は、{  say @x  if an @x  }の使い方に由来する。
=wk an|any|non-empty

=kw numeric|数値
=kw --inf|inf|--nan|nan|num|number|isnum
=kw nump
 sub numP( ARG )
 Returns true if ARG looks like a number.  $_ used when ARG is omitted.
 Inf and NaN values are not treated as numbers when the respective --skipinf or --skipnan options are given.
 ARGが数値に見える場合、真を返す。ARGを省略すると$_が使われる。
 --skipinfと--skipnanのオプションを指定するとInf値とNaN値はそれぞれ数値として扱われない。
=wk nump

=kw num0
 sub num0( ARG )
 Returns ARG if it looks like a number, otherwise 0.  $_ used when ARG is omitted.
 ARGが数値に見えればARGを返し、そうでなければ０を返す。ARGを省略すると$_が使われる。
=wk num0

=kw num1
 sub num1( ARG )
 Returns ARG if it looks like a number, otherwise 1.  $_ used when ARG is omitted.
 ARGが数値に見えればARGを返し、そうでなければ１を返す。ARGを省略すると$_が使われる。
=wk num1

=kw allnum
 sub allNum( @ARG )
 sub allnum( @ARG )
 Returns true if all elements of @ARG look like numbers.
 @ARGの要素がすべて数値に見える場合、真を返す。
=wk allnum

=kw donum
 sub doNum( {CODE} val... )
 sub donum( {CODE} val... )
 If all vals look numerical, run CODE and forward its return value, otherwise returns the missing value mark (defaulting to the empty string if not set).
 val...が全て数値に見える場合、{CODE}を実行した返り値を返すが、
 val...の内に数値に見えない値があれば、{CODE}を実行せずに欠損値の印(デフォルト値は空文字列)を返す。
=wk donum

=kw nums
 sub nums( @ARG )
 Returns numerical elements of @ARG.
 @ARGに含まれている数値を返す。
=wk nums
=wk valuefilters|value filters|値フィルター
=wk --inf|inf|--nan|nan|num|number|isnum

=kw numerical functions
=head2  Numerical Functions|数値演算関数

=kw bemax|bemin|beminmax
 sub bemax( @VAL )
 Set first element of @VAL to maximum value of @VAL.  undef values are treated as smaller than any number, other non-numerical elements are quietly skipped.
 @VALの先頭要素を@VALの最大値にする。undefは無限に小さい値として扱われ、その他の数値でない要素は無視される。

 sub bemin( @VAL )
 Set first element of @VAL to minimum value of @VAL.  undef values are treated as larger than any number, other non-numerical elements are quietly skipped.
 @VALの先頭要素を@VALの最小値にする。undefは無限に大きい値として扱われ、その他の数値でない要素は無視される。

 sub beminmax( @VAL )
 Set first  element of @VAL minimum as in bemin( @VAL ), and second element of @VAL maximum as in bemax( @VAL ).
 @VALの先頭要素をbemin( @VAL )のように最小値にし、@VALの２番目の要素をbemax( @VAL )にする。
=wk bemax|bemin|beminmax

=kw bigger
 sub bigger( STRING1, STRING2 )
 Returns iff string STRING1 would be preferable to STRING2 when looking for large numbers.  Tries to handle special numerical values (e.g. 'inf') and non-numerical strings in a robust way.  For example bigger( 'dog', 5) returns false.
 大きい数値を探している場合、STRING1がSTRING2より良ければ、真を返す。infのような特殊な数値や非数値の値を渡されてもエラーを投げずに適切な返り値を返す。例えば、bigger( 'dog', 5 )は偽を返す。
=wk bigger
=kw lesser
 sub lesser( STRING1, STRING2 )
 Returns iff string STRING1 would be preferable to STRING2 when looking for small numbers.  Tries to handle special numerical values (e.g. 'inf') and non-numerical strings in a robust way.  For example bigger( 'dog', 5) returns false.
 小さい数値を探している場合、STRING1がSTRING2より良ければ、真を返す。infのような特殊な数値や非数値の値を渡されてもエラーを投げずに適切な返り値を返す。例えば、bigger( 'dog', 5 )は偽を返す。
=wk lesser
=wk numeric|数値
=wk numerical functions

=head2  Statistical Functions|統計関数
=kw statistics|統計
=kw sum|mean|average|geomean|geometric mean|stdev|qtiles|quantiles|quartiles|percentiles|stats|和|平均|平均値|幾何平均|幾何平均値|標準偏差|分位値|分位数
The statistical functions: sum, mean, geomean, stdev, qtiles, xqtiles, stats; share some properties.
 *  They ignore non-numerical data values.
 *  Calling them with non-empty data has the side effect of setting an internal array.
 *  When called with an empty data [e.g. mean()], if the internal array has been set, they use those values.
 *  They die with error message if no numerical values are available to perform their operations.
 *  They try to handle special numerical values such as {inf, -inf, nan, -NaN} reasonably.
統計関数、sum, mean, geomean, stdev, qtiles, xqtiles, statsは以下の共通点を持つ。
 •  データに含まれる数値以外の値は無視する。
 •  副作用として渡されたデータを内部配列に保存しておく。
 •  データ無しで呼ばれたら[例えば、mean()]、内部配列に保存してあるデータを使う。
 •  {inf, -inf, nan, -NaN}のような特殊の数値を適切に処理するように努力する。
=wk sum|mean|average|geomean|geometric mean|stdev|qtiles|quantiles|quartiles|percentiles|stats|和|平均|平均値|幾何平均|幾何平均値|標準偏差|分位値|分位数


=kw sum|和
 sub sum( @VAL )
 Returns the sum of the numerical values in @VAL.
 @VALに含まれている数値の和を返す。
=wk sum|和

=kw mean|average|平均|平均値
 sub mean( @VAL )
 Returns the mean of the numerical values in @VAL.
 @VALに含まれている数値の平均値を返す。
=wk mean|average|平均|平均値

=kw geomean|geometic mean|average|幾何平均|幾何平均値
 sub geomean( @VAL )
 Returns the geometic mean of the numerical values in @VAL.
 @VALに含まれている数値の幾何平均値を返す。
=wk geomean|geometic mean|average|幾何平均|幾何平均値

=kw stdev
 sub stdev( @VAL )
 Returns the mean of the numerical values in @VAL.
 @VALに含まれている数値の標準偏差を返す。
=wk stdev

=kw qtiles|xqtiles|quantile|median|quartile|percentile|分位値|分位数|中央値|４分位値|四分位値|４分位数|四分位数|百分位値|パーセンタイル
 sub qtiles( Q, @VAL ),   xqtiles( Q, @VAL )
 Return  Q-quantile of @VAL.  xqtiles includes minimum and maximum values as well.
 For example,  qtiles( 2,  1..5 ) --> returns   (3).
              xqtiles( 2,  1..5 ) --> returns (1,3,5).
 @VALのQ分位値を返す。xqtilesの場合は、Q分位値に最小値と最大値も含める。
 例えば、 qtiles( 2,  1..5 )は  、(3)  を返すが、
 例えば、xqtiles( 2,  1..5 )は、(1,3,5)を返す。
=wk qtiles|xqtiles|quantile|median|quartile|percentile|分位値|分位数|中央値|４分位値|四分位値|４分位数|四分位数|百分位値|パーセンタイル

=kw stats|mean|stdev|median|quartile|平均値|標準偏差|中央値|４分位値|四分位値|４分位数|四分位数
 stats( @VAL )
 Returns a string containing descriptive stats of @VAL.
 @VALのいくつかの代表値を含んだ文字列を返す。
=wk stats|mean|stdev|median|quartile|平均値|標準偏差|中央値|４分位値|四分位値|４分位数|四分位数
=wk statistics|統計

=head2  Statistical Functions|統計関数
=kw math functions|数学関数
=kw log|log2|log10|logarithm|対数
 log2( @VAL )
 log10( @VAL )
 Return log to the base 2 or 10 of values in @VAL.  Like Perl log(), $_ is used when @VAL is omitted.
 はそれぞれ、@VALの数値要素の２(もしくは10)を底にした引数の対数を返す。Perlのlog()関数と同様、$VALを省略すると$_が使われる。

 log2 and log20 differ from Perl log() in these respects:
 *  A list of arguments @VAL may be given.
    (Dies with error message when called in scalar context if @VAL has more than one element.)
 *  Non-numerical values are passed through unchanged.
 *  When passed integer powers of 2(10) integer values,
    so ++$freq[log2 8] and ++$freq[log10 1000] are equivalent to ++$freq[3].

 log2及びlog10の挙動は以下の点でPerl log()と異なる、
 ・ リストの引数@VALを受け付ける。
    (スカラ文脈で呼ぶ場合、@VALの引数の個数がふたつ以上ならperltabはエラー文を吐いて終了する。)
 ・ 数値でない値はそのまま通される。
 ・ @VALの値が整数かつ２(10)の冪数であれば、整数を返す。
    例えば、++$freq[log2 8]と++$freq[log10 1000]は++$freq[3]と同じ。
=wk log|log2|log10|logarithm|対数
=wk math functions|数学関数


=head2  Other Functions|その他の関数

=kw Dumper|dump
 Invokes Data::Dumper::Dumper.
 Data::Dumper::Dumperを呼び出す。
=wk Dumper|dump

=kw ih|inhead|in head
 sub ih()
 Returns true iff the current line is the head line.  Mnemonic for 'In Head' or 'Is Head'.
 現在行が先頭行であれば真を返す。"ih"はIn HeadやIs Headの略。

 For example, to upper case the labels, but leave other rows unchanged, one might use:
 例えば、先頭行のラベルを大文字にするには、
 % perltab -pe  '@F= map {uc} @F  if ih'  input.tsv

 instead of using -H,
 は-Hを利用する代わりになる。
 % perltab -pH  '@F= map {uc} @F'  input.tsv
=wk ih|inhead|in head

=kw SNset
 sub SNset( \% $ )
 SNset %S, PATHNAME
 Make %S into a Serial Number set, by adding (LINE=>LINE_NUM) entries to %S, LINE_NUM starts at one.  LINE is the (chomped) LINE_NUMth line of the file.  Reads keys in from stdin if PATHNAME is '-' or 'stdin' (case insensitive).  Mnemonic, both senses of the word set (to assign, and a set of items) apply.
 %Sに(LINE=>LINE_NUM)を加え、入力ファイルの行と通し番号(Serial Number)の集合にする。LINE_NUMは１から数えられ、LINEは入力ファイルのLINE_NUM番目の(chompした)行。PATHNAMEが'-'か'stdin'(や'STDIN')の場合、集合の要素は標準入力から読み込まれる。覚え方、ここでsetは英単語としての両方の意味(代入と集合)が当てはまる。
=wk SNset

=kw push2
  push2( @a, @b, $val1, $val2 )
 is a shortcut for:
 は、
  push( @a, $val1 );  push( @b, $val2 )
 の略。以下の例のように$val1, $val2が@valに入っている場合は特に便利。
 especially handy when $val1, $val2 are the elements of an array @val.
  push2( @a, @b, @val )  <-->   push( @a, $val[0] );  push( @b, $val[1] )
=wk push2

=kw push3
  push3( @a, @b, @c, $val1, $val2, $val3 )
 is a shortcut for:
 は、
  push( @a, $val1 );  push( @b, $val2 );  push( @c, $val3 )
 の略。以下の例のように$val1, $val2, $val3が@valに入っている場合は特に便利。
 especially handy when $val1, $val2, $val3 are the elements of an array @val.  Compare these:
  push3( @a, @b, @c, @val );  #Push one value each onto @a, @b, and @c.
=wk push3


=head1  EXAMPLES|使用例

 Output columns 'height' and 'weight'.
 身長と体重の列を表示する。
 % perltab -e 'say F(qw(height weight))' heightWeight.tsv
 or more succinctly by letting perltab add the qw for you (see -h qw).
 又は、qw()の省略し、perltabにそれを自動的に付け加えてもらう(-h qw を参照)。

 % perltab -e 'say F(height weight)' heightWeight.tsv
 or even more succinctly,
 又は、更に短く、
 % perltab -e 'say F(hei wei)' heightWeight.tsv
 or
 又は、
 % perltab -e 'say F(h w)' heightWeight.tsv
 Any unique prefix of the desired column labels will work (see -h label).  Here we assume no other headers starts with 'h' or 'w'.
 列ラベルは一意的に決まる限り、ラベルの前置詞で略しても良い(-h labelを参照)。ここでは'h'と'w'から始まる列ラベルはそれぞれ'height'と'weight'のみと仮定した。

=kw grep|@H
 This command uses grep and the @H array to print columns whose labels match a regular expression.
 このコマンドは、grepと列ラベルを格納している@H配列を利用し、ある正規表現に合うラベルの列を表示する。
 % perltab -e 'say F(grep /eigh/, @H)' heightWeight.tsv
=wk grep|@H

=kw an|any|non-empty
 Print height, weight and shoe size columns of rows for which at least one of those has a value.
 列height、weight及びshoe sizeのいづれかが非空の値を持つ行に対し、その３つの列を表示する。
 % perltab -e '@x= F(hei wei shoe); say @x if an @x'  heightWeightShoesize.tsv
=wk an|any|non-empty

=kw search
 Print row ID and column label of NaN data entries.
 NaN値の行IDと列ラベルを表示する。
 % perltab -e '$F[$_]=~ /^nan/i  and  say F(ID), $H[$_]  for 0..$#F'  input.tsv
=wk search

=kw swap
 Swap first and final field.
 最初と最後の列を交換する。
 % perltab -pe 'swap 0, -1'  input.tsv
=wk swap

 Swapping columns can also be done (more efficiently) by directly accessing @F.
 列の交換には@Fを直接扱う、(より効率的な)方法もある。
 % perltab -e 'say  @F[-1, 1..$#F-1, 0]'  input.tsv

 Swap fields labeled 'height' and 'weight'.
 heightとweightの列を交換する。
 % perltab -pe 'swap qw(height weight)'  heightWeight.tsv

 Swapping columns by label can also be done (more efficiently) by using N() to access @F directly.
 ラベルで指定した列の交換にはN()を利用して@Fを直接変更する、(より効率的な)方法もある。
=settab    ==
 % perltab -pe '@I= N(height weight); @F[@I]= @F[reverse @I]'  heightWeight.tsv
=unsettab

=kw -F|-OF|--OF|--field-sep|--output-field-separator|convert|csv|カンマ区切り
 Convert comma separated fields file into tab separated fields format.
 カンマ区切り形式ファイルをタブ区切り形式に変換する。
 % perltab -pF=',' -OF=$'\t'  input.csv
=wk -F|-OF|--OF|--field-sep|--output-field-separator|convert|csv|カンマ区切り

=kw merge|dit|-in2
 Given two tsv files:  height.tsv, weight.tsv  with some common IDs but possibly in a different order and only partially overlapping, output a table with column 'weight' from weight.tsv appended onto the table in height.tsv.  The new column will hold the missing value mark for any IDs in height.tsv but absent in weight.tsv.
 IDが一部共通するが行の順番が異なるweight.tsvとheight.tsvという２つのタブ区切りファイルを与えられたら、weight.tsvのweight列を(height.tsv中のID順に従って)最後の列としてweight.tsvに加えた表を出力する。height.tsvのIDの内weight.tsvにないIDがある場合そのIDのweightの値は欠損値として扱う。
=settab     ==
 % perltab  -e '$V{ F(ID)}= F(weight)'  -e2 'dit @F, $V{ F(ID)}' weight.tsv  -in2 height.tsv
=unsettab
 An alternative way to do this is:
 もうひとつの方法、
=settab     ==
 % perltab  -e '$V{ F(ID)}= F(weight)'  -pe2 'insaft -1, "weight", $V{ F(id)} //""'  weight.tsv  -in2 height.tsv
=unsettab

  weight.tsv:         height.tsv:
  ----------------    ----------------
  ID  weight              ID  height
  1   73                  3   160
  2   78                  2   175
  5   55                  6   180
  3   0                   1   170
  ================    ================

  output:
  -------------------------
  ID  height  weight
   3   160     0
   2   175     78
   6   180     
   1   170     73
  =========================
=wk merge|dit|-in2

 Move column 'ID' to directly after column 'weight'.
 列IDを列weightの後に移動する。
 % perltab -pe 'movaft qw(weight ID)'  heightWeight.tsv

 Manually move column 'ID' to directly after column 'weight', assuming column ID comes before colun weight in the input.
 手動で列IDを列weightの後に移動する。入力におけるIDがweightの前に来る前提の場合。
=settab    ==
 % perltab -e '($x,$y)= N(ID weight);  say @F[0..$x-1,$x+1..$y,$x,$y+1..$#F]'   heightWeight.tsv
=unsettab

=kw del|grep|@H
 Delete columns starting in 'c', 'd' or 'e'.
 c,d,eのいづれかから始まるラベルを削除する。
 % perltab -pe 'del grep /^[cde]/, @H'  animals.tsv
=wk del|grep|@H

=kw speak|missing
 Echo input file, skipping data lines with any missing (empty string) values.
 欠損値(空文字列)を含むデータ行を飛ばしながら入力ファイルを表示する。
 % perltab -e 'speak @F'  heightWeight.tsv
=wk speak|missing

 Output BMI (Body Mass Index) from rows in which both height and weight have values.
 身長と体重の値がふたつとも記録してある行の体格指数"BMI"を表示する。
=settab    ==
 % perltab -e 'donum{say 10000*$_[1]/$_**2} F(height weight)'  heightWeight.tsv
=unsettab

=kw insaft
 Add BMI (Body Mass Index) column to table in heightWeight.tsv, skipping missing (empty string) values.  New column is placed after "weight" column.
 欠損値(空文字列)を飛ばしながら、体格指数"BMI"の列をheightWeight.tsvの表に加える。新しい列はweight列の後に入る。
=settab    ==
 % perltab -pe 'insAft  qw(weight BMI)=> doNum{10000*$_[1]/$_**2} F(height weight)'  heightWeight.tsv
=unsettab
=wk insaft

=kw reform|format
 Reformat numbers in field 'BMI'.
 列BMI中の数字の表示形式を変える。
 % perltab -pe 'reform "%.2f", F(BMI)'  heightWeightBMI.tsv

=kw -N|--no-head|nohead|-F|--field-sep|field-separator
 Given data from a LaTeX table, separated by ' & ', and with no head line; reformat the numbers in columns 2-4.
 LaTeX表のデータ(ラベル行無しで、列は' & 'で区切られている形式)で２〜４番号の列中の数値の小数点以下の桁数を変更する。
 % perltab -N -F=' & '  -pe 'reform "%5.3f", @F[2..4]'  tableBody.tex
=wk -N|--no-head|nohead|-F|--field-sep|field-separator
=wk reform|format

=kw number|numeric|isnum|数値
=kw speak|nums
 One way to print data in column 1 for rows in which it has a numerical value.
 列１のデータの内、数値となっている行を表示する方法のひとつ。
 % perltab -d 'speak nums $F[1]'  heightWeight.tsv
=wk speak|nums

=kw donum
 One way to print data in column 1 for rows in which it has a numerical value.
 列１のデータの内、数値となっている行を表示する方法のひとつ。
 % perltab -d 'donum {say} $F[1]'  heightWeight.tsv

=kw --skipinf|--skipnan|inf|nan
 Print data in colums 1,2 for rows in which both have a numerical value.  Special values {'inf', '-inf', 'nan', '-NaN'}, etc. are treated as numbers.
 列1,2のデータの内、両方が数値となっている行を表示する。{'inf', '-inf', 'nan', '-NaN'}のような特殊な値を数値と見なす。
 % perltab -d 'donum {say @_} @F[1,2]'  heightWeight.tsv

 As above, print data in colums 1,2 for rows in which both have a numerical value; but do not treat {'inf', '-inf', 'nan', '-NaN'} etc. as numbers.
 上記のコマンドと同様、列1,2のデータの内、両方が数値となっている行を表示するが、{'inf', '-inf', 'nan', '-NaN'}のような特殊な値を数値と見なさない。
 % perltab --skipinf --skipnan -d 'donum {say @_} @F[1,2]'  heightWeight.tsv
=wk --skipinf|--skipnan|inf|nan
=wk donum
=wk number|numeric|isnum|数値

 Display column labels with their numbers.
 タブ区切りファイルのラベルとその番号を表示する。
 % perltab -H 'say $_,$H[$_] for 0..$#H' input.tsv

 Remove column named 'height'.
 'height'という列を削除する。
 % perltab -pe 'del "height"' heightWeight.tsv
 or,
 又は、
 % perltab -e 'say F( grep {!/^height$/} @H )' heightWeight.tsv 

=kw SNset|-g|--grep|grep
 Select rows from heightWeight.tsv matching ids listed in file 'ids.txt' (holding one id per line).
 heightWeight.tsvの内、IDリストにある行を抽出する。IDリストは、各行にひとつのIDという形式のファイルids.txtから読み込む。
=settab    ==
 % perltab -b 'SNset %S, "ids.txt"'  -ge '$S{ F(ID)}'  heightWeight.tsv   > selected.tsv
=unsettab

 After selecting those rows, sort them so that the ids are in the same order as in "/tmp/ids".
 ids.txtにあるIDの行を抽出した後、ids.txtと同じ順番に並べ替える。
 % perltab -b 'SNset %S, "ids.txt"' -s '$S{ F(ID) }' selected.tsv
=wk SNset|-g|--grep|grep

=kw bigger
  Print rows with weight > 60.
  weightが60より大きい行を表示する。
  % perltab -gd 'bigger F(weight), 60'
=wk bigger

=kw lesser
  Print rows with weight < 60.
  weightが60より小さい行を表示する。
  % perltab -gd 'lesser F(weight), 60'
=wk lesser

=kw bemin
 Print minimum value of column 'banana', quietly ignoring non-numerical values.
 列'banana'の最小値を出力する。非数字の値(欠損値など)は静かに無視される。
 % perltab -e 'bemin $m, F(banana)' -z 'say $m'
=wk bemin

=kw beminmax|-p|-e2
 Linearly normalize numbers in column 'banana' to be in range [0,1].
 列'banana'にある数値が[0,1]の範囲に入るように線形正規化を行う。
=settab    ==
 % perltab -p -e 'beminmax $min, $max, F(banana)'   -e2 'donum  sub{ $_= ($_-$min) / ($max-$min)}, F(banana)'   fruit.tsv
=unsettab
=wk beminmax|-p|-e2

=kw -H|header|label|grep
 Print column labels containing 'eight'.
 'eight'を含む列ラベルを出力する。
 % perltab  -H 'say  grep /eight/, @H'
=wk -H|header|label|grep

=kw allnum|-g|--grep|grep
 Remove any data rows with non-numerical entries.
 数値でない値を持つ行を取り除く。
 % perltab -gd 'allnum @F'
=wk allnum|-g|--grep|grep

=kw num0
 Average all columns in a file (Non-numeric data treated as 0).
 各列の平均値を出力する(非数値データはゼロとして扱われる)。
=settab     ==
 % perltab  -d '$s[$_]+= num0 $F[$_] for 0..$#F'  -z 'say map {$_/(NR-1)} @s'
=unsettab
=wk num0

=kw -s2|--sort2
 Sort heightWeight.tsv so that IDs found in file age.tsv come first, in the same order as they appear in age.tsv.
 age.tsvとheightWeight.tsvの間の共通IDがage.tsvの中と同じ順番でheightWeight.tsvの先頭に来るようにheightWeight.tsvのデータ行を並べ替える。
=settab    ==
 % perltab -e '$R{ F(ID)}= NR'  -s201U '$R{ F(ID)}'   age.tsv  -in2 heightWeight.tsv
=unsettab

 Same as above but with the block of common ids at the end.
 上記の例と同様であるが、共通IDをage.tsvを先頭ではなく、最後に移す。
 age.tsvとheightWeight.tsvの間の共通IDがage.tsvの中と同じ順番でheightWeight.tsvの最後に来るようにheightWeight.tsvのデータ行を並べ替える。
=settab    ==
 % perltab -e '$R{ F(ID)}= NR'  -s2U01 '$R{ F(ID)}'   age.tsv  -in2 heightWeight.tsv
=unsettab

 Same as above but sorted in place, i.e. sort the common IDs to mirror their order in age.tsv, but do not move them as a block to the top or bottom.
 上記の例と同様に共通のIDをage.tsvと同じ順番に並べ直すが、先頭や最後に移さない。つまり、共通しないIDの行の位置は変わらない。
=settab    ==
 % perltab -e '$R{ F(ID)}= NR' -s201 '$R{ F(ID)}'  age.tsv  -in2 heightWeight.tsv
=unsettab

 or one can use the default sort spec 'AB01NM',
 又はソート指定のデフォルト値'AB01NM'も利用できる。
=settab    ==
 % perltab -e '$R{ F(ID)}= NR'  -s2 '$R{ F(ID)}'  age.tsv -in2 heightWeight.tsv
=unsettab
=wk -s2|--sort2

=kw insaft
 Add a Japanese Heisei fiscal year column 'FY', with entries like "H27" to a file with YYYYMMDD format column 'Date'.
 YYYYMMDD形式の入力ファイルに平成年度の列'FY'を追加する。
=settab    ==
 % perltab -pe 'insaft qw(Date FY), sub{ ($y,$m)= unpack "a4a2", F(Date);  "H". (($m>3)+ $y-1989)}   YYYYMMDD.tsv
=unsettab
=wk insaft

=kw -z|--end|transpose|転置
 Transpose rows and columns.  'chop' removes trailing tab characters.
 行と列の転置。chopは行末のタブ文字を削除する。
=settab    ==
 % perltab -e '$T[$_] .= "$F[$_]\t" for 0..$#F'   -z 'chop, say  for @T'  input.tsv
=unsettab
=wk -z|--end|transpose|転置

=kw NR
 Prepend row number column to each row.
 各行の先頭に行番号の列を入れる。
 % perltab -e 'say NR,@F'  input.tsv
=wk NR

=kw -p|--print-each-row
 Lexically sort rows by label, except for the first.  Comment lines are also output by -p.
 最初の行を除き、行をラベルをABC順に並べ替える。-pではコメント行もそのまま表示される。
 % perltab -pe '@F= F($H[0], sort @H[1..$#H])'  fruit.tsv
=wk -p|--print-each-row

 Similar to the above, but skipping comment lines.
 同上にラベルを並べ替えるが、コメント行は表示されない。
 % perltab -e 'say F($H[0], sort @H[1..$#H])'  fruit.tsv

=kw donum|-g2|--grep2|grep
 Output the rows with the maximum height value among rows in two files.
 ふたつの入力ファイルにあるデータから身長(height値)が最大の行を表示する。
=settab    ==
 % perltab -d 'bemax $m, F(hei)' -gd2 'donum {$m==$_} F(hei)'  heightWeight.tsv more_heightWeight.tsv  -in2 heightWeight.tsv more_heightWeight.tsv
=unsettab

 or if there are no missing values, more simply:
 もし欠損値がなければ、このコマンドでも良い、
=settab    ==
 % perltab -d 'bemax $m, F(hei)' -gd2 '$m==F(hei)'  heightWeight.tsv more_heightWeight.tsv  -in2 heightWeight.tsv more_heightWeight.tsv
=unsettab
=wk donum|-g2|--grep2|grep

=kw qtiles|xqtiles|quantile|median|quartile|percentile|分位値|分位数|中央値|４分位値|四分位値|４分位数|四分位数|百分位値|パーセンタイル
=head2  Examples with statistical functions|統計関数の使用例
=wk qtiles|xqtiles|quantile|median|quartile|percentile|分位値|分位数|中央値|４分位値|四分位値|４分位数|四分位数|百分位値|パーセンタイル
=kw statistics|統計
=kw stats|mean|stdev|median|quartile|平均値|標準偏差|中央値|４分位値|四分位値|４分位数|四分位数
 Print a few descriptive stats on feature avgT...
 特徴量avgT…についていくつかの代表値を表示する。
 % perltab -d 'push @d, F(avgT)'  -z 'say stats @d'  input.tsv
=wk stats|mean|stdev|median|quartile|平均値|標準偏差|中央値|４分位値|四分位値|４分位数|四分位数

=kw qtiles|xqtiles|quantile|median|quartile|percentile|分位値|分位数|中央値|４分位値|四分位値|４分位数|四分位数|百分位値|パーセンタイル
 Print quintiles of feature givenM...  Four values are output.
 特徴量givenM…の５分位値を表示する。出力は４つの数値となる。
 % perltab  -d 'push @d, F(givenM)'  -z 'say qtiles(5,@d)'  input.tsv

 Print quintiles of feature givenM...  With xqtiles the minimum and maximum values are included to produce 6 values.
 特徴量givenM…の５分位値を表示する。xqtilesを使うと、出力は最小値と最大値を含める６つの数値となる。
 % perltab  -d 'push @d, F(givenM)'  -z 'say xqtiles(5,@d)'  input.tsv
=wk qtiles|xqtiles|quantile|median|quartile|percentile|分位値|分位数|中央値|４分位値|四分位値|４分位数|四分位数|百分位値|パーセンタイル
 
=wk statistics|統計


=head1  SORTING ROWS|行の並べ替え
=kw -s|--sort|sort|ソート|整列|並べ替え
 perltab can be used to sort files.  The command:
 perltabで行を並べ替えることはできる。以下のコマンド、

 % perltab -s 'F(height)' input.tsv

 sorts the file input.tsv according to ascending order of height, with non-numerical values following (see details below).
 はinput.tsvを身長の昇順に並べ替える。数字以外の値は最後になる(詳細は以下)

 % perltab -s21 'F(height)' input.tsv

 Same as above but in descending order of height.
 上の例とほとんど同じであるが、身長は降順に並べ替えられる。

 In general,
 一般に、

 % perltab     -s[CMP1][,CMP2...] KEY_SCRIPT input.tsv
 % perltab --sort[CMP1][,CMP2...] KEY_SCRIPT input.tsv

 The commands above sorts the rows of input.tsv, by the row specific LIST of values returned by KEY_SCRIPT for each row.  The first element of LIST is the first key, which is to be sorted according to CMP1.  For rows with equal first keys, the 2nd LIST element is compared according to CMP2,...  as long as ties persist and more LIST elements are available.  If not enough CMPs are given, the last one is reused.
 以上のコマンドはinput.tsvの行をKEY_SCRIPTが行毎に返すリストLISTをキーとして並べ替える。まず、LISTの最初の要素をCMP1に従って比較し、同値であれば、LISTの２番目の要素をCMP2に従って比較し、同値がなくなるか、LISTの要素が尽きるまでそれを進めていく。CMP指定の数が足りない場合、最後のCMP指定を再利用する。

 CMP is a case insensitive string composed of 01|10 N M U AB|BA atoms in the order of precedence.
 CMPは 01|10 N M U AB|BA の五つの値のタイプを優先順に並べた文字列(大文字/小文字の区別はない)。
=settab  ==
 01      numerical  ascending.
 10      numerical descending.
 N       Not-a-Number strings, such as 'NaN' and '-nan'.
 M       the missing value mark.
 U       perl's undef value (unrelated to the string 'undef').
 AB      anything else in  ascending Unicode order.
 BA      anything else in descending Unicode order.
 01      数値の昇順
 10      数値の降順
 N       'NaN'や'-nan'のようなnot a number文字列。
 M       欠損値の印。
 U       Perlのundef (文字列'undef'とは関係ない)
 AB      その他の文字列のUnicode昇順
 BA      その他の文字列のUnicode降順
=unsettab
 When CMP is entirely omitted the default value of 'AB01NM' is used.
 CMPをまるごと省略すると'AB01NM'が使われる。

 When neither 'ab' nor 'ba' are given, they are given lowest priority.
 When 'M' is omitted, the missing value mark is treated like any other string, and sorted according to 'ab' or 'ba'.
 When 'N' is omitted, strings looking like not-a-number are treated like any other string, and sorted according to 'ab' or 'ba'.
 'ab'と'ba'のどちらも指定しないと、(省略されたUを除けば)その優先順位は最後となる。
 'M'を省くと、欠損値の印は他の文字列と区別せず、'ab'もしくは'ba'の指定に従って並べ替えられる。
 'N'を省くと、NaNは他の文字列と区別せず、'ab'もしくは'ba'の指定に従って並べ替えられる。

 When 'U' is omitted from CMP, rows for which KEY_SCRIPT returns undef are pegged, i.e. not moved by the sort.
 CMPが'U'を含まない場合、スクリプトKEY_SCRIPTがundefを返した行の位置は固定したまま、ソートの対象にならない。

 Sort a ditto mode file of cities first by country, then by descending population.
 同上形式で都市を格納したファイルを国名のabc順、そして同じ国の中で人口の降順で並べ替える。
 % perltab -D -sab,10 'F(coun pop)'  NEAsiaMegacities_ditto.tsv

 Similarly, but with the row for Tokyo pegged.
 Tokyoの行の固定したまま、他の行を並べ替える。
=settab    ==
 % perltab -D -sab,10 'F(city)=~/Tokyo/? undef : F(coun pop)'  NEAsiaMegacities_ditto.tsv
=unsettab
=wk -s|--sort|sort|ソート|整列|並べ替え


=head1  SCRIPT FILE|スクリプトファイル
=kw -f|--script-file|script-file
The script to be run by perltab may be given with option -f|--script-file PATHNAME, instead of inline script options such as -e|row-script.  For convenience, the -n|--print-script option can be used to convert inline commands into script file format.  For example, after:
オプション-f|script-fileでファイル名を指定すると、perltabの実行スクリプトはそのファイルから読み込まれる。-nでコマンド行に書き込まれたスクリプトをスクリプトファイル形式に変換できる。例えば、以下のコマンドを実行した後、
=kw donum|-g|--grep|grep
 % perltab -nge 'donum {$_>70} F(Weight)' > listHeavy.perltab

  listHeavy.perltab
  --------------------------------------------------
  #perltab -g

  ROWS{
    donum {$_>70} F(qw(Weight))
  }
  ==================================================

 % perltab -f listHeavy.perltab  heightWeight.tsv

Should produce the same output as:
でlistHeavy.perltabを読み込むコマンドは、

 % perltab -ge 'donum {$_>70} F(Weight)'  heightWeight.tsv

のようにスクリプトを全部コマンド行に書き込んだコマンドと同じ出力をするはず。両方ともWeight列の値が数値データで70を越える行だけを出力する。donumの役割は数値データ以外の行を静かに飛ばすであり、Weight列のすべての値が数値であれば、-ge 'F(Weight) > 70'でも良い。
Both output the rows for which column 'Weight' has a numerical value over 70.  The donum{} construct serving to silently skip non-numerical values; if all values in column Weight are numerical, -ge 'F(Weight) > 70' would work as well.
=wk donum|-g|--grep|grep

=kw BLOCK|BEGIN|DATA|ROW|END
=head2  CODE BLOCKS IN SCRIPT FILE|スクリプト・ファイルでのコード・ブロック

The correspondence between inline script command line options and named code blocks is as follows:
コマンド行に書き込む各種スクリプトとスクリプト・ファイルに於けるブロック名の対応は以下の通り、


  b |begin          BEGIN
  b2|begin2         BEGIN2
  d |data-script    DATA
  d2|data-script2   DATA2
  e |rows-script    ROWS
  e2|rows-script2   ROWS2
  z |end            END

At the time of this writing (20170114) the use of script files with perltab has been tried out by not tested.
現時点(20170114)でperltabのファイルからスクリプトを読み込む機能は試されてはいるが検証はされていない。
=wk BLOCK|BEGIN|DATA|ROW|END
=wk -f|--script-file|script-file


=head1  TROUBLESHOOTING|問題解決
=head2  No output?|出力がない？

Maybe you forgot to add the -p|print-each-row flag.
-p|print-each-rowオプションを指定し忘れた？

 % perltab -e 'movbef qw(dog cat lion tiger)' animals.tsv

Silently moves the felines before dog, but prints nothing.
これは静かに猫科の連中を犬の前に並べ替えるが、何も表示はしない。

 % perltab -pe 'movbef qw(dog cat lion tiger)' animals.tsv

Prints the updated table.
こうすると更新した表は表示される。

=head2  Weird behavoir|変の挙動
When perltab is used like: %perltab -e 'SCRIPT', it is easy to accidently terminate the script prematurely.
perltabを% perltab -e 'SCRIPT'のように使う場合SCRIPT中の引用符がSCRIPTを途中で切ってしまうことがあります。
For example:
例えば、
 % perltab -H 'say '@F is: ',@F'  input.tsv

Which bash parses as
はbashで以下のように区切られる。
 % perltab  '-H'  'say @F'  'is:'  ',@F'  'input.tsv'

In cases like this q{} can be convenient.
この場合、q{}で引用すると便利だったりする。
 % perltab -H 'say q{@F is: },@F' input.tsv

The flexibility of perl syntax can be confusing at times.
perlの柔軟な構文は便利だが解りにくかったりもする。
以下のコマンドで構文がどのように解析されているかが調べられる。
If you are not sure how perl is parsing your script try:
 % perl -MO::Deparse -e  'YOUR SCRIPT'


=head1  CHECKING RESULTS|結果の尤もらしさを確認
=head2  LOOK|目視確認
Open in a speadsheet program (e.g. localc) and check by eye.  Some programs prefer .csv as a filename suffix, even when the fields are tab separated.
まず、localcなどの表計算プログラムで結果を見てみると良いでしょう。
表計算プログラムの一部はタブ区切りのファイルでも.csvのファイル名拡張子を好む。

=head2  COLUMN COUNT|列数の確認
One can check if the number of columns is the same for each row with this command:
各行の列数が一致しているかは以下のコマンドで確認できる。

  % perl -nE 'say y/\t//' data.tsv  |  uniq

=head2  COLUMNS ONLY PERMUTED?|列の順番以外に変更がない？
Given a script:
もし、

  % perltab  -e 'SCRIPT'  input.tsv  >  output.tsv

which only permutes characters within rows, the following two commands should give the same output.
というスクリプトがあり、そのスクリプトが（他の変更をせずに）行内の文字を並べ変えるだけなら、以下のふたつのコマンドの結果は一致するはず。

  % perl  -ne 'print sort split'  input.tsv  |  cksum

  % perl  -ne 'print sort split' output.tsv  |  cksum


=head1  PREREQUISITES|依存モジュール

perltab uses the following modules:
perltabは以下のモジュールを利用する。

  List::Util qw(first max maxstr min minstr reduce shuffle sum);
  List::MoreUtils qw(:all);
  POSIX qw(ceil floor);
  Scalar::Util qw(looks_like_number dualvar);
  Scalar::Util::Numeric qw(isfloat isint isneg);
  Hash::Util qw(lock_keys lock_keys_plus unlock_keys lock_hash);
  Statistics::Basic;
  Statistics::Distributions;
  Statistics::RankCorrelation;
  Unicode::Collate;
  Data::Lock qw(dlock dunlock);
  Text::Balanced


=head1  AUTHOR|著作者

The perltab program and this documentation was written by Paul Horton.
perltabプログラム及びこの説明文の著者はPaul Horton。


=head1  LICENSE|使用条件

Copyright 2016 Paul B. Horton <>.  perltab may be used, modified or redistributed under the GNU Public License GPLv3.  For possible use under others licenses, please contact the author.
著作権 Paul B. Horton 2016。perltabの利用、改良及び再配布はGNU Public License GPLv3の条件に合う範囲内なら可能である。他のライセンス下での使用の可能性については事前に著者にご連絡してください。


=head1  SEE ALSO|参照

fastapl, rename, Text::CSV

=head1  END_DOC
