## Supplementary material for "A Simulation Study of mRNA Diffusion and Mitochondrial Localization": Document source and data files: makefile

```
# Paul Horton 2017, 2019

.SUFFIXES:

.PRECIOUS: %.png

.PHONY: clean all


all: MLRfollowupSimulationManuscript_bioRxiv.pdf


DRAWINGS:= $(wildcard drawings/*.odg)

IMAGE_FILES:= $(DRAWINGS:.odg=.png)

$(IMAGE_FILES) : %.png : %.odg
	libreoffice --convert-to png --outdir drawings $<   #May silently fail if libreoffice is already running


%.pdf : %.tex %.bcf $(IMAGE_FILES)
	biber -E 'UTF-8' -U --bblsafechars $*
	lualatex --halt-on-error --shell-escape $(TEXFLAGS) $*
	@if grep 'Please rerun LaTeX' $*.log; then lualatex --halt-on-error --shell-escape $*; fi


%.bcf : %.tex
	lualatex --halt-on-error --shell-escape $*


clean :
	@rm -rf gnuplottex
	@rm -f *.gnuploterrors
	@rm -f drawings/*.png
	@rm -f *.aux
	@rm -f *.bcf
	@rm -f *.bbl
	@rm -f *.blg
	@rm -f *.log
	@rm -f *.run.xml
```
