## Supplementary material for "A Simulation Study of mRNA Diffusion and Mitochondrial Localization": Document source and data files: MLRfollowupSimulationManuscript.bib

@Online{PIH19B,
  longKey    = {PoulsenTM_HortonP_Hallmarks_BioRxiv2019},
  author     = {Thomas M. Poulsen  and  Kenichiro Imai  and  Martin C. Frith  and  Paul Horton},
  title      = {{Hallmarks of slow translation initiation revealed in mitochondrially localizing mRNA sequences}},
  year       = {2019},
  eprint     = {614255},
  eprinttype = {bioRxiv},
}


@Article{SDP13,
  longKey   = {ShahP_PlotkinJB_Cell2013_Rate},
  author    = {Premal Shah  and  Yang Ding  and  Malwina Niemczyk  and  Grzegorz Kudla  and  Joshua B. Plotkin},
  title     = {Rate-limiting steps in yeast protein translation},
  journal   = {Cell},
  year      = {2013},
  volume    = {153},
  number    = {7},
  pages     = {1589-1601},
  numPages  = {13},
  pmid      = {23791185},
  abstract  = {Deep sequencing now provides detailed snapshots of ribosome occupancy on mRNAs.  We leverage these data to parameterize a computational model of translation, keeping track of every ribosome, tRNA, and mRNA molecule in a yeast cell.  We determine the parameter regimes in which fast initiation or high codon bias in a transgene increases protein yield and infer the initiation rates of endogenous Saccharomyces cerevisiae genes, which vary by several orders of magnitude and correlate with 5’ mRNA folding energies.  Our model recapitulates the previously reported 5’-to-3’ ramp of decreasing ribosome densities, (in contrast to IngoliaNT_WeissmanJS_Science2009_Genome) our analysis shows that this ramp is caused by rapid initiation of short genes rather than slow codons at the start of transcripts.  We conclude that protein production in healthy yeast cells is typically limited by the availability of free ribosomes, whereas protein production under periods of stress can sometimes be rescued by reducing initiation or elongation rates.}
}

@Article{SVJ03,
  longKey   = {SylvestreJ_JacqC_GenomBio2003_Long},
  author    = {Julien Sylvestre  and  Stéphane Vialette  and  Marisol Corral-Debrinski  and  Claude Jacq},
  title     = {Long {mRNAs} coding for yeast mitochondrial proteins of prokaryotic origin preferentially localize to the vicinity of mitochondria},
  journal   = {Genome Biol.},
  year      = {2003},
  volume    = {4},
  number    = {7},
  pages     = {R44},
  numPages  = {9},
  pmid      = {12844360},
  PHSummary = {Use microarrays to measure the localization of mRNAs to the periphery of the mitochondria.  Introduce a measure called MFI in which 3 chips are used, one to measure the mitochondrial (and E.R.) membrane associated mRNA abundance, another to measure the abundance in the cytosolic fraction and a third to measure total mRNA expression.  They claim this measure is more reliable than MLR, but the evidence they gave for that was not completely clear to me.  They note that mRNA of genes of prokaryotic origin (more precisely with detectable homologs in the bacteria world) have a tendency to localize to the mitochondria, as do mRNA's of longer proteins.  In fact MFI is roughly proportional to the log of the protein length.  Their MLR values include both mitochondrial and E.R. membranes (and they are very up front about this and in their Fig. 2, they show that on average mRNAs encoding for E.R. resident proteins have higher MLR values than those encoding mitochonrial proteins)},
  abstract  = {BACKGROUND: Subcellular messenger RNA localization is important in most eukaryotic cells, even in unicellular organisms like yeast for which this process has been underestimated. Microarrays are rarely used to study subcellular mRNA localization at whole-genome level, but can be adapted to that purpose.  This work focuses on studying the repartition of yeast nuclear transcripts encoding mitochondrial proteins between free cytosolic polysomes and polysomes bound to the mitochondrial outer membrane.  RESULTS: Combining biochemical fractionations with oligonucleotide array analyses permits clustering of genes on the basis of the subcellular sites of their mRNA translation.  A large fraction of yeast nuclear transcripts known to encode mitochondrial proteins is found in mitochondrial outer-membrane-bound fractions.  These results confirm and extend a previous analysis conducted with partial genomic microarrays.  Interesting statistical relations among mRNA localization, gene origin and mRNA lengths were found: longer and older mRNAs are more prone to be localized to the vicinity of mitochondria.  These observations are included in a refined model of mitochondrial protein import.  CONCLUSIONS: Mitochondrial biogenesis requires concerted expression of the many genes whose products make up the organelle.  In the absence of any clear transcriptional program, coordinated mRNA localization could be an important element of the time-course of organelle construction.  We have built a 'MitoChip' localization database from our results which allows us to identify interesting genes whose mRNA localization might be essential for mitochondrial biogenesis in most eukaryotic cells.  Moreover, many components of the experimental and data-analysis strategy implemented here are of general relevance in global transcription studies.}
}


@Article{SZ10,
  longKey   = {SiwiakM_ZielenkiewiczP_PLoSCompBio2010},
  author    = {Marlena Siwiak and Piotr Zielenkiewicz},
  title     = {A comprehensive, quantitative, and genome-wide model of translation},
  journal   = {PLoS Comput. Biol.},
  year      = {2010},
  volume    = {6},
  number    = {7},
  pages     = {1-15},
  numPages  = {15},
  pmid      = {20686685},
}

@Article{WJW14,
  longKey   = {WilliamsCC_WeissmanJS_Science2014_Targeting},
  author    = {Christopher C. Williams  and  Calvin H. Jan  and  Jonathan S. Weissman},
  title     = {Targeting and plasticity of mitochondrial proteins revealed by proximity-specific ribosome profiling},
  journal   = {Science},
  year      = {2014},
  volume    = {7},
  number    = {346},
  pages     = {748--751},
  numPages  = {4},
  pmid      = {25378625},
  abstract  = {Used proximity-specific ribosome profiling to comprehensively measure translation at the mitochondrial surface in yeast.  Most inner-membrane proteins were cotranslationally targeted to mitochondria, reminiscent of proteins entering the endoplasmic reticulum (ER).  Comparison between mitochondrial and ER localization demonstrated that the vast majority of proteins were targeted to a specific organelle.  A prominent exception was the fumarate reductase Osm1, known to reside in mitochondria.  We identified a conserved ER isoform of Osm1, which contributes to the oxidative protein-folding capacity of the organelle.  This dual localization was enabled by alternative translation initiation sites encoding distinct targeting signals.  These findings highlight the in vivo specificity of organellar targeting mechanisms.},
  PHnote    = {Provide full data tables for download. Mention in passing that most mitochondrially localized mRNA are for proteins with 180 codons or more, but do not further pursue that observation}
}
